# Warmer temperatures result in maladaptive learning of sexual preferences

**DOI:** 10.1101/2020.11.05.369561

**Authors:** Marie-Jeanne Holveck, Doriane Muller, Bertanne Visser, Arthur Timmermans, Lidwine Colonval, Fabrice Jan, Michel Crucifix, Caroline M. Nieberding

**Affiliations:** Evolutionary Ecology and Genetics group, Earth and Life Institute, UCLouvain, Louvain-la-Neuve, Belgium; Evolution and Ecophysiology group, Department of Functional and Evolutionary Entomology, University of Liège - Gembloux Agro-Bio Tech, Passage des Déportés 2, 5030 Gembloux, Belgium; Earth and Climate, Earth and Life Institute, UCLouvain, Louvain-la-Neuve, Belgium

**Keywords:** Reproductive fitness, Climate warming, Seasonal polyphenism, Simulations, Learning

## Abstract

1. The impact of learning ability and sexual selection on the climate and biodiversity crisis are currently unclear.
2. Using the African butterfly *Bicyclus anynana*, which shows strong phenotypic plasticity (i.e., polyphenism) in response to temperature, we tested whether learning affects mate preferences under climate warming.
3. We first modelled climate warming scenarios and then showed experimentally that as temperature becomes an unreliable cue to the onset of the dry season, adult butterflies displayed the wet season rather than the dry season form.
4. Experienced females that were exposed to different male seasonal phenotypes during sexual maturation changed sexual preferences.
5. Female fertilization success was reversed for naive compared to experienced females, likely reducing female fitness following climate warming.
6. Our results emphasize the importance of sexual selection, learning, and their fitness consequences for understanding (mal)adaptation of natural populations to climate warming.

## Introduction

Climate warming is expected to lead to a global extinction rate of 15% by 2100 (Cahill et al., 2013; Urban, 2015). This predicted extinction rate should be taken with caution, however, because estimation models do not incorporate key biological responses of species (Urban, 2015), such as learning ability (Candolin & Wong, 2012a). Learning, which is a modification of behaviour resulting from experience with the environment, has emerged as a major response of species to human-induced rapid environmental change (see Table 1 in (Barrett, Zepeda, Pollack, Munson, & Sih, 2019) for empirical examples; (Nieberding, Marcantonio, Voda, Enriquez, & Visser, 2021)). Learning is important, because it can significantly change behaviours, for example aversive or reversal learning in response to danger or fear (Ozawa & Johansen, 2018; Rodrigues, Goodner, & Weiss, 2010; Tedjakumala & Giurfa, 2013) or biased learning (Westerman, Hodgins-Davis, Dinwiddie, & Monteiro, 2012). Compared to innate behaviour, learned behaviours may thus produce widely different predictions about species responses to human-induced rapid environmental change. Moreover, the most recent documented cases of learning in response to human-induced rapid environmental change have shown that learned behaviours are actually maladaptive (Barrett et al., 2019; Greggor, Trimmer, Barrett, & Sih, 2019). Maladaptation occurs whenever a “strategy” in a population becomes prevalent while it decreases the fitness of individuals (Crespi, 2000; Nesse, 2005). Learning may, therefore, accentuate the extinction risk of species rather than alleviate it (as is usually assumed (Thorpe, 1963)), leading to so-called evolutionary traps (i.e., adaptive traits become maladaptive (Robertson, Rehage, & Sih, 2013; Schlaepfer, Runge, & Sherman, 2002)).

**Table 1.**
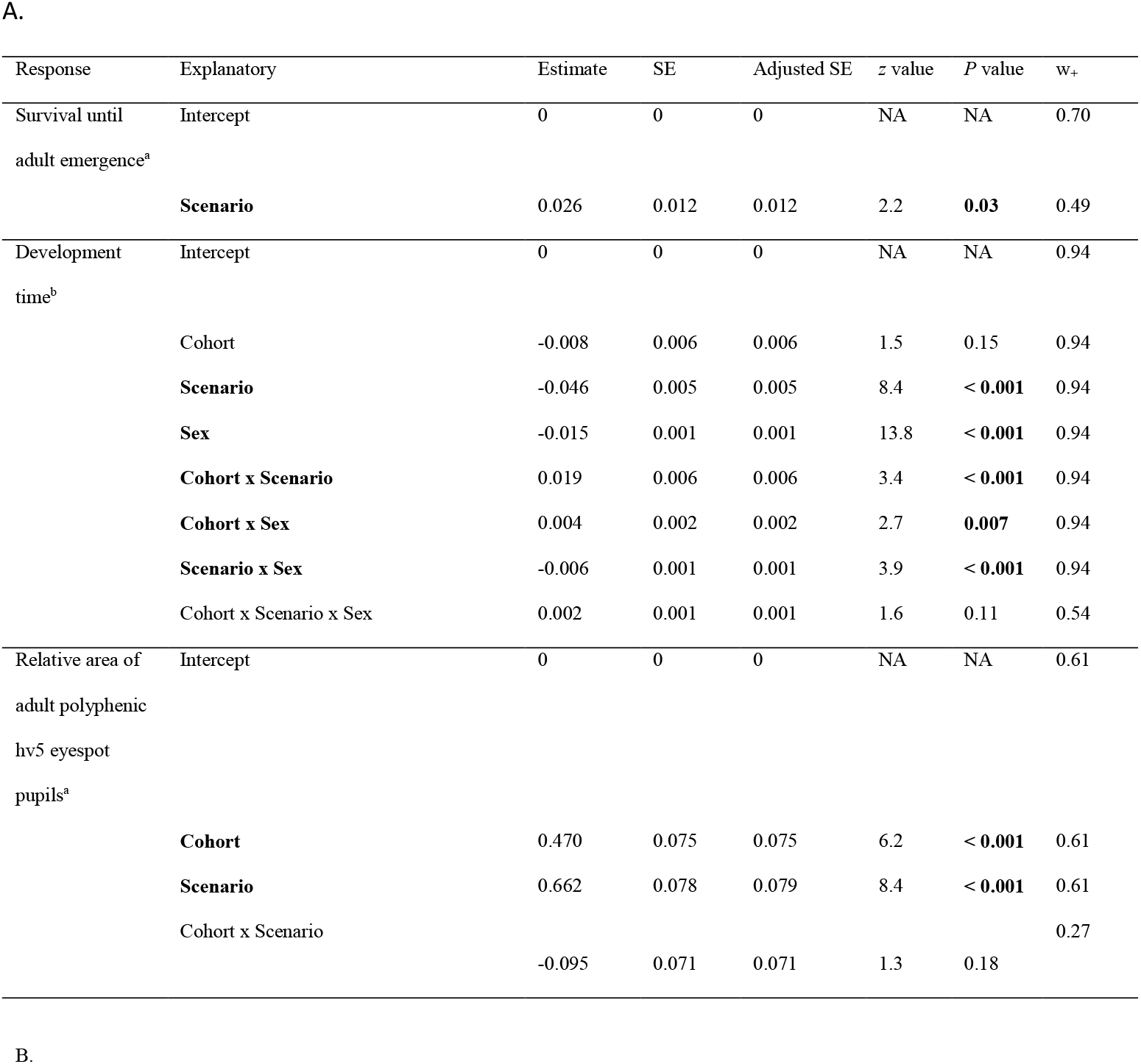
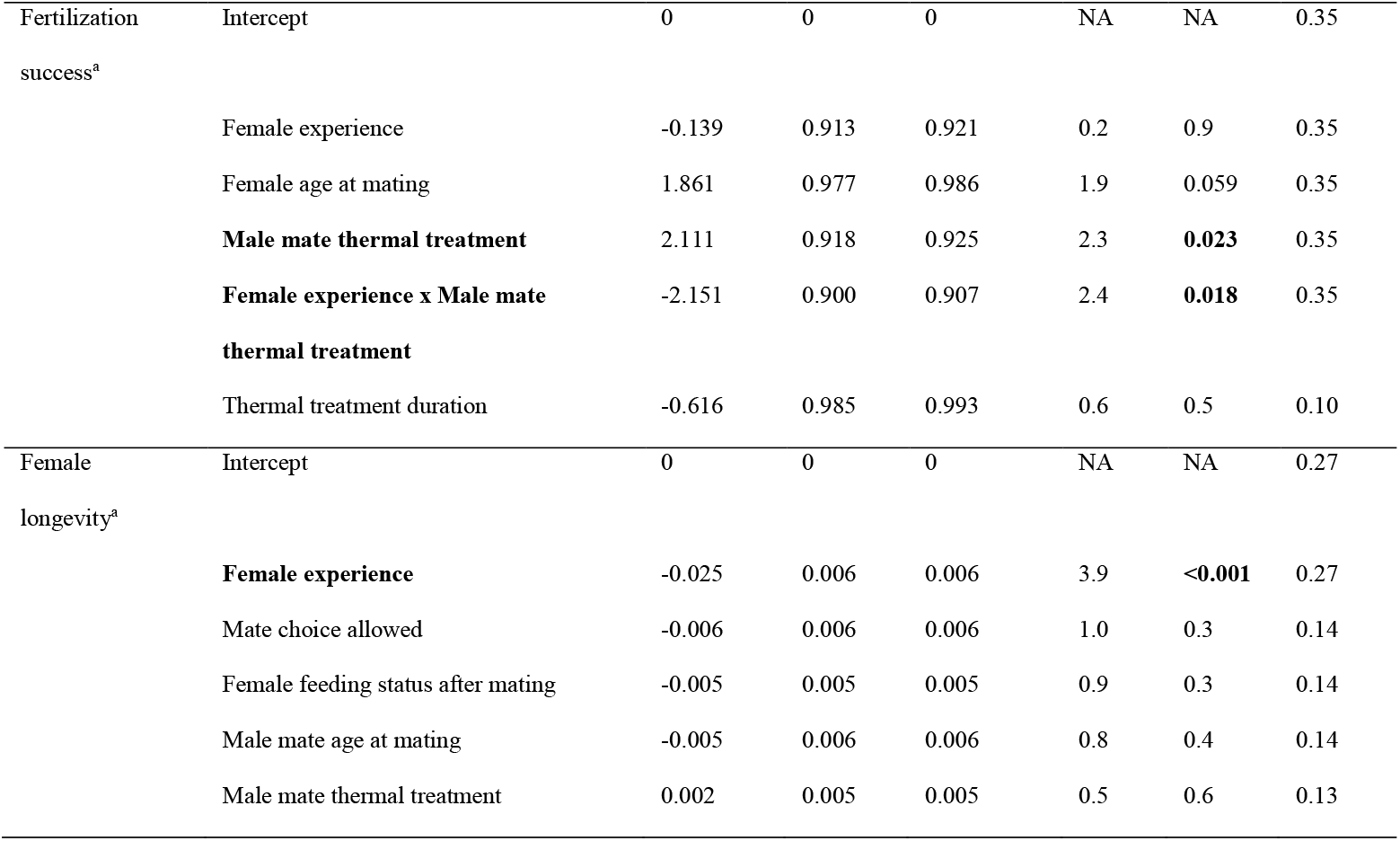
A) Statistical output for experiment 1 presented in Figure 3. B) Statistical output for experiment 2 presented in Figure 5. For both experiments, we used model-averaged standardized coefficients following conditional average of the set of supported candidate models for each response variable. For each conditional average of supported candidate models (Δ_*i*_ < 2), we provide the relative importance of explanatory variables (*P* ≤ 0.05 in bold). Of note, sexes differ in their developmental response to climate warming (Fig. 3b): males emerged earlier than females; female time to emergence (i.e., development time) decreased faster with warming (scenario-by-sex interaction) and increased faster throughout the seasonal transition (cohort-by-sex interaction) compared to males. These sex differences become smallest under the warmer scenario and at the beginning of the seasonal transition. Δ_*i*_ = difference in AIC^b^ (or AICc^a^) of model *i* with the best candidate model (i.e., the model with the smallest AIC^b^ or AICc^a^). SE = Conditional standard errors; Adjusted SE = Adjusted standard error estimator with improved coverage. w_*+*_ = sum of the normalized Akaike weights across all candidate models (Δ_*i*_ <2) in which the variable occurred. NA = not applicable.

| Response | Explanatory | Estimate | SE | Adjusted SE | z value | P value | $w_+$ |
| --- | --- | --- | --- | --- | --- | --- | --- |
| Survival until<br>adult emergence<sup>a</sup> | Intercept | 0 | 0 | 0 | NA | NA | 0.70 |
|  | **Scenario** | 0.026 | 0.012 | 0.012 | 2.2 | **0.03** | 0.49 |
| Development<br>time<sup>b</sup> | Intercept | 0 | 0 | 0 | NA | NA | 0.94 |
|  | Cohort | -0.008 | 0.006 | 0.006 | 1.5 | 0.15 | 0.94 |
|  | **Scenario** | -0.046 | 0.005 | 0.005 | 8.4 | **< 0.001** | 0.94 |
|  | **Sex** | -0.015 | 0.001 | 0.001 | 13.8 | **< 0.001** | 0.94 |
|  | **Cohort x Scenario** | 0.019 | 0.006 | 0.006 | 3.4 | **< 0.001** | 0.94 |
|  | **Cohort x Sex** | 0.004 | 0.002 | 0.002 | 2.7 | **0.007** | 0.94 |
|  | **Scenario x Sex** | -0.006 | 0.001 | 0.001 | 3.9 | **< 0.001** | 0.94 |
|  | Cohort x Scenario x Sex | 0.002 | 0.001 | 0.001 | 1.6 | 0.11 | 0.54 |
| Relative area of<br>adult polyphenic<br>hv5 eyespot<br>pupils<sup>a</sup> | Intercept | 0 | 0 | 0 | NA | NA | 0.61 |
|  | **Cohort** | 0.470 | 0.075 | 0.075 | 6.2 | **< 0.001** | 0.61 |
|  | **Scenario** | 0.662 | 0.078 | 0.079 | 8.4 | **< 0.001** | 0.61 |
|  | Cohort x Scenario |  |  |  |  |  | 0.27 |
|  |  | -0.095 | 0.071 | 0.071 | 1.3 | 0.18 |  |
| B. |  |  |  |  |  |  |  |
| Fertilization<br>success<sup>a</sup> | Intercept | 0 | 0 | 0 | NA | NA | 0.35 |
|  | Female experience | -0.139 | 0.913 | 0.921 | 0.2 | 0.9 | 0.35 |
|  | Female age at mating | 1.861 | 0.977 | 0.986 | 1.9 | 0.059 | 0.35 |
|  | **Male mate thermal treatment** | 2.111 | 0.918 | 0.925 | 2.3 | **0.023** | 0.35 |
|  | **Female experience x Male mate**<br>**thermal treatment** | -2.151 | 0.900 | 0.907 | 2.4 | **0.018** | 0.35 |
|  | Thermal treatment duration | -0.616 | 0.985 | 0.993 | 0.6 | 0.5 | 0.10 |
| Female<br>longevity<sup>a</sup> | Intercept | 0 | 0 | 0 | NA | NA | 0.27 |
|  | **Female experience** | -0.025 | 0.006 | 0.006 | 3.9 | **<0.001** | 0.27 |
|  | Mate choice allowed | -0.006 | 0.006 | 0.006 | 1.0 | 0.3 | 0.14 |
|  | Female feeding status after mating | -0.005 | 0.005 | 0.005 | 0.9 | 0.3 | 0.14 |
|  | Male mate age at mating | -0.005 | 0.006 | 0.006 | 0.8 | 0.4 | 0.14 |
|  | Male mate thermal treatment | 0.002 | 0.005 | 0.005 | 0.5 | 0.6 | 0.13 |

Most work on climate warming documents (mal)adaptation under natural selection, but another key biological mechanism to consider for improving the forecasting of species extinction rates is sexual selection. Over 50% of recorded climate change impacts are not due to direct effects of increased temperature, but to biotic interactions (Cahill et al., 2013). Sexual selection is an important biotic force of evolution (West-Eberhard, 2014) that is also affected by climate warming (Candolin & Heuschele, 2008; Candolin & Wong, 2012b; Grazer & Martin, 2012b). For example, a long-term study with the collard flycatcher *Ficedula albicollis* revealed that rising temperatures altered male ornamentation (i.e., reduced forehead patch size), reducing male mating success (Evans & Gustafsson, 2017). In the fly *Drosophila melanogaster*, lifespan and reproductive success decreased in response to an increase in temperature as a consequence of increased sexual conflict (García-Roa, Chirinos, & Carazo, 2019). Sexual selection may thus act against natural selection, increasing the risk of population extinction ((Kokko & Brooks, 2003; Martínez-Ruiz & Knell, 2017; Tanaka, 1996), but see (Svensson & Connallon, 2019)).

Sexual interactions are strongly affected by learning (in vertebrates: (Verzijden et al., 2012), and insects: (Dion, Monteiro, & Nieberding, 2019)), but whether and how animal responses to the climate crisis are mitigated by learned sexual behaviours remains virtually unexplored (Barrett et al., 2019; Greggor et al., 2019). We used a generalist insect as a model, the African butterfly *Bicyclus anynana* (Butler 1879; Nymphalidae), which has become one of the best-known examples of developmental plasticity in response to temperature. During the wet season, which lasts about ~5 months, temperature is high (Fig. 1) and there is an abundance of resources (Brakefield & Mazzotta, 1995; Brakefield, Pijpe, & Zwaan, 2007; Brakefield & Reitsma, 1991; Windig, Brakefield, Reitsma, & Wilson, 1994). High temperatures (> 23ºC; (Brakefield & Mazzotta, 1995)) lead to the development of a typical wet season male morph that is characterized by conspicuous eyespots to avoid predation (Chan, Rafi, & Monteiro, 2019; Kooi & Brakefield, 1999; Lyytinen, Brakefield, Lindstrfom, & Mappes, 2004). Several successive generations of wet season butterflies appear during the wet season. During the seasonal transition from wet to dry, temperatures are lower (between ~19ºC and ~23ºC; (Brakefield & Mazzotta, 1995)), leading to a higher frequency and development of an intermediate male phenotype (Fig. 1). During the dry season, there is reduced vegetation cover and an overall harsher, more stressful environment. A low temperature (<19ºC; (Brakefield & Mazzotta, 1995)) leads to development of the typical dry season male form, which is dull with reduced eyespots (Fig. 1). Only one generation spans the entire dry season, which lasts ~7 months.

**Fig. 1.**
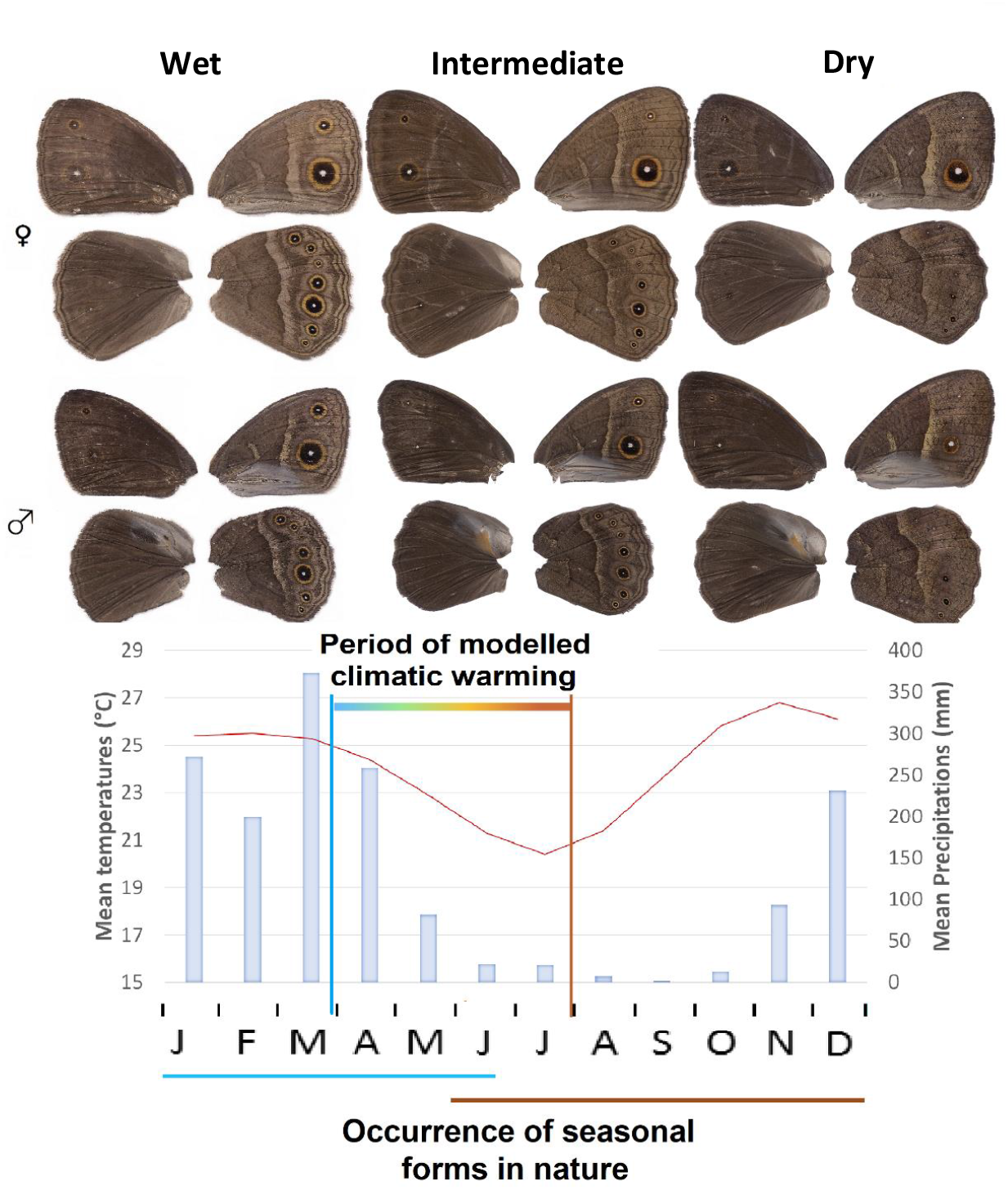
Morphological differences of females (top 2 wing pairs) and males (bottom 2 wing pairs) between dry, intermediate, and wet season phenotypes (Brakefield & Reitsma, 1991), as well as monthly average values of precipitation (blue bars) and temperature (red curve) in the field in Malawi, Africa (data available at (World-Bank-Group, 2019). The duration of the wet season is indicated by the blue line underneath the graph, while the duration of the dry season is represented by the orange line. The 120-day period of the climatic warming we modelled (Julian day 88 to 192, as in Fig. 3) is also depicted.

During the seasonal transition in nature, the three male phenotypes co-occur (Brakefield & Reitsma, 1991). Wet, intermediate, and dry season wing patterns are determined primarily by larval and pupal rearing temperature (Brakefield & Mazzotta, 1995). Both wet and dry season forms are adapted to the conditions of their respective seasons: wet season individuals favour reproduction, while dry season individuals have adapted to cope with limited food availability, while increasing longevity (Brakefield et al., 2007). Mating and mate preference have been studied predominantly in the wet season phenotype, where males show extensive courtship activity and females choose a mate based on visual and chemical cues (Muller et al., 2019; San Martin, Bacquet, & Nieberding, 2011). Previous work showed that all seasonal female forms prefer mating with the dry season male phenotype when offered a choice, increasing fitness (i.e., egg number) (Prudic, Jeon, Cao, & Monteiro, 2011). Another study revealed that a previous exposure to a male with more eyespots increases female preference for these artificially enhanced ornaments (Westerman, Chirathivat, Schyling, & Monteiro, 2014). *B. anynana* thus provides morphological evidence of adaptation to changing, seasonal climates that is easy to track (Muller et al., 2019), while mate preferences are under strong selection, particularly in changing environments.

Here, we combined climate modelling to test the effect of realistic climate warming scenarios on butterfly morphology, mating preference and fitness. We first modelled the seasonal transition as it will occur in 2100 in the native population of our African butterfly, located in Malawi. We then performed an experiment where we let individuals develop under the current and predicted climatic scenarios (experiment 1). Under the most stringent climate modelling scenario, the dry season temperature in 2100 will be comparable to the wet season temperature in 1988. We thus expected that the predicted increase in temperature under climate warming will lead to the production of wet season individuals at the beginning of the dry season. As a consequence, only maladapted male phenotypes, i.e., wet season, would be available for mating. To test how male seasonal form affects mate preferences, we then exposed wet season females to dry, wet or intermediate male phenotypes and measured female mate preference (experiment 2). We expected that a previous experience with any male phenotype (either wet, dry, or intermediate) would change mate preference towards that male phenotype, meaning that females learn to prefer mates that are actually present in the current environment. To estimate how female experience and male seasonal phenotype affect fitness, we subsequently measured the proportion of successfully hatched eggs, representing fertilization success, and female longevity (experiment 3). Our results revealed that, all else being equal, an increase in temperature will alter the appearance of seasonal forms (more wet season males at the onset of the dry season), reducing mating opportunities with the most adaptive and preferred seasonal form (i.e., dry season). We further showed that a previous experience of females with available males produces learned sexual preferences for intermediate season males that become maladaptive under climate warming by reducing the proportion of successfully hatched eggs and thus fitness.

## Methods

### Modelling climatic scenarios of the seasonal transition in 1988 and 2100

We modelled past and future natural, hourly, circadian temperature cycles for 104 consecutive days representing the transition from the wet to the dry season in Malawi (from March 30^th^ to July 12^th^). We modelled three climatic scenarios: the past climate in 1988, and two future scenarios, i.e., RCP4.5 and RCP8.5 (with a 3°C and 6°C increase in the region of interest, respectively). The photoperiod followed the light decrease throughout the seasonal transition with a 15-min monthly step. We encoded the three modelled climatic scenarios in three separate incubators. Given the difficulty of obtaining reliable, in-situ meteorological data, we constructed an idealized, smooth seasonal cycle of temperature by averaging 10 years of meteorological data during 1979-1989 [data from the Modern-Era Retrospective analysis for Research and Applications (MERRA), at latitude 12S and longitude 34E (Global-Modeling-and-Assimilation-Office-(GMAO)]. We then considered climate modulations available from the CMIP5 database at the same location (Taylor, Stouffer, & Meehl, 2012). Specifically, we computed the seasonal cycle of temperature anomalies (period 2080-2099 *versus* 1970-1990) of three models (CCSM4, HadGEM2-ES, and IPSL-CM5A-MR; at the time these data were downloaded, the HadGEM2-ES historical data were available until 1984 only), and added them to the reference scenario. We retained the HadGEM2-ES model, because its climate sensitivity is intermediate and considers two future climatic scenarios, i.e., RCP4.5 (about 3°C increase in the region of interest) and RCP8.5 (6°C increase)(IPCC, Pachauri, & Meyer, 2014), in addition to reference climatology for 1979-1989.

### Insect rearing

Experiments were performed with a laboratory population of *Bicyclus anynana* that was originally established from 80 gravid females collected in Malawi in 1988 (Brakefield, Beldade, & Zwaan, 2009). Between 400 and 600 butterflies per generation were used to maintain high levels of heterozygosity and polymorphism (Van’t Hof et al., 2005) to avoid loss of genetic diversity in the laboratory population. Larvae were fed with maize, *Zea mays*, and adults with moist, organic banana, *Musa acuminata* (Brakefield et al., 2009). Insects were reared, kept, and tested in a large climate-controlled room (5.8 × 5.7 m and 2.27 m high) under a standard temperature regime of 27°C (± 1°C), a 12:12h light:dark cycle, and a high relative humidity of 70% (±5%) representing the wet season in the field (Brakefield et al., 2009). For experiments, eggs were repeatedly collected from the laboratory population. Eggs of all thermal treatments were first reared in the standard rearing environment at 27°C. The developmental temperature experienced from the 4^th^ larval instar onwards determines the expression of adult polyphenism (Bear & Monteiro, 2013; Brakefield & Reitsma, 1991; Kooi & Brakefield, 1999; Oostra et al., 2011a). Larvae were, therefore, subsequently placed in groups for the different thermal treatments from the 4^th^ larval instar until the second half of their pupal stage (Fig. 2). To do so, each group of 4^th^ instar larvae was placed on a single plant inside a sleeve that covers the plant and protects the developing larvae (n = 25-50/sleeve). Newly emerged adults were sexed and individually marked on the ventral forewing with an indelible felt-tip pen. Groups of the same sex, seasonal form, and age of at most 10 males or 20 females were placed in cylindrical netted cages (diameter: 30 cm; height: 38 cm), except when stated otherwise. All experiments with adult butterflies were performed at a temperature of 27°C, unless mentioned otherwise.

**Fig. 2:**
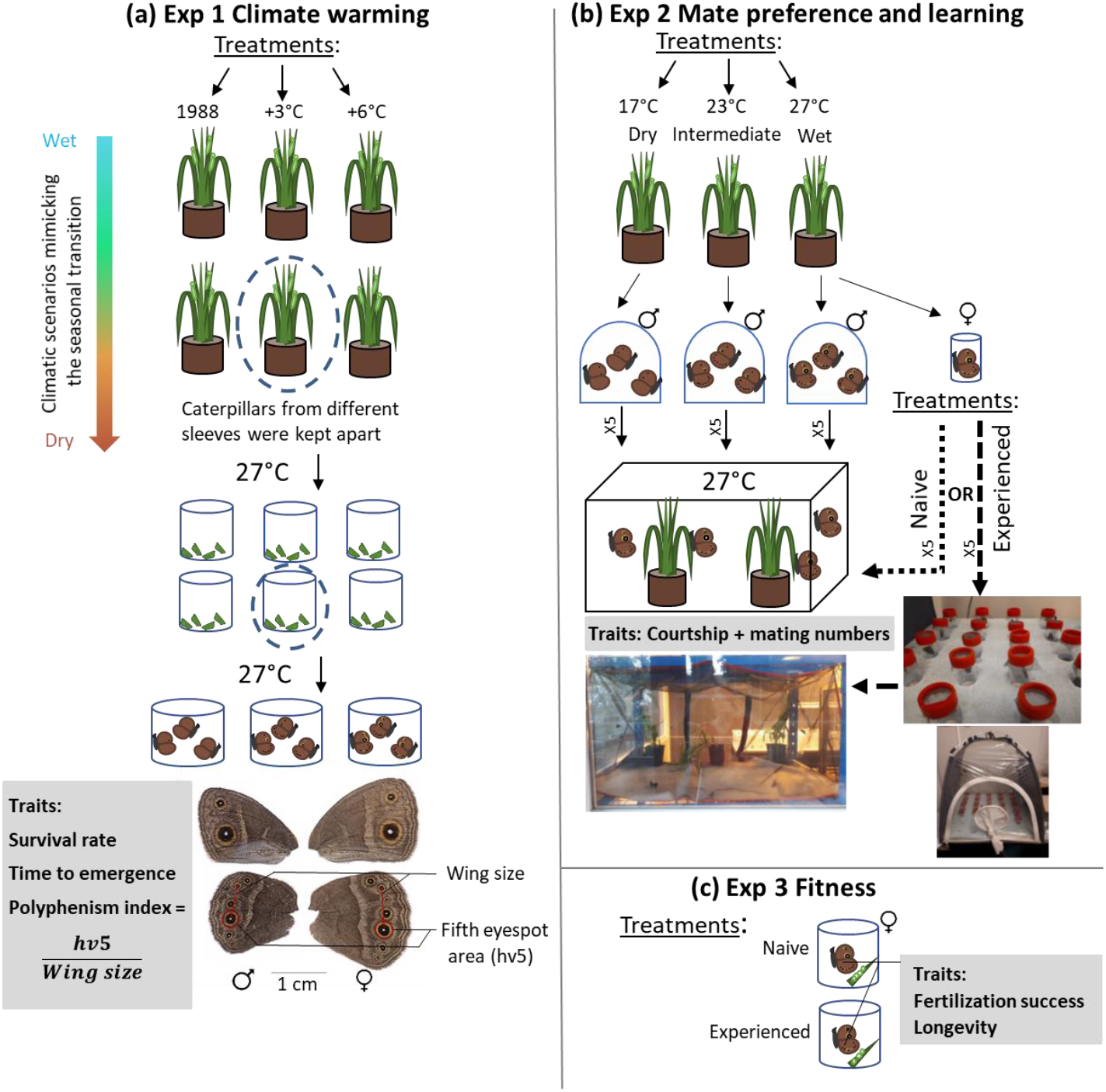
Experimental design for the climate warming experiment (a), the mate preference and learning experiment (b), and the fitness experiment (c), including the traits that were measured for each experiment. Prior to experiments, eggs were collected and kept at 27°C for 15 days; hence 15 day old larvae were used for subsequent experiments.

#### Experiment 1: Effects of climatic scenario on survival, time to first emergence, and seasonal phenotype

To test if climate warming would lead to the production of wet season individuals at the beginning of the dry season, we exposed butterflies to the modelled climatic scenarios. To cover the entire seasonal transition, we successively reared three cohorts of butterflies: the first 24 days for cohort 1, the following 28 days for cohort 2, and the final 35 to 50 days for cohort 3 (Fig. 1, Fig. 3). We evaluated the effects of increasing temperature on survival until adult emergence (i.e., the proportion of surviving individuals between the start of the thermal treatment, at the 4^th^ larval stage, and the first day of adult emergence), time to emergence as a proxy for development time (where day 1 is the date at which the first butterfly of a cohort emerged). Emerging adults were subsequently frozen at emergence at −80°C to measure adult polyphenism using the relative area of the fifth eyespot on the right ventral hindwing (i.e., hv5; Fig. 2). These traits were chosen, because they show the largest differences between seasonal forms (Muller et al., 2019).

**Fig. 3.**
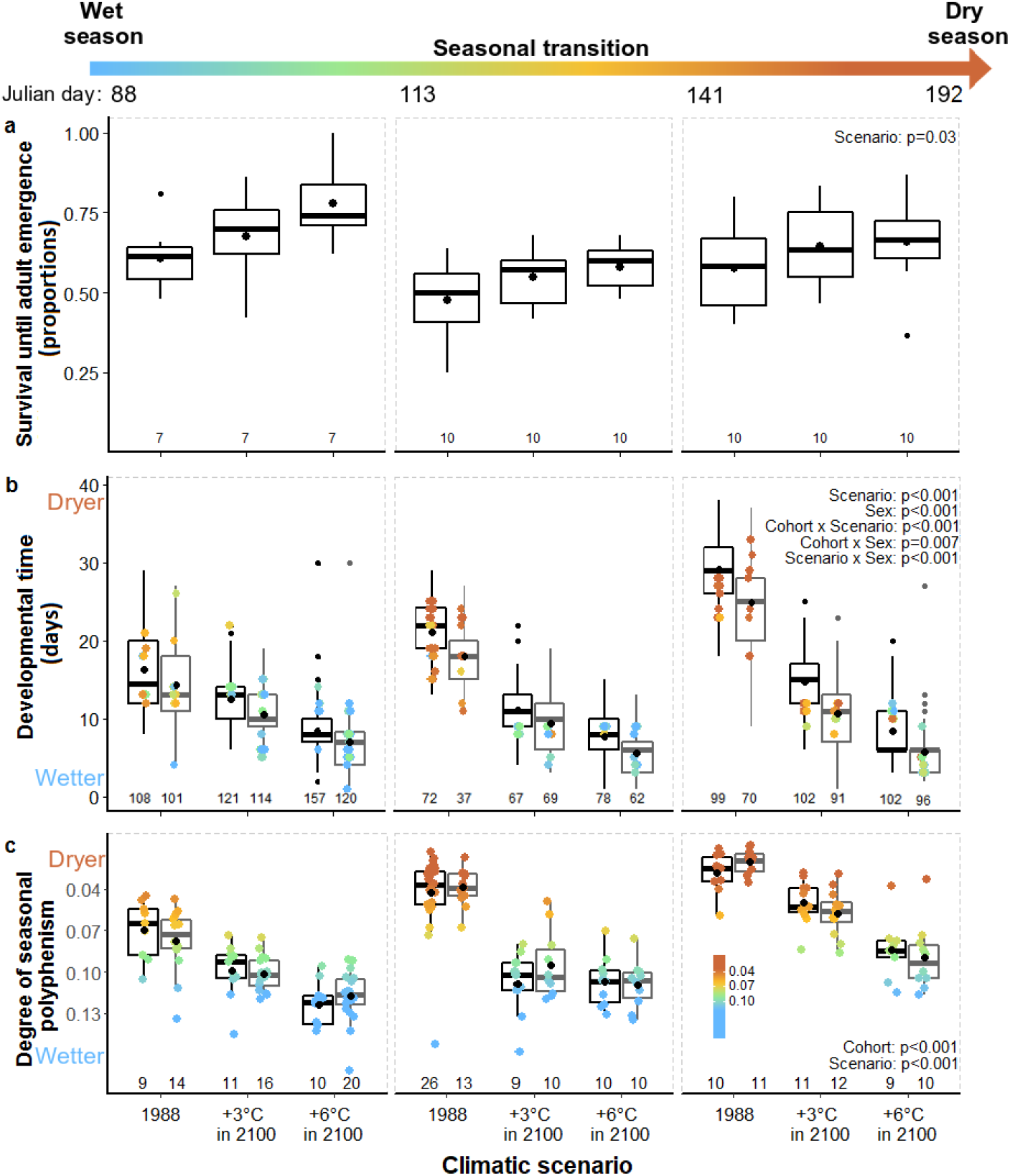
Climate warming during the wet to dry seasonal transition increases the proportion of surviving larvae until adult emergence (a), reduces the time to first emergence (i.e., development time); (b), and produces wet season butterflies at the onset of the dry season (based on the polyphenism index, as described in Fig. 1)(c). Statistics are presented in Table 1. The colour scale (in the bottom right graph) represents the change in polyphenism from extremely dry (red) to extremely wet (blue). Data are presented from left to right using three cohorts of larvae whose development cover the entire seasonal transition (cohort 1: from the 88^th^ day of the year to the 112^th^, cohort 2 from the 113^th^ to the 140^th^ day and cohort 3 from the 141^st^ to the 192^th^ day). For (b-c), colours indicate whether the individuals display the dry (brown), intermediate (yellow and green), or wet (blue) season forms, based on the relative eyespot area of females (black) and males (grey) (as in (Muller et al., 2019)). Mean values are indicated (black dot inside each boxplot) and samples sizes (below each boxplot) are the number of sleeves (n = 25-50 larvae/sleeve) in (a) and individuals in (b-c).

#### Experiment 2: Mate preference of naive and experienced females for different male seasonal forms

To test how the seasonal phenotype of males during the seasonal transition affects sexual selection, we produced the wet (temperature of 27°C), intermediate (23°C), and dry (17°C) seasonal forms (Brakefield et al., 2007; Brakefield & Reitsma, 1991; Muller et al., 2019; Oostra et al., 2011a; Prudic et al., 2011). We obtained experienced wet season females by exposing freshly emerged females to 8-16-day-old males of one seasonal form inside a flight cage (10 to 15 males per cage of 60 × 60 × 60 cm) for 3 hours (Fig. 2b). During exposure, females were kept separately in a small plastic cylindrical container (diameter 3.5 cm, height 7 cm, 60mL) with a custom-modified, wire mesh cap (7 × 7 mm), allowing the females to see, smell and touch the flying males above them, but preventing mating. This setup further prevented any visual and gustative contacts with females that were exposed in the flight case at the same time (with a maximum of 30 females spaced 5 cm apart). All females were fed with a slice of banana in their individual container. Males used for exposure were different than those used for subsequent mating experiments.

Mate preference experiments were performed with naive wet season females (20 replicates; control), as well as experienced wet season females exposed to wet (20 replicates), intermediate (17 replicates), or dry season males (15 replicates). For each replicate, 15 virgin males (i.e., 5 selected randomly from each of the three thermal treatments), 8 to 11 days of age, were left to compete in the presence of 5 virgin, wet season females, 3 to 6 days of age. Eight to 11-day-old virgin males were used, because all three components of the male sex pheromone are present at significant levels at this age, which plays a key role in mate choice decisions (Heuskin et al., 2014; Nieberding et al., 2008, 2012). We used 3-6-day-old virgin females, because females readily mate at this age in the laboratory (Brakefield et al., 2009; Nieberding et al., 2008, 2012). These ages also match the general age of females typically used in behavioural experiments with this species (Holveck, Gauthier, & Nieberding, 2015; Nieberding & Holveck, 2017). Before experiments started, we first introduced the males into the experimental area and left them to adapt and interact for 1 hour, after which females were added (Fig. 2b). The experimental area consisted of experimental cages (120 × 59 × 60 cm) containing two maize plants to increase perching and hiding opportunities and food. Using this larger cage size results in butterfly densities that are below the recommended threshold of 0.1 individual/dm^3^ for *B. anynana*. Together with a sex ratio of 3:1 males:females, this set-up creates a semi-natural condition that allows on the one hand the full expression of male courtship displays (including flight) and on the other hand female mate choice, where the female can reject a male by flying away (Holveck et al., 2015; Nieberding & Holveck, 2017, 2018). Each replicate was observed for 10 min every 30 min, with a total observation duration of 30 minutes per replicate. Direct behavioural observations for these 10 min were encoded using the software The Observer 5.0 (Noldus Information Technology, Wageningen, the Netherlands). When a male started courting a female, we followed the stereotyped courtship sequence until it ended before focusing on another male. Throughout each experiment, we noted the identity of all formed couples. As in many insects, *B. anynana* females do not engage in mating unless a male initiates courtship. We used mating proportions for analyses, representing the number of mated females out of the total number of tested females, i.e., observed mating proportions. These observed mating proportions were then corrected by male courtship number, i.e., theoretical mating proportions, because courtship activity differs among seasonal forms (Bear & Monteiro, 2013) and predicts mating success (Holveck et al., 2015; Nieberding & Holveck, 2017, 2018).

#### Experiment 3: Fitness-related trait measurements of wet season females mated with different male seasonal forms

To test the fitness consequences of a previous experience in females, we measured fertilization success as the proportion of successfully hatched eggs laid by either naive or experienced wet season females mated with males of different seasonal forms. We further measured female longevity. To do so, both naive and experienced females were placed singly in a 1-dm^3^ plastic box with wet cotton wool and a maize leaf for oviposition (Brakefield et al., 2009) after experiment 2 was finished. Feeding status differed between females, because part of the females obtained from the mate preference experiment (i.e., all 57 naive females, and 9 females exposed to wet season males) also received banana slices *ad libitum*. Leaves were replaced and egg fertilization success was recorded every three days. Longevity was recorded daily and calculated as the number of days from butterfly emergence to death.

### Statistical analyses

#### Experiment 1 Climate warming

We performed all statistics with Rstudio 1.0.143 (RStudio-Team, 2016). For all statistical models, we used model simplification based on the AIC values. We analysed the effect of the climatic scenarios in interaction with cohort on the survival using generalized linear mixed models GLMM (lme4 package in R (Bates, Maechler, Bolker, & Walker, 2015) with a binomial distribution (logit link function) and bobyqa optimizer. Random factors were sleeve identity nested in cohort identity nested in scenario identity (see (Barr, Levy, Scheepers, & Tily, 2013) for a discussion on using factorial variables as fixed and random effect). The model formula was Survival ~ Scenario * Cohort + (random effects: Scenario/Cohort/SleeveID) (Script S1 and Dataset S1). We added the interaction with sex (categorical variable) to analyse fixed effects on development time (time to emergence) and on polyphenism index (with a Box-Cox transformation to achieve normality) using GLMM with a Poisson distribution, a bobyqa optimizer and an observation-level random effect to capture overdispersion (Elston, Moss, Boulinier, Arrowsmith, & Lambin, 2001; Holveck et al., 2015), and with a normal distribution, respectively. Random and nested factors were scenario(cohort(sleeve)). The model formulae were Development time ~ Scenario*Cohort*Sex + (random effects: Scenario/Cohort/SleeveID) (see Script S2 and Dataset S2) and Polyphenism index ~ Scenario*Cohort*Sex + (random effects: Scenario/Cohort/SleeveID) (Script S3 and Dataset S3), respectively.

#### Experiment 2 Mate preference and learning

We tested for homogeneity among replicates per experiment using a Pearson’s Chi-squared test (with simulated *P*-value based on 1^e+06^ replicates to obtain a correct approximation; all *P* > 0.05) before performing analyses on the pooled data set. We then tested differences between observed and theoretical mating proportions of the three male thermal treatments per experiment using Pearson’s Chi-squared tests of goodness-of-fit. We subsequently searched for the male phenotype(s) responsible for these differences by comparing observed to theoretical mating proportions for each of the three male thermal treatments per experiment and in paired comparisons between male phenotypes per experiment, applying Pearson’s Chi-squared tests (reporting *P*-values with and without Holm-Bonferroni correction for multiple comparisons)(Millot, 2009). We also performed Fisher’s Exact tests when the expected frequencies were below 5 (Script S4, which contains the raw data).

#### Experiment 3 Fitness

We analysed how female fertilization success (as a binary variable) varied in response to mating with each seasonal phenotype and female experience (naive or exposed to wet season males). We added the age at mating of both sexes and the thermal treatment duration as fixed factors and experimenter(experiment(replicate)) as random factors. We used a GLMM with a binomial distribution, a bobyqa optimizer and an observation-level random effect to capture overdispersion. The model formula was Egg fertilization success ~ Male seasonal phenotype * Female experience + Female age at mating + Male age at mating + Treatment duration + (random effects: Experimenter/Experiment/Replicate). Next, we ran a similar GLMM on female longevity with a Poisson distribution, but added female feeding status after mating in interaction with male seasonal phenotype, and whether or not females could choose among males of different seasonal forms as fixed factors. The model formula was Longevity ~ Male seasonal phenotype * Female experience + Male seasonal phenotype **\*** Feeding status + Mate choice for seasonal forms + Female age at mating + Male age at mating + Treatment duration + (random effects: Experimenter/Experiment/Replicate). In all these analyses (Script S5 and Dataset S3), we rescaled and centred the continuous explanatory variables on the mean.

## Results

### Modelling predicts that 1988 wet season temperatures become common during the dry season in 2100

We mimicked the entire wet to dry seasonal transition (120 days) as it occurs in Malawi (Fig. 1; (Brakefield et al., 2009)) by implementing hour-to-hour modelled changes in temperature and monthly changes in photoperiod. Overall, climatic modelling showed that the alternation of wet and dry seasons was maintained in 2100, with a wet season of 5 to 6 months. Under the warmest scenario, however, the average temperature of both seasons increased by a quantity comparable to the amplitude of the seasonal cycle (6°C). Consequently, the higher temperatures typical of the wet season in 1988 will be common during the dry season in 2100.

#### Experiment 1: Higher temperatures lead to a maladaptive increase in wet season forms at the onset of the dry season

Comparing the 1988 data to the +6°C climatic scenario, climate warming increased survival by 21% (Fig. 3a; Table 1). Climate warming further reduced the time to emergence (as a proxy for development time) by 50% at the beginning of the seasonal transition (from 15.4 to 7.8 days), and by 74% (from 27.4 to 7.2 days) at the end (Fig. 3b; Table 1). Regarding seasonal phenotypes, we found that climate warming increased the degree of polyphenism, with more individuals showing the wet season phenotype throughout the seasonal transition (Fig. 3c): Butterflies displayed an increasingly dry season phenotype throughout the seasonal transition under the current scenario (which can be expected as temperature decreases during the seasonal transition; Fig. 3c), but development under the +3°C and +6°C climatic scenarios produced a wet season phenotype at the end of the seasonal transition. Under the +6°C climatic scenario, the phenotype during the dry season is indistinguishable from the phenotype in the 1988 wet season, in line with our expectation (Fig. 2c).

#### Experiment 2: Experience with different male seasonal forms modifies female mate preference

Naive females preferred to mate with dry season males over intermediate or wet season males (Fig. 4a; Table 2): the mating proportions with dry season males were significantly higher, while the mating proportions with intermediate and wet season males were lower than expected. Exposure of virgin wet season females to males of different seasonal forms induced a long-term (3 to 6 days) suppression of innate female preference for dry season males (Fig. 4; Table 2), meaning that long-term memory formation was involved. Furthermore, exposure to different male seasonal forms increased female mate preference towards intermediate season males compared to naive females (Fig. 4; Table 2). Indeed, observed mating proportions with intermediate season males either matched (Fig. 4b, d) or exceeded (Fig. 4c) the theoretical mating proportions (corrected for courtship activity)(Fig. 4a). The preference of wet season females for intermediate season males was highest when wet season females were exposed to intermediate season males, as expected (Fig. 4c; Table 2). This means that wet season females learn to prefer mating with intermediate season males after a previous exposure to intermediate males. More complex learned mate preferences were observed when females were exposed to dry or wet season males (Fig. 4b, 4d).

**Table 2.**
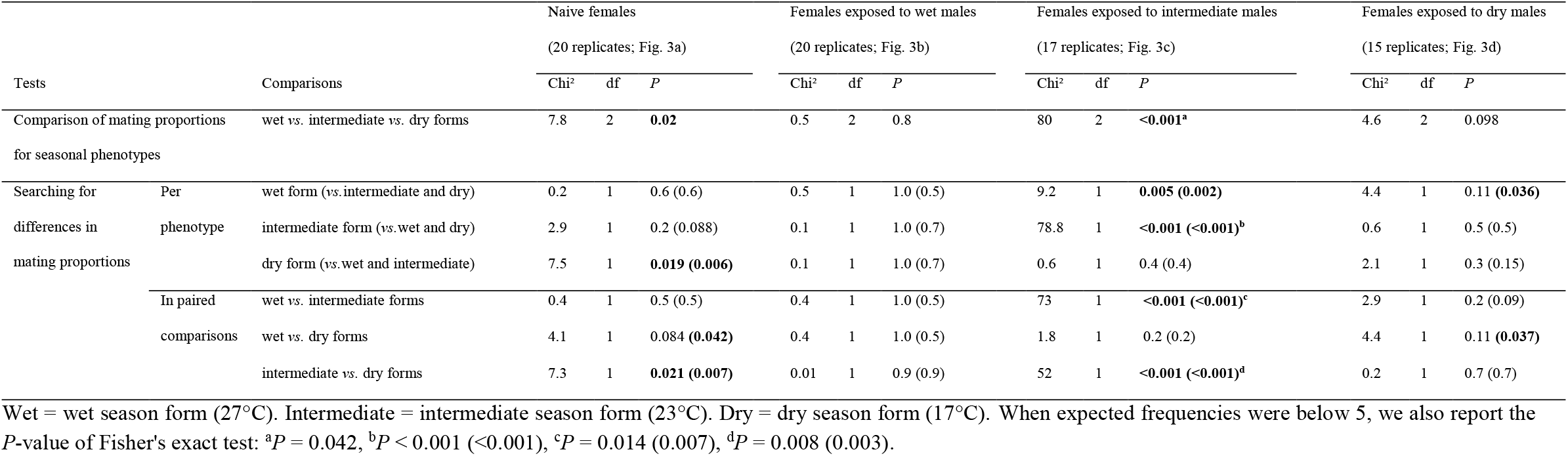
Statistical output of experiment 2 presented in Figure 4. Using Chi-squared tests for given probabilities, we first tested for differences in mating proportions with the different seasonal phenotypes; second, we searched for differences between observed and theoretical mating proportions for each of the three seasonal phenotypes, and in paired comparisons between seasonal phenotypes, by computing *P*-values with and without (in brackets) Holm-Bonferroni correction for multiple comparisons. *P* ≤ 0.05 are in bold.

**Fig. 4.**
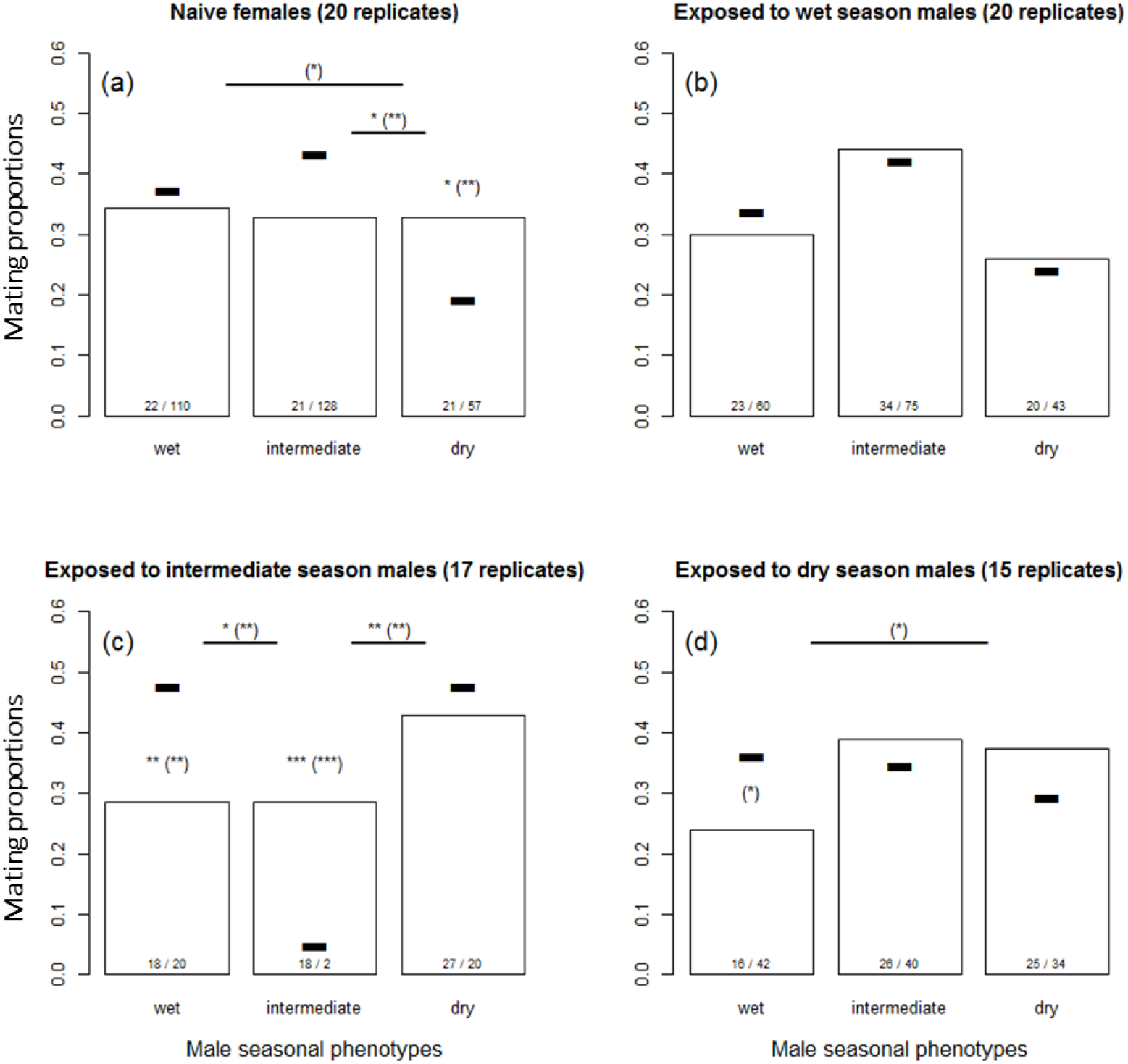
Female mate preference represented by observed (bars) and theoretical (black dashes) mating proportions. Significant differences between observed and theoretical mating proportions are indicated: *<0.5, **<0.01, ***<0.001 (using adjusted Holm-Bonferroni *P*-values and uncorrected *P*-values in brackets). Comparisons are made for each male phenotype and in paired comparisons between phenotypes. Samples sizes inside each bar are numbers of male matings/courtships. Statistics are presented in Table 2.

#### Experiment 3: Learning affects fertilization success and reduces longevity

Experienced females displayed reversed fertilization success with different male seasonal phenotypes compared to naive females (Fig. 5a and 5b). The proportion of successfully fertilized eggs of naive females was lower when females mated with dry rather than with intermediate or wet season males (Fig. 5a). Experienced females, however, had highest fertilization success when mating with dry season males (Fig. 5b). As expected, female experience and male seasonal phenotypes affect fertilization success. Furthermore, mating with wet season males is maladaptive for females following exposure to these males during sexual maturation. We further found that experienced females had reduced longevity (Fig. 5c), suggesting that learning generates some costs (Barrett et al., 2019).

**Fig. 5.**
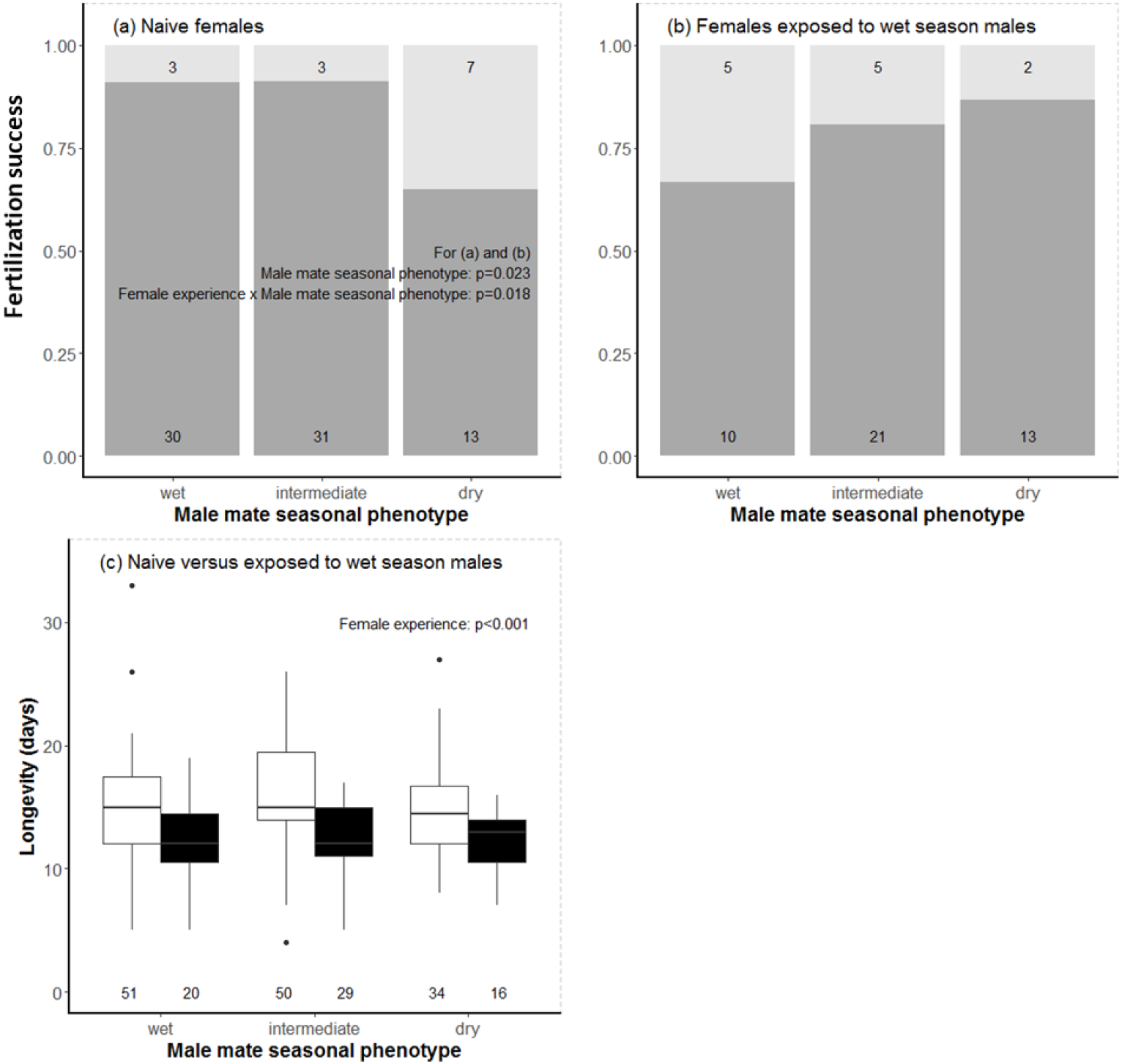
The proportion of wet season females whose eggs were fertilized (in dark grey) differ depending on whether the females are naive (a) or have been exposed previously to wet season males (b). The same was found for longevity, where naive and experienced females are depicted by white or black boxplots, respectively (c). Fertilization success, but not longevity, also depends on the seasonal form (wet, intermediate, dry) of males. Sample sizes are given inside each bar or below each boxplot.

## Discussion

This study provides, to the best of our knowledge, a first example of a learned sexual interaction that produces maladaptation under climate warming (Barrett et al., 2019; Greggor et al., 2019). As environmental cues become unreliable, in the case of *B. anynana* through temperature values that no longer predict the onset of the dry season, several maladaptive responses became apparent: *i)* production of the wet season form; *ii)* suppression of naive mate preference for dry season males after exposure to wet or intermediate season males; and *iii)* reduced fertilization success by experienced females mating with wet season males.

Our experimental results are relevant and important to consider for predicting the fitness consequences of sexual selection after the climate has warmed in nature. In the field, most virgin females are expected to get exposed to potential mates during the few days of sexual maturation following adult emergence, because this species is protandrous (i.e., males emerge before females), which we show induces sexual imprinting. Females remaining naive from any type of (social) exposure to available males until mating are likely rare in the field, also because this species is abundant. We thus expect that the reproductive fitness effect of experienced, rather than naive, females when mating with different male phenotypes better reflects what happens in the field. Overall, male availability, the thermal history of females, but also other factors, such as rainfall, can have major effects on fitness and survival of the population in nature.

The broader relevance of our results rests on two pillars. First, while learning is usually assumed to improve immediate adaptation to environmental variation (within the lifetime of an organism), the fitness effects of learning remain rarely assessed (Candolin & Wong, 2012b; Dion et al., 2019; Morand-Ferron, Cole, & Quinn, 2016; Nieberding, Van Dyck, & Chittka, 2018). Here, we discovered that learning reverses both mate preference and reproductive fitness. Our results thus reveal that using behavioural data from naive individuals to predict sexual responses of animals may lead to erroneous conclusions. We further found that learning reduced longevity. Whether learning increases or limits adaptation to sexual selection under climate warming remains largely undocumented so far (but see (Botero, Boogert, Vehrencamp, & Lovette, 2009) who found that more complex, learned, male bird songs were present in more unpredictable and unfavourable climatic environments). Maladaptive or suboptimal learning was also previously found for foraging (Avarguès-Weber, Lachlan, & Chittka, 2018), and two recent reviews document that learning in response to human-induced rapid environmental change is maladaptive for a larger number of environmental threats and behaviours (Barrett et al., 2019; Greggor et al., 2019). We suggest that maladaptive learning in sexual interactions under human-induced rapid environmental change may occur in many species, as most animals studied to date modify their sexual behaviours through learning (Dion et al., 2019; Nieberding et al., 2021; Verzijden et al., 2012), which could lead to evolutionary traps (Robertson et al., 2013).

Second, our results highlight the importance of sexual selection as a strong evolutionary force to include for quantifying species responses to human-induced rapid environmental change, such as climate warming (Candolin, 2019; Candolin & Heuschele, 2008). The effect of natural selection on species responses to the climate crisis has received considerable interest. Studies have shown that at the intraspecific level, climate warming leads to adaptive plastic and genetic changes in key phenotypic traits, including dispersal capacity and other morphological, physiological, behavioural, and life-history traits associated to phenology (Musolin & Saulich, 2012; Parmesan, 2006; Thackeray et al., 2016). However, the effects of anthropogenic environmental changes on sexual selection matter as well, because sexual selection may increase the extinction risk of populations (Holland, 2002; Martínez-Ruiz & Knell, 2017; Parrett & Knell, 2018; Plesnar-Bielak, Skrzynecka, Prokop, & Radwan, 2012). It is indeed increasingly acknowledged that human activities disturb important aspects of sexual selection (e.g., (Evans & Gustafsson, 2017; García-Roa et al., 2019)). This can be expected, as sexual selection has a strong environmental component (Ingleby, Hunt, & Hosken, 2010; Parrett & Knell, 2018). Regarding the climate crisis, sexual selection is altered through the availability of sexual partners (Twiss, Thomas, Poland, Graves, & Pomeroy, 2007) with potential effects on the proportion of extra-pair mating opportunities (Bichet, Allainé, Sauzet, & Cohas, 2016), relative fitness of mono-versus polyandrous females (Grazer & Martin, 2012a), assortative mating (Santos, Vieira, & Monteiro, 2018), sexual conflict (García-Roa et al., 2019), secondary sexual traits (Evans & Gustafsson, 2017), and mate preferences (Candolin, 2019; Fuxjäger et al., 2019).

Predicting the effects of maladaptive learned mating preference on the long-term maintenance of the butterfly population in the wild requires consideration of the effects of climate warming on the reduction in generation time and increased survival of the butterflies. Indeed, our climate modelling predicted that a true “dry season” will be maintained in 2100, the lack of dry season males could have consequences beyond reproductive outcomes, assuming that wet season males are incapable of surviving the dry season. Physiologically, dry season males are more resistant to starvation, have a lower resting metabolic rate and accumulate more fat compared to wet season males (Brakefield et al., 2007; Brakefield & Reitsma, 1991; Oostra et al., 2011b). Dry season males are predicted to become rare after climate warming, meaning that females will have to mate with wet season males, which reduces reproductive fitness. Mating with wet instead of dry season males may thus drastically reduce the proportion of females and developing eggs surviving the harsh, dry season until the next wet season by cascading effects of learned sexual interactions and fitness of reproducing females at the onset of the dry, harsh, tropical season. It remains difficult, however, to predict the fate of *B. anynana* populations in Malawi based on available information.

Our case study using the tropical butterfly *B. anynana* may be representative of a more general pattern, because butterflies are arthropods that represent most of the animal biomass and species diversity on Earth (Bar-On, Phillips, & Milo, 2018), and generally show strong phenotypic plasticity to cope with alternating seasons (Simpson, Sword, & Lo, 2011). Furthermore, tropical biomes both contain most biodiversity and are highly threatened by climate warming (Deutsch et al., 2008; Hoffmann, Chown, & Clusella-Trullas, 2013; Lister & Garcia, 2018; Sunday, Bates, & Dulvy, 2011), but see (Vasseur et al., 2014; Williams, Henry, & Sinclair, 2014). Understanding how maladaptations arise is increasingly important as humans rapidly transform the Earth into an environment where maladaptation is expected to become prevalent for most species (Crespi, 2000; Robertson et al., 2013). Our results suggest that global warming can have additive effects reducing, more than has been considered so far, the long-term survival probabilities of various taxa once learning and sexual selection are included as important biological mechanisms of species responses (Bell, 2013; Bradshaw & McNeilly, 1991). Our results thus advocate for integrating sexual selection, learning, and their fitness consequences to better understand (mal)adaptation to climate warming.

## Supporting information

Datasets and R Scripts

## Acknowledgements

We would like to thank Philippe Vernon, Franjo Weissing, Magdalena Kozielska, Georges Lognay, and Mathilde Scheifler for careful and insightful comments on previous versions of this manuscript. We are further grateful to the anonymous reviewers for their constructive feedback on previous versions of this manuscript. This research was supported by a “FRIA” PhD grant to DM and a “Chargée de Recherches” postdoctoral grant to MJH provided by the Fonds National de Recherche Scientifique (FNRS) of Belgium. This work was further funded through UCLouvain and the Fédération Wallonie-Bruxelles (Grant ARC 17/22-086) to CN. BV was supported by the Fonds National de Recherche Scientifique.

