## Supplementary material for "Warmer temperatures result in maladaptive learning of sexual preferences": Datasets and R Scripts: Script S1 survival.R

```
#Dataset S1
setwd("C:\\Users\\mullerd\\Desktop\\Projet anynana\\Th?se\\exp?rience volet 2\\exp2b\\ms finalisation\\ms_learning_soumission1")
setwd("F:/20180711_PCtravail/travail/Postdoc UCL/ms/B anynana/Holveck Muller/data and analyses")
data3 <- read.table("DatasetS1.txt", header=TRUE, sep="\t",dec=",")

summary(data3)
str(data3)

RealNOlarvae<-data3$NOlarvae-data3$NOlarvaeexcluded
data4<-cbind(data3,RealNOlarvae)
summary(data4)
mean(data4[data4$C=="3",]$RealNOlarvae)

#the needed variables
data4OK<-data4[,c("SleeveID","NOlarvae","NOlarvaeexcluded","NOadults","Survival","Scenario","Cohort","RealNOlarvae")]
summary(data4OK)

hist(data4OK$Survival)

tapply(data4OK$Survival,data4OK$Scenario,mean)
tapply(data4OK$Survival,data4OK$Scenario,sd)

0.6613757-0.5472116#at  27.06 - at 20.19
0.6613757*100/0.5472116-100

#GLM
mod<-glm(cbind(NOadults,RealNOlarvae-NOadults)~scale(Scenario,scale=T,center=T)*scale(Cohort,scale=T,center=T), data=data4OK, binomial)
summary(mod)#OVERDISPERSION (Residual deviance > residual d.f.)
mod2<-glm(cbind(NOadults,RealNOlarvae-NOadults)~scale(Scenario,scale=T,center=T)*scale(Cohort,scale=T,center=T), data=data4OK, quasibinomial)
summary(mod2)#OVERDISPERSION (Residual deviance > residual d.f.)

#GLMM with individual-level variability (accounting for overdispersion) = adding an observation-level random effect will capture the overdispersion.
library(lme4)
mlogit<-glmer(cbind(NOadults,RealNOlarvae-NOadults)~scale(Scenario, center=T, scale=T)*scale(Cohort, center=T, scale=T) + (1|Scenario/Cohort/SleeveID), data=data4OK, binomial)
mcloglog<-glmer(cbind(NOadults,RealNOlarvae-NOadults)~scale(Scenario, center=T, scale=T)*scale(Cohort, center=T, scale=T) + (1|Scenario/Cohort/SleeveID), data=data4OK, binomial(link = "cloglog"))
mprobit<-glmer(cbind(NOadults,RealNOlarvae-NOadults)~scale(Scenario, center=T, scale=T)*scale(Cohort, center=T, scale=T) + (1|Scenario/Cohort/SleeveID), data=data4OK, binomial(link = "probit"))
mcauchit<-glmer(cbind(NOadults,RealNOlarvae-NOadults)~scale(Scenario, center=T, scale=T)*scale(Cohort, center=T, scale=T) + (1|Scenario/Cohort/SleeveID), data=data4OK, binomial(link = "cauchit"))

mlogit_Nelder<- update(mlogit, control = glmerControl(optimizer="Nelder_Mead"))
mlogit_bobyqa<- update(mlogit, control = glmerControl(optimizer="bobyqa"))
library(optimx)
mlogit_nlminb<- update(mlogit, control = glmerControl(optimizer = "optimx", calc.derivs = FALSE,
                                                      optCtrl = list(method = "nlminb", starttests = FALSE, kkt = FALSE)))

mcloglog_Nelder<- update(mcloglog, control = glmerControl(optimizer="Nelder_Mead"))
mcloglog_bobyqa<- update(mcloglog, control = glmerControl(optimizer="bobyqa"))
library(optimx)
mcloglog_nlminb<- update(mcloglog, control = glmerControl(optimizer = "optimx", calc.derivs = FALSE,
                                                          optCtrl = list(method = "nlminb", starttests = FALSE, kkt = FALSE)))

mprobit_Nelder<- update(mprobit, control = glmerControl(optimizer="Nelder_Mead"))
mprobit_bobyqa<- update(mprobit, control = glmerControl(optimizer="bobyqa"))
library(optimx)
mprobit_nlminb<- update(mprobit, control = glmerControl(optimizer = "optimx", calc.derivs = FALSE,
                                                        optCtrl = list(method = "nlminb", starttests = FALSE, kkt = FALSE)))

mcauchit_Nelder<- update(mcauchit, control = glmerControl(optimizer="Nelder_Mead"))
mcauchit_bobyqa<- update(mcauchit, control = glmerControl(optimizer="bobyqa"))
library(optimx)
mcauchit_nlminb<- update(mcauchit, control = glmerControl(optimizer = "optimx", calc.derivs = FALSE,
                                                          optCtrl = list(method = "nlminb", starttests = FALSE, kkt = FALSE)))


library(AICcmodavg)
AICc(mlogit)#the default is ok
AICc(mprobit)
AICc(mcauchit)
AICc(mcloglog)#avec le plus petit AICc
#I don’t think the C-Log-Log link is used much today 
#because there is not a nice relationship between the βs for counting “successes” and the βs for counting “failures”.


#GLMER dispersion
library(blmeco) 
dispersion_glmer(mlogit) #it should be 1 and it shouldn't be over 1.4
dispersion_glmer(mprobit) #it should be 1 and it shouldn't be over 1.4
dispersion_glmer(mcauchit) #it should be 1 and it shouldn't be over 1.4
dispersion_glmer(mcloglog) #it should be 1 and it shouldn't be over 1.4

mlogit<-glmer(cbind(NOadults,RealNOlarvae-NOadults)~scale(Scenario, center=T, scale=T)*scale(Cohort, center=T, scale=T) + (1|Scenario/Cohort/SleeveID), data=data4OK, binomial,na.action = "na.fail")
mlogit_bobyqa<- update(mlogit, control = glmerControl(optimizer="bobyqa"))
summary(mlogit_bobyqa) 

#VIF
library(car)
mod2<-glm(cbind(NOadults,RealNOlarvae-NOadults)~scale(Scenario,scale=T,center=T)+scale(Cohort,scale=T,center=T), data=data4OK, quasibinomial)
vif(mod2)

#information theory model selection
#AICc should be used instead AIC when sample size is small in comparison to the number of estimated
#parameters (Burnham & Anderson 2002 recommend its use when n/K < 40); n=sample size; K=no of parameters.
81/7 

library(MuMIn)
mod3<-dredge(mlogit_bobyqa, rank = "AICc",
             beta = "sd")#coefficients standardized by SD
mod3
write.table(mod3, "F:/20180711_PCtravail/travail/Postdoc UCL/Doriane/ms2/data & stat/myAIC.txt",sep="\t") 
mod_average <- summary(model.avg(mod3, subset = delta < 2))
mod_average
capture.output(mod_average, file = "F:/20180711_PCtravail/travail/Postdoc UCL/Doriane/ms2/data & stat/mysummary.txt")

#BEST MODEL
m1best<-glmer(cbind(NOadults,RealNOlarvae-NOadults)~ scale(Scenario,center=T,scale=T) + (1|Scenario/Cohort/SleeveID),family=binomial, data=data4OK,na.action = "na.fail")
m1best_bobyqa<- update(m1best, control = glmerControl(optimizer="bobyqa"))
summary(m1best_bobyqa)
library(car)
Anova(m1best_bobyqa)#Analysis of Deviance Table (Type II tests)

#FULL MODEL
mlogit<-glmer(cbind(NOadults,RealNOlarvae-NOadults)~scale(Scenario, center=T, scale=T)*scale(Cohort, center=T, scale=T) + (1|Scenario/Cohort/SleeveID), data=data4OK, binomial,na.action = "na.fail")
mlogit_bobyqa<- update(mlogit, control = glmerControl(optimizer="bobyqa"))
summary(mlogit_bobyqa)
library(car)
Anova(mlogit_bobyqa)#Analysis of Deviance Table (Type II tests)

#GRAPH
library(ggplot2)
data4OK$Cplot <- ifelse(data4OK$Cohort=="0.9", "3", ifelse(data4OK$Cohort=="2.5", "2", "1"))
cohort.labs <- c("88 to 112 days", "113 to 140 days","141 to 190 days")
names(cohort.labs) <- c("1","2","3")

anno <- data.frame(x1 = c(2.87), y1 = c(0.99),  Cplot =c("3"), 
                   lab = c("Scenario: p=0.03"))
anno

sample<-data.frame(x1 = c(1,2,3,1,2,3,1,2,3), y1 = c(0.05,0.05,0.05,0.05,0.05,0.05,0.05,0.05,0.05),  Cplot =c("1","1","1","2","2","2","3","3","3"), 
                   lab = c("7", "7", "7","10","10","10","10","10","10"))
sample


p<-ggplot(data4OK, aes(x=as.factor(Scenario), y=Survival)) +
  geom_boxplot(color=c("black"),lwd=1.01)+ 
  theme(panel.background = element_blank(),axis.line.x = element_line(colour = "black",size=0.75), axis.line.y = element_line(colour = "black",size=0.75), axis.text.y=element_text(size=13), axis.text.x=element_blank(),axis.title.y=element_text(size=14,face="bold"), axis.title.x=element_blank())+
  facet_wrap( ~ Cplot, scales="free_x",labeller = labeller(Cplot = cohort.labs))+
  theme(strip.text.x = element_blank(),strip.background = element_rect(color="grey70", fill="white", size=1.5, linetype="solid"))+
  xlab("Climatic scenarii") + ylab("Survival until adult emergence\n(proportions)")+
  theme(legend.position="none",panel.border = element_rect(linetype = "dashed", fill = NA, color='grey80'),strip.background = element_blank())+
  scale_fill_manual(values=c("white","white"))+
  scale_x_discrete(labels=c("1988","+3?C \nin 2100","+6?C \nin 2100","1988","+3?C \nin 2100","+6?C \nin 2100","1988","+3?C \nin 2100","+6?C \nin 2100"))+
  geom_text(data = anno, aes(x = x1,  y = y1, label = lab),size=4)+
  geom_text(data = sample, aes(x = x1,  y = y1, label = lab),size=3)
p
p<-p+stat_summary(fun.y=mean, geom="point", shape=16, size=2.5,position =position_dodge(width=0.75))
p
```
