## Supplementary material for "Warmer temperatures result in maladaptive learning of sexual preferences": Datasets and R Scripts: Script S2 development.R

```
#Dataset S2
setwd("C:\\Users\\mullerd\\Desktop\\Projet anynana\\Th?se\\exp?rience volet 2\\exp2b\\ms finalisation\\ms_learning_soumission1")
setwd("F:/20180711_PCtravail/travail/Postdoc UCL/ms/B anynana/Holveck Muller/data and analyses")
data2 <- read.table("DatasetS2.txt", dec=",",sep="\t",header=TRUE)
summary(data2)
str(data2)

#the needed variables
dataOK<-data2[,c("ButterflyID","SleeveID","Sex","DevTime","Scenario","Cohort","Relativehv5")]
summary(dataOK)
str(dataOK)

hist(dataOK$DevTime)

#GLM
mod<-glm(DevTime~scale(Scenario,center=T,scale=T)*scale(Cohort,center=T,scale=T)*Sex,family=poisson,data=dataOK)
summary(mod) #OVERDISPERSION (Residual deviance > residual d.f.)
mod2<-glm(DevTime~scale(Scenario,center=T,scale=T)*scale(Cohort,center=T,scale=T)*Sex,family=quasipoisson,data=dataOK)
summary(mod2) #OVERDISPERSION (Residual deviance > residual d.f.)

#VIF
library(car)
vif(glm(formula = dataOK$DevTime ~ scale(dataOK$Cohort, center = TRUE, scale = TRUE) + 
          scale(dataOK$Scenario, center = TRUE, scale = TRUE) + dataOK$Sex),poisson)


#GLMM with individual-level variability (accounting for overdispersion) = adding an observation-level random effect will capture the overdispersion.
library(lme4)
m0<-glmer(DevTime~ scale(Scenario,center=T,scale=T)*scale(Cohort,center=T,scale=T)*Sex + (1|Scenario/Cohort/SleeveID),family=poisson, data=dataOK,na.action = "na.fail")
summary(m0)
m0_Nelder<- update(m0, control = glmerControl(optimizer="Nelder_Mead"))
m0_bobyqa<- update(m0, control = glmerControl(optimizer="bobyqa"))
summary(m0_bobyqa)

#GLMER dispersion
library(blmeco) 
dispersion_glmer(m0_bobyqa) #it should be 1 and it shouldn't be over 1.4

dataOK$obs <- 1:nrow(dataOK)
m1<-glmer(DevTime~ scale(Scenario,center=T,scale=T)*scale(Cohort,center=T,scale=T)*Sex + (1|Scenario/Cohort/SleeveID)+(1|dataOK$obs),family=poisson, data=dataOK,na.action = "na.fail")
summary(m1)
m1_Nelder<- update(m1, control = glmerControl(optimizer="Nelder_Mead"))
m1_bobyqa<- update(m1, control = glmerControl(optimizer="bobyqa"))
summary(m1_bobyqa)

#information theory model selection
#AICc should be used instead AIC when sample size is small in comparison to the number of estimated
#parameters (Burnham & Anderson 2002 recommend its use when n/K < 40); n=sample size; K=no of parameters.
1666/12 

library(MuMIn)
mod3<-dredge(m1_bobyqa, rank = "AIC",
             beta = "sd")#coefficients standardized by SD
mod3
write.table(mod3, "F:/20180711_PCtravail/travail/Postdoc UCL/Doriane/ms2/data & stat/myAIC.txt",sep="\t") 
mod_average <- summary(model.avg(mod3, subset = delta < 2))
mod_average
capture.output(mod_average, file = "F:/20180711_PCtravail/travail/Postdoc UCL/Doriane/ms2/data & stat/mysummary.txt")

#BEST MODEL
m1best<-glmer(DevTime~ scale(Scenario,center=T,scale=T)*scale(Cohort,center=T,scale=T)*Sex + (1|Scenario/Cohort/SleeveID)+(1|dataOK$obs),family=poisson, data=dataOK,na.action = "na.fail")
m1best_bobyqa<- update(m1best, control = glmerControl(optimizer="bobyqa"))
summary(m1best_bobyqa)
library(car)
Anova(m1best_bobyqa)#Analysis of Deviance Table (Type II tests)


#GRAPH
interaction.plot(dataOK$Cohort,dataOK$Sex,dataOK$DevTime)
is.factor(dataOK$Cohort)
interaction.plot(dataOK$Scenario,dataOK$Sex,dataOK$DevTime)
is.factor(dataOK$Scenario)


library(ggplot2)

library(reshape2)
dataOK$Cplot <- ifelse(dataOK$Cohort=="0.9", "3", ifelse(dataOK$Cohort=="2.5", "2", "1"))
cohort.labs <- c("88 to 112 days", "113 to 140 days","141 to 190 days")
names(cohort.labs) <- c("1","2","3")

anno <- data.frame(x1 = c(2.85,3.03,2.51,2.71,2.622), y1 = c(39,37,35,33,31),  Cplot =c("3","3","3","3","3"), 
                   lab = c("Scenario: p<0.001","Sex: p<0.001", "Cohort x Scenario: p<0.001", "Cohort x Sex: p=0.007","Scenario x Sex: p<0.001"))
anno
sample<-data.frame(x1 = c(0.8,1.2,1.8,2.2,2.8,3.2,0.8,1.2,1.8,2.2,2.8,3.2,0.8,1.2,1.8,2.2,2.8,3.2), y1 = c(0.05,0.05,0.05,0.05,0.05,0.05,0.05,0.05,0.05,0.05,0.05,0.05,0.05,0.05,0.05,0.05,0.05,0.05),  Cplot =c("1","1","1","1","1","1","2","2","2","2","2","2","3","3","3","3","3","3"), 
                  lab = c("108", "101", "121","114","157","120","72","37","67","69","78","62","99","70","102","91","102","96"))
sample

# attention les figures sont invers?es celles qui sont justes sont dans le ms
p1<-ggplot(dataOK, aes(x=as.factor(Scenario), y=DevTime)) +
  geom_boxplot(aes(fill=Sex),color=c("black","grey40","black","grey40","black","grey40","black","grey40","black","grey40","black","grey40","black","grey40","black","grey40","black","grey40"),lwd=1.01)+ 
  theme(panel.background = element_blank(),axis.line.x = element_line(colour = "black",size=0.75), axis.line.y = element_line(colour = "black",size=0.75), axis.text.y=element_text(size=13), axis.text.x=element_blank(),axis.title.y=element_text(size=14,face="bold"), axis.title.x=element_blank())+
  facet_wrap( ~ Cplot, scales="free_x")+
  theme(strip.text.x = element_text(size=14),strip.background = element_rect(color="grey70", fill="white", size=1.5, linetype="solid")) + ylab("Developmental time\n(days)")+
  theme(legend.position="none",panel.border = element_rect(linetype = "dashed", fill = NA, color='grey80'), strip.background = element_blank(),strip.text.y = element_blank(), strip.text.x = element_blank())+
  scale_fill_manual(values=c("white","white"))+
  scale_x_discrete(labels=c("1988","+3?C \nin 2100","+6?C \nin 2100","1988","+3?C \nin 2100","+6?C \nin 2100","1988","+3?C \nin 2100","+6?C \nin 2100"))+
  geom_point(aes(color=Relativehv5,fill=Sex),position = position_jitterdodge(jitter.width = 0.1, dodge.width = 0.7),shape=16, size=2.5)+
  scale_colour_gradientn(na.value = "NA", colours = c("sienna3","sienna3","goldenrod1","palegreen2","steelblue1","steelblue1","steelblue1"), breaks=c(0.04,0.07,0.10),name="" )+
  geom_text(data = sample, aes(x = x1,  y = y1, label = lab), size=3)+
  geom_text(data = anno, aes(x = x1,  y = y1, label = lab), size=4)
p1
#female in black; male in grey
p1<-p1+stat_summary(aes(fill=Sex),fun.y=mean, geom="point", shape=16, size=2,position =position_dodge(width=0.75))
p1

with(dataOK[dataOK$Sex=="f",], tapply(DevTime,list(Scenario,Cplot),length))
with(dataOK[dataOK$Sex=="m",], tapply(DevTime,list(Scenario,Cplot),length))
tapply(dataOK$DevTime,list(dataOK$Scenario,dataOK$Cohort),mean)
```
