## Supplementary material for "Warmer temperatures result in maladaptive learning of sexual preferences": Datasets and R Scripts: Script S3 polyphenism.R

```
#Dataset S2
setwd("C:\\Users\\mullerd\\Desktop\\Projet anynana\\Th?se\\exp?rience volet 2\\exp2b\\ms finalisation\\ms_learning_soumission1")
setwd("F:/20180711_PCtravail/travail/Postdoc UCL/ms/B anynana/Holveck Muller/data and analyses")
eyespot <- read.table("DatasetS2.txt", dec=",",header=TRUE, sep="\t")
summary(eyespot)
str(eyespot)

#the needed variables
eyespotOK<-na.omit(eyespot)
summary(eyespotOK)
str(eyespotOK)

hist(eyespotOK$Relativehv5)

eyespotOK$Relativehv5<-eyespotOK$hv5/(eyespotOK$Wing^2)

#GLMM with individual-level variability (accounting for overdispersion) = adding an observation-level random effect will capture the overdispersion.
library(lme4)
model3<-lmer(Relativehv5~scale(Cohort,center=TRUE, scale=TRUE)*scale(Scenario,center=TRUE, scale=TRUE)*Sex+(1|Scenario/Cohort/SleeveID), data=eyespotOK)
summary(model3)
shapiro.test(residuals(model3))
#transformation
BOXCOX
library(MASS)
bc<-boxcox(eyespotOK$Relativehv5~scale(eyespotOK$Cohort,center=TRUE, scale=TRUE)*scale(eyespotOK$Scenario,center=TRUE, scale=TRUE)*eyespotOK$Sex,lambda=seq(-8,8,0.01))
lambda<-bc$x[which.max(bc$y)]
lambda#--> similar to sqrt()
BoxCox transformation 
eyespotOK$bcL<-(((eyespotOK$Relativehv5^lambda)- 1)/lambda)
hist(eyespotOK$bcL)
hist(eyespotOK$Relativehv5)
#model
model4<-lmer(bcL~scale(Cohort,center=TRUE, scale=TRUE)*scale(Scenario,center=TRUE, scale=TRUE)*Sex + (1|Scenario/Cohort/SleeveID), data=eyespotOK,na.action = "na.fail",REML = FALSE)
summary(model4)
shapiro.test(residuals(model4)) 
library(car)
Anova(model4, type="II")#Analysis of Deviance Table (Type II tests)

#VIF
library(car)
vif(lm(formula = eyespotOK$bcL ~ scale(eyespotOK$Cohort, center = TRUE, scale = TRUE) + 
         scale(eyespotOK$Scenario, center = TRUE, scale = TRUE) + eyespotOK$Sex))

#information theory model selection
#AICc should be used instead AIC when sample size is small in comparison to the number of estimated
#parameters (Burnham & Anderson 2002 recommend its use when n/K < 40); n=sample size; K=no of parameters.
221/11

library(MuMIn)
mod3a<-dredge(model4, rank = "AICc",
              beta = "sd")#coefficients standardized by SD
mod3a
mod_averagea <- summary(model.avg(mod3a, subset = delta < 2))
mod_averagea 
#BEST MODEL
modelmin<-lmer(bcL~scale(Cohort,center=TRUE, scale=TRUE)+scale(Scenario,center=TRUE, scale=TRUE)+Sex + (1|Scenario/Cohort/SleeveID), data=eyespotOK)#t>2 suffit ? dire que c'est significatif
summary(modelmin)
library(car)
Anova(modelmin)#Analysis of Deviance Table (Type II tests)


#GRAPH TO FINISH!!!!
interaction.plot(eyespotOK$Cohort,eyespotOK$Sex,eyespotOK$Relativehv5)
is.factor(eyespotOK$Cohort)
interaction.plot(eyespotOK$Scenario,eyespotOK$Sex,eyespotOK$Relativehv5)
is.factor(eyespotOK$Scenario)


eyespotOK$Cplot <- ifelse(eyespotOK$Cohort=="0.9", "3", ifelse(eyespotOK$Cohort=="2.5", "2", "1"))
cohort.labs <- c("88 to 112 days", "113 to 140 days","141 to 190 days")
names(cohort.labs) <- c("1","2","3")


###### attention!! variable C degressive maintenant
library(ggplot2)
library(dplyr)

anno <- data.frame(x1 = c(2.85,2.93), y1 = c(0.17,0.158),  Cplot =c("3","3"), 
                   lab = c("Scenario: p<0.001","Cohort: p<0.001"))
anno
sample<-data.frame(x1 = c(0.8,1.2,1.8,2.2,2.8,3.2,0.8,1.2,1.8,2.2,2.8,3.2,0.8,1.3,1.8,2.2,2.8,3.2), y1 = c(0.175,0.175,0.175,0.175,0.175,0.175,0.175,0.175,0.175,0.175,0.175,0.175,0.175,0.175,0.175,0.175,0.175,0.175),  Cplot =c("1","1","1","1","1","1","2","2","2","2","2","2","3","3","3","3","3","3"), 
                   lab = c("9", "14", "11","16","10","20","26","13","9","10","10","10","10","11","11","12","9","10"))
sample
anno <- data.frame(x1 = c(2.85,2.93), y1 = c(0.165,0.153),  Cplot =c("3","3"), 
                   lab = c("Scenario: p<0.001","Cohort: p<0.001"))

p2<-ggplot(eyespotOK, aes(x=as.factor(Scenario), y=Relativehv5)) +
  geom_boxplot(aes(fill=Sex),color=c("black","grey40","black","grey40","black","grey40","black","grey40","black","grey40","black","grey40","black","grey40","black","grey40","black","grey40"),lwd=1.01,outlier.shape=NA)+ 
  theme(panel.background = element_blank(),axis.line.x = element_line(colour = "black",size=0.75), axis.line.y = element_line(colour = "black",size=0.75), axis.text=element_text(size=13),axis.title=element_text(size=14,face="bold"))+
  facet_grid( ~ Cplot, scales="free_x",labeller = labeller(Cplot = cohort.labs))+
  theme(strip.text.x = element_text(size=14),strip.background = element_rect(color="grey70", fill="white", size=1.5, linetype="solid"))+
  xlab("Climatic scenarii") + ylab("Degree of seasonal \npolyphenism")+
  theme(legend.position=c(0.7,0.5),legend.justification=c(0,0),panel.border = element_rect(linetype = "dashed", fill = NA, color='grey80'),strip.background = element_blank(),strip.text.y = element_blank(), strip.text.x = element_blank(), legend.title = element_text(size=10,face = "bold"),legend.text = element_text(size=9))+
  scale_fill_manual(values=c("white","white"))+
  scale_x_discrete(labels=c("1988","+3°C \nin 2100","+6°C \nin 2100","1988","+3°C \nin 2100","+6°C \nin 2100","1988","+3°C \nin 2100","+6°C \nin 2100"))+
  scale_y_continuous(breaks=c(0.04,0.07,0.10,0.13))+
  scale_y_reverse()+
  geom_point(aes(color=Relativehv5, fill=Sex),position = position_jitterdodge(jitter.width = 0.2, dodge.width = 0.7),shape=16, size=2.5)+
  scale_colour_gradientn(colours = c("sienna3","sienna3","goldenrod1","palegreen2","steelblue1","steelblue1","steelblue1"), breaks=c(0.04,0.07,0.10),name="", trans = "reverse")+
  guides(fill=FALSE)+
  theme(legend.key.size = unit(0.4, "cm"), legend.key.width = unit(0.3, "cm"),legend.title.align=0.5)+
  geom_text(data = anno, aes(x = x1,  y = y1, label = lab),size=4)+
  geom_text(data = sample, aes(x = x1,  y = y1, label = lab),size=3)  
p2
#female in black; male in grey
p2<-p2+stat_summary(aes(fill=Sex),fun.y=mean, geom="point", shape=16, size=2,position =position_dodge(width=0.75))
p2

library(egg) 
library(cowplot)
library(gridExtra)
require(grid) # for unit.pmax()
plotp1 <- ggplotGrob(p1) # convert to gtable
plotp2 <- ggplotGrob(p2) # convert to gtable
plotp <- ggplotGrob(p) # convert to gtable

p1.widths <- plotp1$widths[1:3] # extract the first three widths, 
# corresponding to left margin, y lab, and y axis
p2.widths <- plotp2$widths[1:3] # same for mpg plot
p.widths <- plotp$widths[1:3] # same for mpg plot

max.widths <- unit.pmax(p1.widths, p2.widths, p.widths) # calculate maximum widths
plotp1$widths[1:3] <- max.widths # assign max. widths to iris gtable
plotp2$widths[1:3] <- max.widths # assign max widths to mpg gtable
plotp$widths[1:3] <- max.widths # assign max widths to mpg gtable

# plot_grid() can work directly with gtables, so this works

plot_grid(plotp, plotp1, plotp2, labels = "auto",ncol=1, align = c("v"))


with(eyespotOK[eyespotOK$Sex=="f",], tapply(Relativehv5,list(Scenario,Cplot),length))
with(eyespotOK[eyespotOK$Sex=="m",], tapply(Relativehv5,list(Scenario,Cplot),length))
```
