## Supplementary material for "Warmer temperatures result in maladaptive learning of sexual preferences": Datasets and R Scripts: Script S4 mating success.R

```
setwd("F:/20180711_PCtravail/travail/Postdoc UCL/ms/B anynana/Holveck Muller/data and analyses")


PLAN
1. Test for heterogeneity among replicates
2. Chi2 on Mating probability
3. Chi2 on Mating probability per male form across experiments
4. Chi2 on Mating probability between exp1 and the other exps pairwise

###########
1. Test for heterogeneity among replicates
###########

#EXP1 (20 replicats) - naive females

#Test for heterogeneity among replicates using a Pearson's Chi-squared test 
#(or a Fisher's Exact test if the expected frequencies were close to 5 or less) before performing the analyses on the pooled data set

M <- as.table(rbind(c(1,0,2), c(1,2,0),c(1,1,2),c(1,1,2),
                    c(0,1,1),c(1,2,0),c(1,0,1),c(0,0,1),
                    c(1,1,1),c(1,1,0),c(1,0,1),c(0,1,2),
                    c(1,3,1),c(3,0,0),c(0,0,2),c(2,2,0),
                    c(3,2,0),c(3,1,1),c(1,1,2),c(0,2,2)))
dimnames(M) <- list(replicat = c("1", "2", "3","4", "5", "6","7", "8", "9","10","11", "12", "13","14", "15", "16","17", "18", "19","20"),
                    phenotypeM = c("WS","IS", "DS"))

(Xsq <- chisq.test(M))  # Prints test summary
chisq.test(M,simulate=TRUE,B=1000000)#use simulated p-values
fisher.test(M)
#if not significant = homogeneity = no heterogeneity among replicates = replicates are independant


#EXP2 (20 replicats) - females exposed to WS males

#Test for heterogeneity among replicates using a Pearson's Chi-squared test 
#(or a Fisher's Exact test if the expected frequencies were close to 5 or less) before performing the analyses on the pooled data set

M <- as.table(rbind(c(0,3,1), c(2,0,0),c(1,0,0),c(2,0,0),
                    c(1,2,1),c(1,2,1),c(1,1,2),c(1,2,1),
                    c(1,3,0),c(2,0,4),c(2,1,0),c(0,5,0),
                    c(1,2,1),c(0,3,1),c(1,3,0),c(0,1,1),
                    c(3,2,1),c(1,2,1),c(2,1,2),c(1,1,3)))
dimnames(M) <- list(replicat = c("1", "2", "3","4", "5", "6","7", "8", "9","10","11", "12", "13","14", "15", "16","17", "18", "19","20"),
                    phenotypeM = c("WS","IS", "DS"))

(Xsq <- chisq.test(M))  # Prints test summary
chisq.test(M,simulate=TRUE,B=1000000)#use simulated p-values
fisher.test(M)
#if not significant = homogeneity = no heterogeneity among replicates = replicates are independant


#EXP3 (17 replicats) - females exposed to IS23 males

#Test for heterogeneity among replicates using a Pearson's Chi-squared test 
#(or a Fisher's Exact test if the expected frequencies were close to 5 or less) before performing the analyses on the pooled data set

M <- as.table(rbind(c(0,0,2), c(1,2,1),c(2,0,1),c(0,1,1),
                    c(1,2,2),c(0,0,3),c(2,0,3),c(2,1,1),
                    c(1,0,2),c(1,1,3),c(2,1,1),c(2,1,0),
                    c(2,1,1),c(1,2,1),c(0,1,2),c(1,2,2),
                    c(0,3,1)))
dimnames(M) <- list(replicat = c("1", "2", "3","4", "5", "6","7", "8", "9","10","11", "12", "13","14", "15", "16","17"),
                    phenotypeM = c("WS","IS", "DS"))

(Xsq <- chisq.test(M))  # Prints test summary
chisq.test(M,simulate=TRUE,B=1000000)#use simulated p-values
fisher.test(M)
#if not significant = homogeneity = no heterogeneity among replicates = replicates are independant


#EXP4 (15 replicats) - females exposed to DS males

#Test for heterogeneity among replicates using a Pearson's Chi-squared test 
#(or a Fisher's Exact test if the expected frequencies were close to 5 or less) before performing the analyses on the pooled data set

M <- as.table(rbind(c(1,3,1), c(0,2,3),c(1,2,0),c(1,1,2),
                    c(1,0,2),c(1,0,3),c(1,2,2),c(1,3,0),
                    c(2,1,2),c(2,2,1),c(2,2,1),c(0,3,2),
                    c(0,2,3),c(2,1,1),c(1,2,2)))
dimnames(M) <- list(replicat = c("1", "2", "3","4", "5", "6","7", "8", "9","10","11", "12", "13","14", "15"),
                    phenotypeM = c("WS","IS", "DS"))

(Xsq <- chisq.test(M))  # Prints test summary
chisq.test(M,simulate=TRUE,B=1000000)#use simulated p-values
fisher.test(M)
#if not significant = homogeneity = no heterogeneity among replicates = replicates are independant


###########
2. Chi2 on Mating probability
###########
par(mfrow=c(2, 2))

#EXP1 - naive females
total <- as.table(rbind(cT<-c(22,21,21)))
dimnames(total) <- list(mating=c("#obs"),mated=c("WS","IS", "DS"))
(Xsq <- chisq.test(total))  # Prints test summary

total <- as.table(rbind(cT<-c(22,21,21)))
dimnames(total) <- list(mating=c("#obs"),mated=c("WS","IS", "DS"))
pi.theo<-c(110/295,128/295,57/295)#correction by no of courtship attempts
chisq.test(total,p=pi.theo)

#graph:
graphics.off()
pi<-total/margin.table(total)
barplot(pi,ylim=range(0,max(pi,pi.theo)+0.15),xlab= "Male phenotype", ylab="Mating proportions",main="Naive females",cex.lab=1.3,beside=TRUE,col=c("green","brown","grey"),space=c(0,0.2))
points(barplot(1:length(total),plot=FALSE),pi.theo, pch="-",cex=5)


##without correction for courtship
#2 by 2 comparisons and chi2 test with uncorrected p-value
total <- as.table(rbind(cT<-c(22,42)))#wet males
dimnames(total) <- list(mating=c("#obs"),mated=c("WS","IS&DS"))
pi.theo<-c(1/3,2/3)
chisq.test(total,p=pi.theo)

total <- as.table(rbind(cT<-c(43,21)))#intermediate males
dimnames(total) <- list(mating=c("#obs"),mated=c("WS&DS","IS"))
pi.theo<-c(2/3,1/3)
chisq.test(total,p=pi.theo)

total <- as.table(rbind(cT<-c(43,21)))#dry males
dimnames(total) <- list(mating=c("#obs"),mated=c("WS&IS","DS"))
pi.theo<-c(2/3,1/3)
chisq.test(total,p=pi.theo)

#2 by 2 comparisons and corrected P-value 
obs1 <- as.data.frame(matrix(rep(c("WS","IS", "DS"), times=c(22,21,21)), ncol=1)) 

names(obs1)<-c("Phenotype")
if(length(which(is.na(obs1) == TRUE))!=0){nombre.lig.na <- length(table(which(is.na(obs1) == TRUE, arr.ind = TRUE)[,1]))} else{nombre.lig.na<-0} 
cat("Number of lines with NA:", nombre.lig.na, "\n")
obs2<-na.omit(obs1) 
obs2[, 1]<-factor(obs2[, 1]) 
dataobs <- table(obs2, dnn = names(obs1))
addmargins(dataobs)
cat("Check that dataobs table corresponds to the table used to test multiple comparisons\n")
k<-length(dataobs )
n<-margin.table(dataobs )
pi.theo<-c(1/3,1/3,1/3)

classe<-NULL; p.value<-NULL
for(i in 1:k){classe<- c(classe, names(dataobs)[i]) ; obs<- data.frame(dataobs[i], n- dataobs[i]) ; 
obstheo<- chisq.test(obs, p=c(pi.theo[i], 1-pi.theo[i]))$expected ; n.cases.inf.5<-length(which(obstheo<5)) ; 
if(n.cases.inf.5 == 0){ p.value<-c(p.value, round(chisq.test(c(dataobs[i], n- dataobs[i]), p=c(pi.theo[i], (1- pi.theo[i])))$p.value, 9))} 
else {p.value<- c(p.value, NA)}}
p.mult<-data.frame(classe, p.value)
p.mult
p.mult2<-transform(p.mult, Holm=round(p.adjust(p.mult$p.value, method="holm"), 6))#p.value of WS x1, p.value of IS x2, p.value of DS x3
p.mult2

#p of Holm-Sidak
#1-(1-p)^(C-i+1)
#p=p.value
#C=number of tests (3)
#i=1st, 2nd, 3rd p.value
1-(1-0.9295680)^(3-3+1)
1-(1-0.9295680)^(3-2+1)
1-(1-0.8596838)^(3-1+1)

#p of Holm-Bonferroni
#THE ONE in p.mult2
#(C-i+1)*p
#p=p.value
#C=number of tests (3)
#i=1st, 2nd, 3rd p.value
(3-3+1)*0.9295680
(3-2+1)*0.9295680
(3-1+1)*0.8596838


##with correction for courtship
#2 by 2 comparisons and chi2 test with uncorrected p-value
total <- as.table(rbind(cT<-c(22,42)))#wet males
dimnames(total) <- list(mating=c("#obs"),mated=c("WS","IS&DS"))
pi.theo<-c(110/295,(128+57)/295)#correction by no of courtship attempts
chisq.test(total,p=pi.theo)

total <- as.table(rbind(cT<-c(43,21)))#intermediate males
dimnames(total) <- list(mating=c("#obs"),mated=c("WS&DS","IS"))
pi.theo<-c((110+57)/295,128/295)#correction by no of courtship attempts
chisq.test(total,p=pi.theo)

total <- as.table(rbind(cT<-c(43,21)))#dry males
dimnames(total) <- list(mating=c("#obs"),mated=c("WS&IS","DS"))
pi.theo<-c((110+128)/295,57/295)#correction by no of courtship attempts 
chisq.test(total,p=pi.theo)

#2 by 2 comparisons and corrected P-value 
obs1 <- as.data.frame(matrix(rep(c("WS","IS", "DS"), times=c(22,21,21)), ncol=1)) 

names(obs1)<-c("Phenotype")
if(length(which(is.na(obs1) == TRUE))!=0){nombre.lig.na <- length(table(which(is.na(obs1) == TRUE, arr.ind = TRUE)[,1]))} else{nombre.lig.na<-0} 
cat("Number of lines with NA:", nombre.lig.na, "\n")
obs2<-na.omit(obs1) 
obs2[, 1]<-factor(obs2[, 1]) 
dataobs <- table(obs2, dnn = names(obs1))
addmargins(dataobs)
cat("Check that dataobs table corresponds to the table used to test multiple comparisons\n")
k<-length(dataobs )
n<-margin.table(dataobs )
pi.theo<-c(110/295,128/295,57/295)#correction by no of courtship attempts 

classe<-NULL; p.value<-NULL
for(i in 1:k){classe<- c(classe, names(dataobs)[i]) ; obs<- data.frame(dataobs[i], n- dataobs[i]) ; 
obstheo<- chisq.test(obs, p=c(pi.theo[i], 1-pi.theo[i]))$expected ; n.cases.inf.5<-length(which(obstheo<5)) ; 
if(n.cases.inf.5 == 0){ p.value<-c(p.value, round(chisq.test(c(dataobs[i], n- dataobs[i]), p=c(pi.theo[i], (1- pi.theo[i])))$p.value, 9))} 
else {p.value<- c(p.value, NA)}}
p.mult<-data.frame(classe, p.value)
p.mult
p.mult2<-transform(p.mult, Holm=round(p.adjust(p.mult$p.value, method="holm"), 6))#p.value of WS x1, p.value of IS x2, p.value de DS x3
p.mult2

#p of Holm-Sidak
#1-(1-p)^(C-i+1)
#p=p.value
#C=number of tests (3)
#i=1st, 2nd, 3rd p.value
1-(1-0.629850523)^(3-3+1)
1-(1-0.087755339)^(3-2+1)
1-(1-0.006267234)^(3-1+1)

#p of Holm-Bonferroni
#THE ONE in p.mult2
#(C-i+1)*p
#p=p.value
#C=number of tests (3)
#i=1st, 2nd, 3rd p.value
(3-3+1)*0.629850523
(3-2+1)*0.087755339
(3-1+1)*0.006267234


#BETWEEN PHENOTYPE
#2 by 2 comparisons and corrected P-value with Fisher
#one technique is to realize the i contingency tables of all possible combinations,
#to perform Fisher's exact test on each of the tables and then correct the p values
#if the chi2 test conditions are met, then apply this test instead of Fisher's exact test
total <- as.table(rbind(cT<-c(22,21)))
dimnames(total) <- list(mating=c("#obs"),mated=c("WS","IS"))
(Xsq <- chisq.test(total))  # Prints test summary

total <- as.table(rbind(cT<-c(22,21)))
dimnames(total) <- list(mating=c("#obs"),mated=c("WS","IS"))
pi.theo<-c(110/238,128/238)#correction by no of courtship attempts 
chisq.test(total,p=pi.theo)


total <- as.table(rbind(cT<-c(22,21)))
dimnames(total) <- list(mating=c("#obs"),mated=c("WS","DS"))
(Xsq <- chisq.test(total))  # Prints test summary

total <- as.table(rbind(cT<-c(22,21)))
dimnames(total) <- list(mating=c("#obs"),mated=c("WS","DS"))
pi.theo<-c(110/167,57/167)#correction by no of courtship attempts 
chisq.test(total,p=pi.theo)


total <- as.table(rbind(cT<-c(21,21)))
dimnames(total) <- list(mating=c("#obs"),mated=c("IS","DS"))
(Xsq <- chisq.test(total))  # Prints test summary

total <- as.table(rbind(cT<-c(21,21)))
dimnames(total) <- list(mating=c("#obs"),mated=c("IS","DS"))
pi.theo<-c(128/185,57/185)#correction by no of courtship attempts 
chisq.test(total,p=pi.theo)

#Uncorrected by courtships
#p of Holm-Sidak
#1-(1-p)^(C-i+1)
#p=p.value
#C=number of tests (3)
#i=1st, 2nd, 3rd p.value
1-(1-1)^(3-3+1)
1-(1-0.8788)^(3-2+1)
1-(1-0.8788)^(3-1+1)

#p of Holm-Bonferroni
#THE ONE in p.mult2
#(C-i+1)*p
#p=p.value
#C=number of tests (3)
#i=1st, 2nd, 3rd p.value
(3-3+1)*1#ISvsDS 
(3-2+1)*0.8788#WSvsIS
(3-1+1)*0.8788#WSvsDS

#Corrected by courtships
#p of Holm-Sidak
#1-(1-p)^(C-i+1)
#p=p.value
#C=number of tests (3)
#i=1st, 2nd, 3rd p.value
1-(1-0.5155)^(3-3+1)
1-(1-0.04198)^(3-2+1)
1-(1-0.007071)^(3-1+1)

#p of Holm-Bonferroni
#THE ONE in p.mult2
#(C-i+1)*p
#p=p.value
#C=number of tests (3)
#i=1st, 2nd, 3rd p.value
(3-3+1)*0.5155#WSvsIS
(3-2+1)*0.04198#WSvsDS
(3-1+1)*0.007071#ISvsDS


#EXP2 - females exposed to WS males
total <- as.table(rbind(cT<-c(23,34,20)))
dimnames(total) <- list(mating=c("#obs"),mated=c("WS","IS", "DS"))
(Xsq <- chisq.test(total))  # Prints test summary

total <- as.table(rbind(cT<-c(23,34,20)))
dimnames(total) <- list(mating=c("#obs"),mated=c("WS","IS", "DS"))
pi.theo<-c(60/178,75/178,43/178)#correction by no of courtship attempts 
chisq.test(total,p=pi.theo)


#Graph
graphics.off()
pi<-total/margin.table(total)
barplot(pi,ylim=range(0,max(pi,pi.theo)+0.15),xlab= "Male phenotype", ylab="Mating proportions",main="Exposed to WS",cex.lab=1.3,beside=TRUE,col=c("green","brown","grey"),space=c(0,0.2))
points(barplot(1:length(total),plot=FALSE),pi.theo, pch="-",cex=5)


##with correction for courtship
#2 by 2 comparisons and chi2 test with uncorrected p-value
total <- as.table(rbind(cT<-c(23,54)))#wet males
dimnames(total) <- list(mating=c("#obs"),mated=c("WS","IS&DS"))
pi.theo<-c(1/3,2/3)
chisq.test(total,p=pi.theo)

total <- as.table(rbind(cT<-c(43,34)))#intermediate males
dimnames(total) <- list(mating=c("#obs"),mated=c("WS&DS","IS"))
pi.theo<-c(2/3,1/3)
chisq.test(total,p=pi.theo)

total <- as.table(rbind(cT<-c(57,20)))#dry males
dimnames(total) <- list(mating=c("#obs"),mated=c("WS&IS","DS"))
pi.theo<-c(2/3,1/3)
chisq.test(total,p=pi.theo)

#2 by 2 comparisons and corrected P-value 
obs1 <- as.data.frame(matrix(rep(c("WS","IS", "DS"), times=c(23,34,20)), ncol=1)) 

names(obs1)<-c("Phenotype")
if(length(which(is.na(obs1) == TRUE))!=0){nombre.lig.na <- length(table(which(is.na(obs1) == TRUE, arr.ind = TRUE)[,1]))} else{nombre.lig.na<-0} 
cat("Number of lines with NA:", nombre.lig.na, "\n")
obs2<-na.omit(obs1) 
obs2[, 1]<-factor(obs2[, 1]) 
dataobs <- table(obs2, dnn = names(obs1))
addmargins(dataobs)
cat("Check that dataobs table corresponds to the table used to test multiple comparisons\n")
k<-length(dataobs )
n<-margin.table(dataobs )
pi.theo<-c(1/3,1/3,1/3)

classe<-NULL; p.value<-NULL
for(i in 1:k){classe<- c(classe, names(dataobs)[i]) ; obs<- data.frame(dataobs[i], n- dataobs[i]) ; 
obstheo<- chisq.test(obs, p=c(pi.theo[i], 1-pi.theo[i]))$expected ; n.cases.inf.5<-length(which(obstheo<5)) ; 
if(n.cases.inf.5 == 0){ p.value<-c(p.value, round(chisq.test(c(dataobs[i], n- dataobs[i]), p=c(pi.theo[i], (1- pi.theo[i])))$p.value, 9))} 
else {p.value<- c(p.value, NA)}}
p.mult<-data.frame(classe, p.value)
p.mult
p.mult2<-transform(p.mult, Holm=round(p.adjust(p.mult$p.value, method="holm"), 6))
p.mult2

#p of Holm-Sidak
#1-(1-p)^(C-i+1)
#p=p.value
#C=number of tests (3)
#i=1st, 2nd, 3rd p.value
1-(1-0.51914860)^(3-3+1)
1-(1-0.17071842)^(3-2+1)
1-(1-0.04395104)^(3-1+1)

#p of Holm-Bonferroni
#THE ONE in p.mult2
#(C-i+1)*p
#p=p.value
#C=number of tests (3)
#i=1st, 2nd, 3rd p.value
(3-3+1)*0.51914860
(3-2+1)*0.17071842
(3-1+1)*0.04395104


##with correction for courtship
#2 by 2 comparisons and chi2 test with uncorrected p-value
total <- as.table(rbind(cT<-c(23,54)))#wet males
dimnames(total) <- list(mating=c("#obs"),mated=c("WS","IS&DS"))
pi.theo<-c(60/178,(75+43)/178)#correction by no of courtship attempts
chisq.test(total,p=pi.theo)

total <- as.table(rbind(cT<-c(43,34)))#intermediate males
dimnames(total) <- list(mating=c("#obs"),mated=c("WS&DS","IS"))
pi.theo<-c((60+43)/178,75/178)#correction by no of courtship attempts 
chisq.test(total,p=pi.theo)

total <- as.table(rbind(cT<-c(57,20)))#dry males
dimnames(total) <- list(mating=c("#obs"),mated=c("WS&IS","DS"))
pi.theo<-c((60+75)/178,43/178)#correction by no of courtship attempts 
chisq.test(total,p=pi.theo)

#2 by 2 comparisons and corrected P-value 
obs1 <- as.data.frame(matrix(rep(c("WS","IS", "DS"), times=c(23,34,20)), ncol=1)) 

names(obs1)<-c("Phenotype")
if(length(which(is.na(obs1) == TRUE))!=0){nombre.lig.na <- length(table(which(is.na(obs1) == TRUE, arr.ind = TRUE)[,1]))} else{nombre.lig.na<-0} 
cat("Number of lines with NA:", nombre.lig.na, "\n")
obs2<-na.omit(obs1) 
obs2[, 1]<-factor(obs2[, 1]) 
dataobs <- table(obs2, dnn = names(obs1))
addmargins(dataobs)
cat("Check that dataobs table corresponds to the table used to test multiple comparisons\n")
k<-length(dataobs )
n<-margin.table(dataobs )
pi.theo<-c(60/178,75/178,43/178)#correction by no of courtship attempts 

classe<-NULL; p.value<-NULL
for(i in 1:k){classe<- c(classe, names(dataobs)[i]) ; obs<- data.frame(dataobs[i], n- dataobs[i]) ; 
obstheo<- chisq.test(obs, p=c(pi.theo[i], 1-pi.theo[i]))$expected ; n.cases.inf.5<-length(which(obstheo<5)) ; 
if(n.cases.inf.5 == 0){ p.value<-c(p.value, round(chisq.test(c(dataobs[i], n- dataobs[i]), p=c(pi.theo[i], (1- pi.theo[i])))$p.value, 9))} 
else {p.value<- c(p.value, NA)}}
p.mult<-data.frame(classe, p.value)
p.mult
p.mult2<-transform(p.mult, Holm=round(p.adjust(p.mult$p.value, method="holm"), 6))
p.mult2

#p of Holm-Sidak
#1-(1-p)^(C-i+1)
#p=p.value
#C=number of tests (3)
#i=1st, 2nd, 3rd p.value
1-(1-0.7194771)^(3-3+1)
1-(1-0.7095675)^(3-2+1)
1-(1-0.4762173)^(3-1+1)

#p of Holm-Bonferroni
#THE ONE in p.mult2
#(C-i+1)*p
#p=p.value
#C=number of tests (3)
#i=1st, 2nd, 3rd p.value
(3-3+1)*0.7194771
(3-2+1)*0.7095675
(3-1+1)*0.4762173


#BETWEEN PHENOTYPE
#2 by 2 comparisons and corrected P-value with Fisher
#one technique is to realize the i contingency tables of all possible combinations,
#to perform Fisher's exact test on each of the tables and then correct the p values
#if the chi2 test conditions are met, then apply this test instead of Fisher's exact test
total <- as.table(rbind(cT<-c(23,34)))
dimnames(total) <- list(mating=c("#obs"),mated=c("WS","IS"))
(Xsq <- chisq.test(total))  # Prints test summary

total <- as.table(rbind(cT<-c(23,34)))
dimnames(total) <- list(mating=c("#obs"),mated=c("WS","IS"))
pi.theo<-c(60/135,75/135)#correction by no of courtship attempts 
chisq.test(total,p=pi.theo)


total <- as.table(rbind(cT<-c(23,20)))
dimnames(total) <- list(mating=c("#obs"),mated=c("WS","DS"))
(Xsq <- chisq.test(total))  # Prints test summary

total <- as.table(rbind(cT<-c(23,20)))
dimnames(total) <- list(mating=c("#obs"),mated=c("WS","DS"))
pi.theo<-c(60/103,43/103)#correction by no of courtship attempts 
chisq.test(total,p=pi.theo)


total <- as.table(rbind(cT<-c(34,20)))
dimnames(total) <- list(mating=c("#obs"),mated=c("IS","DS"))
(Xsq <- chisq.test(total))  # Prints test summary

total <- as.table(rbind(cT<-c(34,20)))
dimnames(total) <- list(mating=c("#obs"),mated=c("IS","DS"))
pi.theo<-c(75/118,43/118)#correction by no of courtship attempts 
chisq.test(total,p=pi.theo)

#Uncorrected by courtships
#p of Holm-Sidak
#1-(1-p)^(C-i+1)
#p=p.value
#C=number of tests (3)
#i=1st, 2nd, 3rd p.value
1-(1-0.6473)^(3-3+1)
1-(1-0.1451)^(3-2+1)
1-(1-0.05676)^(3-1+1)

#p of Holm-Bonferroni
#THE ONE in p.mult2
#(C-i+1)*p
#p=p.value
#C=number of tests (3)
#i=1st, 2nd, 3rd p.value
(3-3+1)*0.6473#WSvsDS 
(3-2+1)*0.1451#WSvsIS
(3-1+1)*0.05676#ISvsDS

#Corrected by courtships
#p of Holm-Sidak
#1-(1-p)^(C-i+1)
#p=p.value
#C=number of tests (3)
#i=1st, 2nd, 3rd p.value
1-(1-0.9274)^(3-3+1)
1-(1-0.534)^(3-2+1)
1-(1-0.5264)^(3-1+1)

#p of Holm-Bonferroni
#THE ONE in p.mult2
#(C-i+1)*p
#p=p.value
#C=number of tests (3)
#i=1st, 2nd, 3rd p.value
(3-3+1)*0.9274#ISvsDS
(3-2+1)*0.534#WSvsIS
(3-1+1)*0.5264#WSvsDS 


#EXP3 - females exposed to IS23 males
total <- as.table(rbind(cT<-c(18,18,27)))
dimnames(total) <- list(mating=c("#obs"),mated=c("WS","IS", "DS"))
(Xsq <- chisq.test(total))  # Prints test summary

total <- as.table(rbind(cT<-c(18,18,27)))
dimnames(total) <- list(mating=c("#obs"),mated=c("WS","IS", "DS"))
pi.theo<-c(20/42,2/42,20/42)#correction by no of courtship attempts 
chisq.test(total,p=pi.theo)

# Fisher test
observed <- c(18,18,27)
expected <- c((18*(20/42)),(18*(2/42)),(27*(20/42)))
MAT <- matrix(c(observed,expected), ncol=2)
fisher.test(MAT)#p.value=0.042

#Graph
graphics.off()
pi<-total/margin.table(total)
barplot(pi,ylim=range(0,max(pi,pi.theo)+0.15),xlab= "Male phenotype", ylab="Mating proportions",main="Exposed to IS",cex.lab=1.3,beside=TRUE,col=c("green","brown","grey"),space=c(0,0.2))
points(barplot(1:length(total),plot=FALSE),pi.theo, pch="-",cex=5)

##without correction for courtship
#2 by 2 comparisons and chi2 test with uncorrected p-value
total <- as.table(rbind(cT<-c(18,45)))#wet males
dimnames(total) <- list(mating=c("#obs"),mated=c("WS","IS&DS"))
pi.theo<-c(1/3,2/3)
chisq.test(total,p=pi.theo)

total <- as.table(rbind(cT<-c(45,18)))#intermediate males
dimnames(total) <- list(mating=c("#obs"),mated=c("WS&DS","IS"))
pi.theo<-c(2/3,1/3)
chisq.test(total,p=pi.theo)

total <- as.table(rbind(cT<-c(36,27)))#dry males
dimnames(total) <- list(mating=c("#obs"),mated=c("WS&IS","DS"))
pi.theo<-c(2/3,1/3)
chisq.test(total,p=pi.theo)


#2 by 2 comparisons and corrected P-value 
obs1 <- as.data.frame(matrix(rep(c("WS","IS", "DS"), times=c(18,18,27)), ncol=1)) 

names(obs1)<-c("Phenotype")
if(length(which(is.na(obs1) == TRUE))!=0){nombre.lig.na <- length(table(which(is.na(obs1) == TRUE, arr.ind = TRUE)[,1]))} else{nombre.lig.na<-0} 
cat("Number of lines with NA:", nombre.lig.na, "\n")
obs2<-na.omit(obs1) 
obs2[, 1]<-factor(obs2[, 1]) 
dataobs <- table(obs2, dnn = names(obs1))
addmargins(dataobs)
cat("Check that dataobs table corresponds to the table used to test multiple comparisons\n")
k<-length(dataobs )
n<-margin.table(dataobs )
pi.theo<-c(1/3,1/3,1/3)

classe<-NULL; p.value<-NULL
for(i in 1:k){classe<- c(classe, names(dataobs)[i]) ; obs<- data.frame(dataobs[i], n- dataobs[i]) ; 
obstheo<- chisq.test(obs, p=c(pi.theo[i], 1-pi.theo[i]))$expected ; n.cases.inf.5<-length(which(obstheo<5)) ; 
if(n.cases.inf.5 == 0){ p.value<-c(p.value, round(chisq.test(c(dataobs[i], n- dataobs[i]), p=c(pi.theo[i], (1- pi.theo[i])))$p.value, 9))} 
else {p.value<- c(p.value, NA)}}
p.mult<-data.frame(classe, p.value)
p.mult
p.mult2<-transform(p.mult, Holm=round(p.adjust(p.mult$p.value, method="holm"), 6))
p.mult2


#p of Holm-Sidak
#1-(1-p)^(C-i+1)
#p=p.value
#C=number of tests (3)
#i=1st, 2nd, 3rd p.value
1-(1-0.4226781)^(3-3+1)
1-(1-0.4226781)^(3-2+1)
1-(1-0.1088094)^(3-1+1)

#p of Holm-Bonferroni
#THE ONE in p.mult2
#(C-i+1)*p
#p=p.value
#C=number of tests (3)
#i=1st, 2nd, 3rd p.value
(3-3+1)*0.4226781#wet males
(3-2+1)*0.4226781#intermediate males
(3-1+1)*0.1088094#dry males


##with correction for courtship
#2 by 2 comparisons and chi2 test with uncorrected p-value
total <- as.table(rbind(cT<-c(18,45)))#wet males
dimnames(total) <- list(mating=c("#obs"),mated=c("WS","IS&DS"))
pi.theo<-c(20/42,(2+20)/42)#correction by no of courtship attempts 
chisq.test(total,p=pi.theo)

total <- as.table(rbind(cT<-c(45,18)))#intermediate males
dimnames(total) <- list(mating=c("#obs"),mated=c("WS&DS","IS"))
pi.theo<-c((20+20)/42,2/42)#correction by no of courtship attempts 
chisq.test(total,p=pi.theo)

observed <- c(45,18)#intermediate males
expected <- c((45*((20+20)/42)),(18*(2/42)))
MAT <- matrix(c(observed,expected), ncol=2)
fisher.test(MAT)

total <- as.table(rbind(cT<-c(36,27)))#dry males
dimnames(total) <- list(mating=c("#obs"),mated=c("WS&IS","DS"))
pi.theo<-c((20+2)/42,20/42)#correction by no of courtship attempts 
chisq.test(total,p=pi.theo)


#2 by 2 comparisons and corrected P-value 
obs1 <- as.data.frame(matrix(rep(c("WS","IS", "DS"), times=c(18,18,27)), ncol=1)) 

names(obs1)<-c("Phenotype")
if(length(which(is.na(obs1) == TRUE))!=0){nombre.lig.na <- length(table(which(is.na(obs1) == TRUE, arr.ind = TRUE)[,1]))} else{nombre.lig.na<-0} 
cat("Number of lines with NA:", nombre.lig.na, "\n")
obs2<-na.omit(obs1) 
obs2[, 1]<-factor(obs2[, 1]) 
dataobs <- table(obs2, dnn = names(obs1))
addmargins(dataobs)
cat("Check that dataobs table corresponds to the table used to test multiple comparisons\n")
k<-length(dataobs )
n<-margin.table(dataobs )
pi.theo<-c(20/42,2/42,20/42)#correction by no of courtship attempts 

classe<-NULL; p.value<-NULL
for(i in 1:k){classe<- c(classe, names(dataobs)[i]) ; obs<- data.frame(dataobs[i], n- dataobs[i]) ; 
obstheo<- chisq.test(obs, p=c(pi.theo[i], 1-pi.theo[i]))$expected ; n.cases.inf.5<-length(which(obstheo<5)) ; 
if(n.cases.inf.5 == 0){ p.value<-c(p.value, round(chisq.test(c(dataobs[i], n- dataobs[i]), p=c(pi.theo[i], (1- pi.theo[i])))$p.value, 9))} 
else {p.value<- c(p.value, NA)}}
p.mult<-data.frame(classe, p.value)
p.mult
p.mult2<-transform(p.mult, Holm=round(p.adjust(p.mult$p.value, method="holm"), 6))
p.mult2

#p of Holm-Bonferroni
#THE ONE in p.mult2
#(C-i+1)*p
#p=p.value
#C=number of tests (3)
#i=1st, 2nd, 3rd p.value
(3-3+1)*0.4492#dry males
(3-2+1)*0.002469#wet males
(3-1+1)*2.2e-16#intermediate males

#Fischer
#Corrected by courtships
#p of Holm-Bonferroni
#THE ONE in p.mult2
#(C-i+1)*p
#p=p.value
#C=number of tests (3)
#i=1st, 2nd, 3rd p.value
(3-1+1)*0.0002538#intermediate males

#BETWEEN PHENOTYPE
#2 by 2 comparisons and corrected P-value with Fisher
#one technique is to realize the i contingency tables of all possible combinations,
#to perform Fisher's exact test on each of the tables and then correct the p values
#if the chi2 test conditions are met, then apply this test instead of Fisher's exact test
total <- as.table(rbind(cT<-c(18,18)))
dimnames(total) <- list(mating=c("#obs"),mated=c("WS","IS"))
(Xsq <- chisq.test(total))  # Prints test summary

total <- as.table(rbind(cT<-c(18,18)))
dimnames(total) <- list(mating=c("#obs"),mated=c("WS","IS"))
pi.theo<-c(20/22,2/22)#correction by no of courtship attempts 
chisq.test(total,p=pi.theo)

observed <- c(18,18)
expected <- c((18*(20/22)),(18*(2/22)))
MAT <- matrix(c(observed,expected), ncol=2)
fisher.test(MAT)


total <- as.table(rbind(cT<-c(18,27)))
dimnames(total) <- list(mating=c("#obs"),mated=c("WS","DS"))
(Xsq <- chisq.test(total))  # Prints test summary

total <- as.table(rbind(cT<-c(18,27)))
dimnames(total) <- list(mating=c("#obs"),mated=c("WS","DS"))
pi.theo<-c(20/40,20/40)#correction by no of courtship attempts 
chisq.test(total,p=pi.theo)


total <- as.table(rbind(cT<-c(18,27)))
dimnames(total) <- list(mating=c("#obs"),mated=c("IS","DS"))
(Xsq <- chisq.test(total))  # Prints test summary

total <- as.table(rbind(cT<-c(18,27)))
dimnames(total) <- list(mating=c("#obs"),mated=c("IS","DS"))
pi.theo<-c(2/22,20/22)#correction by no of courtship attempts 
chisq.test(total,p=pi.theo)

observed <- c(18,27)
expected <- c((18*(2/22)),(27*(20/22)))
MAT <- matrix(c(observed,expected), ncol=2)
fisher.test(MAT)

#Uncorrected by courtships
#p of Holm-Bonferroni
#THE ONE in p.mult2
#(C-i+1)*p
#p=p.value
#C=number of tests (3)
#i=1st, 2nd, 3rd p.value
(3-3+1)*1#WSvsIS 
(3-2+1)*0.1797#WSvsDS
(3-1+1)*0.1797#ISvsDS


#Corrected by courtships
#p of Holm-Bonferroni
#THE ONE in p.mult2
#(C-i+1)*p
#p=p.value
#C=number of tests (3)
#i=1st, 2nd, 3rd p.value
(3-3+1)*0.1797#WSvsDS
(3-2+1)*5.494e-13#ISvsDS
(3-1+1)*2.2e-16#WSvsIS

#Fischer
#Corrected by courtships
#p of Holm-Bonferroni
#THE ONE in p.mult2
#(C-i+1)*p
#p=p.value
#C=number of tests (3)
#i=1st, 2nd, 3rd p.value
(3-3+1)*1#WSvsDS
(3-2+1)*0.006815#WSvsIS
(3-1+1)*0.002749#ISvsDS


#EXP4
total <- as.table(rbind(cT<-c(16,26,25)))
dimnames(total) <- list(mating=c("#obs"),mated=c("WS","IS", "DS"))
(Xsq <- chisq.test(total))  # Prints test summary

total <- as.table(rbind(cT<-c(16,26,25)))
dimnames(total) <- list(mating=c("#obs"),mated=c("WS","IS", "DS"))
pi.theo<-c(42/116,40/116,34/116)#correction by no of courtship attempts 
chisq.test(total,p=pi.theo)

#Graph
graphics.off()
pi<-total/margin.table(total)
barplot(pi,ylim=range(0,max(pi,pi.theo)+0.15),xlab= "Male phenotype", ylab="Mating proportions",main="Exposed to DS",cex.lab=1.3,beside=TRUE,col=c("green","brown","grey"),space=c(0,0.2))
points(barplot(1:length(total),plot=FALSE),pi.theo, pch="-",cex=5)

##without correction for courtship
#2 by 2 comparisons and chi2 test with uncorrected p-value
total <- as.table(rbind(cT<-c(16,51)))#wet males
dimnames(total) <- list(mating=c("#obs"),mated=c("WS","IS&DS"))
pi.theo<-c(1/3,2/3)
chisq.test(total,p=pi.theo)

total <- as.table(rbind(cT<-c(41,26)))#intermediate males
dimnames(total) <- list(mating=c("#obs"),mated=c("WS&DS","IS"))
pi.theo<-c(2/3,1/3)
chisq.test(total,p=pi.theo)

total <- as.table(rbind(cT<-c(42,25)))#dry males
dimnames(total) <- list(mating=c("#obs"),mated=c("WS&IS","DS"))
pi.theo<-c(2/3,1/3)
chisq.test(total,p=pi.theo)


#2 by 2 comparisons and corrected P-value 
obs1 <- as.data.frame(matrix(rep(c("WS","IS", "DS"), times=c(16,26,25)), ncol=1)) 

names(obs1)<-c("Phenotype")
if(length(which(is.na(obs1) == TRUE))!=0){nombre.lig.na <- length(table(which(is.na(obs1) == TRUE, arr.ind = TRUE)[,1]))} else{nombre.lig.na<-0} 
cat("Number of lines with NA:", nombre.lig.na, "\n")
obs2<-na.omit(obs1) 
obs2[, 1]<-factor(obs2[, 1]) 
dataobs <- table(obs2, dnn = names(obs1))
addmargins(dataobs)
cat("Check that dataobs table corresponds to the table used to test multiple comparisons\n")
k<-length(dataobs )
n<-margin.table(dataobs )
pi.theo<-c(1/3,1/3,1/3)

classe<-NULL; p.value<-NULL
for(i in 1:k){classe<- c(classe, names(dataobs)[i]) ; obs<- data.frame(dataobs[i], n- dataobs[i]) ; 
obstheo<- chisq.test(obs, p=c(pi.theo[i], 1-pi.theo[i]))$expected ; n.cases.inf.5<-length(which(obstheo<5)) ; 
if(n.cases.inf.5 == 0){ p.value<-c(p.value, round(chisq.test(c(dataobs[i], n- dataobs[i]), p=c(pi.theo[i], (1- pi.theo[i])))$p.value, 9))} 
else {p.value<- c(p.value, NA)}}
p.mult<-data.frame(classe, p.value)
p.mult
p.mult2<-transform(p.mult, Holm=round(p.adjust(p.mult$p.value, method="holm"), 6))
p.mult2

#p of Holm-Bonferroni
#THE ONE in p.mult2
#(C-i+1)*p
#p=p.value
#C=number of tests (3)
#i=1st, 2nd, 3rd p.value
(3-3+1)*0.4895060
(3-2+1)*0.3419826
(3-1+1)*0.1007248


##with correction for courtship
#2 by 2 comparisons and chi2 test with uncorrected p-value
total <- as.table(rbind(cT<-c(16,51)))#wet males
dimnames(total) <- list(mating=c("#obs"),mated=c("WS","IS&DS"))
pi.theo<-c(42/116,(40+34)/116)#correction by no of courtship attempts 
chisq.test(total,p=pi.theo)

total <- as.table(rbind(cT<-c(41,26)))#intermediate males
dimnames(total) <- list(mating=c("#obs"),mated=c("WS&DS","IS"))
pi.theo<-c((42+34)/116,40/116)#correction by no of courtship attempts 
chisq.test(total,p=pi.theo)

total <- as.table(rbind(cT<-c(42,25)))#dry males
dimnames(total) <- list(mating=c("#obs"),mated=c("WS&IS","DS"))
pi.theo<-c((42+40)/116,34/116)#correction by no of courtship attempts 
chisq.test(total,p=pi.theo)


#2 by 2 comparisons and corrected P-value 
obs1 <- as.data.frame(matrix(rep(c("WS","IS", "DS"), times=c(16,26,25)), ncol=1)) 

names(obs1)<-c("Phenotype")
if(length(which(is.na(obs1) == TRUE))!=0){nombre.lig.na <- length(table(which(is.na(obs1) == TRUE, arr.ind = TRUE)[,1]))} else{nombre.lig.na<-0} 
cat("Number of lines with NA:", nombre.lig.na, "\n")
obs2<-na.omit(obs1) 
obs2[, 1]<-factor(obs2[, 1]) 
dataobs <- table(obs2, dnn = names(obs1))
addmargins(dataobs)
cat("Check that dataobs table corresponds to the table used to test multiple comparisons\n")
k<-length(dataobs )
n<-margin.table(dataobs )
pi.theo<-c(42/116,40/116,34/116)#correction by no of courtship attempts 

classe<-NULL; p.value<-NULL
for(i in 1:k){classe<- c(classe, names(dataobs)[i]) ; obs<- data.frame(dataobs[i], n- dataobs[i]) ; 
obstheo<- chisq.test(obs, p=c(pi.theo[i], 1-pi.theo[i]))$expected ; n.cases.inf.5<-length(which(obstheo<5)) ; 
if(n.cases.inf.5 == 0){ p.value<-c(p.value, round(chisq.test(c(dataobs[i], n- dataobs[i]), p=c(pi.theo[i], (1- pi.theo[i])))$p.value, 9))} 
else {p.value<- c(p.value, NA)}}
p.mult<-data.frame(classe, p.value)
p.mult
p.mult2<-transform(p.mult, Holm=round(p.adjust(p.mult$p.value, method="holm"), 6))
p.mult2

#p of Holm-Bonferroni
#THE ONE in p.mult2
#(C-i+1)*p
#p=p.value
#C=number of tests (3)
#i=1st, 2nd, 3rd p.value
(3-3+1)*0.45657364
(3-2+1)*0.15010752
(3-1+1)*0.03578489


#BETWEEN PHENOTYPE
#2 by 2 comparisons and corrected P-value with Fisher
#one technique is to realize the i contingency tables of all possible combinations,
#to perform Fisher's exact test on each of the tables and then correct the p values
#if the chi2 test conditions are met, then apply this test instead of Fisher's exact test
total <- as.table(rbind(cT<-c(16,26)))
dimnames(total) <- list(mating=c("#obs"),mated=c("WS","IS"))
(Xsq <- chisq.test(total))  # Prints test summary

total <- as.table(rbind(cT<-c(16,26)))
dimnames(total) <- list(mating=c("#obs"),mated=c("WS","IS"))
pi.theo<-c(42/82,40/82)#correction by no of courtship attempts 
chisq.test(total,p=pi.theo)


total <- as.table(rbind(cT<-c(16,25)))
dimnames(total) <- list(mating=c("#obs"),mated=c("WS","DS"))
(Xsq <- chisq.test(total))  # Prints test summary

total <- as.table(rbind(cT<-c(16,25)))
dimnames(total) <- list(mating=c("#obs"),mated=c("WS","DS"))
pi.theo<-c(42/76,34/76)#correction by no of courtship attempts 
chisq.test(total,p=pi.theo)


total <- as.table(rbind(cT<-c(26,25)))
dimnames(total) <- list(mating=c("#obs"),mated=c("IS","DS"))
(Xsq <- chisq.test(total))  # Prints test summary

total <- as.table(rbind(cT<-c(26,25)))
dimnames(total) <- list(mating=c("#obs"),mated=c("IS","DS"))
pi.theo<-c(40/74,34/74)#correction by no of courtship attempts 
chisq.test(total,p=pi.theo)

#Uncorrected by courtships
#p of Holm-Bonferroni
#THE ONE in p.mult2
#(C-i+1)*p
#p=p.value
#C=number of tests (3)
#i=1st, 2nd, 3rd p.value
(3-3+1)*0.8886#ISvsDS 
(3-2+1)*0.1599#WSvsDS
(3-1+1)*0.1228#WSvsIS

#Corrected by courtships
#p of Holm-Bonferroni
#THE ONE in p.mult2
#(C-i+1)*p
#p=p.value
#C=number of tests (3)
#i=1st, 2nd, 3rd p.value
(3-3+1)*0.6596#ISvsDS
(3-2+1)*0.08883#WSvsIS
(3-1+1)*0.03651#WSvsDS 


#GRAPH
#Figure. Male mating proportions as a function of male seasonal phenotype in the experiments with naive females (a) (20 replicates), 
#females exposed to wet season males WS (b) (20 replicates), exposed to intermediate season males IS (c) (17 replicates), and to dry season males DS (d) (15 replicates). 
#The bars show the observed mating proportions. The dashes show the expected mating proportions corrected by male courtship propensity. 
#The adjusted Holm P-values (and uncorrected P-values in brackets) denote differences between observed and expected proportions and in paired comparisons between phenotypes. *<0.5, **<0.01.
graphics.off()
par(mfrow=c(2,2))
#EXP1
total <- as.table(rbind(cT<-c(22,21,21)))
dimnames(total) <- list(mating=c("#obs"),mated=c("wet","intermediate","dry"))
pi.theo<-c(110/295,128/295,57/295)#correction by no of courtship attempts 
chisq.test(total,p=pi.theo)
pi<-total/margin.table(total)
barplot(pi,ylim=range(0,0.6),xlab= "", ylab="Male mating proportions",main="Naive females (20 replicates)",cex.lab=1.3,beside=TRUE,col=c("white","white","white"),space=c(0,0.2))
points(barplot(1:length(total),plot=FALSE),pi.theo, pch="-",cex=5)
text(x = 0.3, y = 0.55, label = "(a)",cex=1.5)
text(x = 3.1, y = 0.38, label = "* (**)",cex=1)#* for DS
text(x = 1.9, y = 0.55, label = "____________",cex=1.5)
text(x = 1.9, y = 0.57, label = "(*)",cex=1)#* WS differs from DS
text(x = 2.5, y = 0.47, label = "_____",cex=1.5)
text(x = 2.5, y = 0.49, label = "* (**)",cex=1)#* IS differs from DS
text(x = 0.7, y = 0.02, label = "22 / 110",cex=0.7)#no of mating (no of courtships) for WS
text(x = 1.9, y = 0.02, label = "21 / 128",cex=0.7)# no of mating (no of courtships) for IS
text(x = 3.1, y = 0.02, label = "21 / 57",cex=0.7)# no of mating (no of courtships) for DS

#EXP2
total <- as.table(rbind(cT<-c(23,34,20)))
dimnames(total) <- list(mating=c("#obs"),mated=c("wet","intermediate","dry"))
pi.theo<-c(60/178,75/178,43/178)#correction by no of courtship attempts 
chisq.test(total,p=pi.theo)
pi<-total/margin.table(total)
barplot(pi,ylim=range(0,0.6),xlab= "", ylab="",main="Exposed to wet season males (20 replicates)",cex.lab=1.3,beside=TRUE,col=c("white","white","white"),space=c(0,0.2))
points(barplot(1:length(total),plot=FALSE),pi.theo, pch="-",cex=5)
text(x = 0.3, y = 0.55, label = "(b)",cex=1.5)
text(x = 0.7, y = 0.02, label = "23 / 60",cex=0.7)#no of mating (no of courtships) for WS
text(x = 1.9, y = 0.02, label = "34 / 75",cex=0.7)# no of mating (no of courtships) for IS
text(x = 3.1, y = 0.02, label = "20 / 43",cex=0.7)# no of mating (no of courtships) for DS

#EXP3
total <- as.table(rbind(cT<-c(18,18,27)))
dimnames(total) <- list(mating=c("#obs"),mated=c("wet","intermediate","dry"))
pi.theo<-c(20/42,2/42,20/42)#correction by no of courtship attempts 
chisq.test(total,p=pi.theo)
# Fisher test
observed <- c(18,18,27)
expected <- c((18*(20/42)),(18*(2/42)),(27*(20/42)))
MAT <- matrix(c(observed,expected), ncol=2)
fisher.test(MAT)#p.value=0.042
pi<-total/margin.table(total)
barplot(pi,ylim=range(0,0.6),xlab= "Male seasonal phenotypes", ylab="Male mating proportions",main="Exposed to intermediate season males (17 replicates)",cex.lab=1.3,beside=TRUE,col=c("white","white","white"),space=c(0,0.2))
points(barplot(1:length(total),plot=FALSE),pi.theo, pch="-",cex=5)
text(x = 0.3, y = 0.55, label = "(c)",cex=1.5)
text(x = 0.7, y = 0.35, label = "** (**)",cex=1)#* for WS
text(x = 1.9, y = 0.35, label = "*** (***)",cex=1)# for IS 
text(x = 1.3, y = 0.55, label = "_____",cex=1.5)
text(x = 1.3, y = 0.57, label = "* (**)",cex=1)#* WS differs from IS
text(x = 2.5, y = 0.55, label = "_____",cex=1.5)
text(x = 2.5, y = 0.57, label = "** (**)",cex=1)#** IS differs from DS
text(x = 0.7, y = 0.02, label = "18 / 20",cex=0.7)#no of mating (no of courtships) for WS
text(x = 1.9, y = 0.02, label = "18 / 2",cex=0.7)# no of mating (no of courtships) for IS
text(x = 3.1, y = 0.02, label = "27 / 20",cex=0.7)# no of mating (no of courtships) for DS

#EXP4
total <- as.table(rbind(cT<-c(16,26,25)))
dimnames(total) <- list(mating=c("#obs"),mated=c("wet","intermediate","dry"))
pi.theo<-c(42/116,40/116,34/116)#correction by no of courtship attempts 
chisq.test(total,p=pi.theo)
pi<-total/margin.table(total)
barplot(pi,ylim=range(0,0.6),xlab= "Male seasonal phenotypes", ylab="",main="Exposed to dry season males (15 replicates)",cex.lab=1.3,beside=TRUE,col=c("white","white","white"),space=c(0,0.2))
points(barplot(1:length(total),plot=FALSE),pi.theo, pch="-",cex=5)
text(x = 0.3, y = 0.55, label = "(d)",cex=1.5)
text(x = 0.7, y = 0.3, label = "(*)",cex=1)#* for WS
text(x = 1.9, y = 0.55, label = "____________",cex=1.5)
text(x = 1.9, y = 0.57, label = "(*)",cex=1)#* WS differs from DS
text(x = 0.7, y = 0.02, label = "16 / 42",cex=0.7)#no of mating (no of courtships) for WS
text(x = 1.9, y = 0.02, label = "26 / 40",cex=0.7)# no of mating (no of courtships) for IS
text(x = 3.1, y = 0.02, label = "25 / 34",cex=0.7)# no of mating (no of courtships) for DS


#Conclusions
#Naive females prefer to mate with DS males than with IS males and to a lower extent than with WS males;
#females exposed to WS males mate at random = expostition to WS males erase the innate preference for DS males;
#females exposed to IS males prefer to mate with IS males than with the other 2 male phenotypes;
#females exposed to DS males tend to prefer to mate with DS males as compared to WS males, but not as compared to IS males = exposition to DS males dot not reinforce the innate preference for DS males, but rather decrease it as females exposed to DS males prefer DS and IS males equally, while naive females prefered DS to IS males.


###########
3. Chi2 on Mating probability per male form across experiments
###########
par(mfrow=c(2, 2))


#WS males: EXP1 (20 replicats) vs. EXP2 (20 replicats) 
total <- as.table(rbind(cT<-c(22,23)))
dimnames(total) <- list(mating=c("#obs"),mated=c("WSexp1","WSexp2"))
(Xsq <- chisq.test(total))  # Prints test summary

#2 by 2 comparisons and chi2 test
total <- as.table(rbind(cT<-c(22,23)))#wet males
dimnames(total) <- list(mating=c("#obs"),mated=c("WSexp1","WSexp2"))
pi.theo<-c(1/2,1/2)
chisq.test(total,p=pi.theo)

#graph:
graphics.off()
pi<-total/margin.table(total)
barplot(pi,ylim=range(0,max(pi,pi.theo)+0.15),xlab= "Experiment", ylab="Mating proportions",main="WS males",cex.lab=1.3,beside=TRUE,col=c("green","brown"),space=c(0,0.2))
points(barplot(1:length(total),plot=FALSE),pi.theo, pch="-",cex=5)


#IS males: EXP1 (20 replicats) vs. EXP2 (20 replicats) 
total <- as.table(rbind(cT<-c(21,34)))
dimnames(total) <- list(mating=c("#obs"),mated=c("ISexp1","ISexp2"))
(Xsq <- chisq.test(total))  # Prints test summary

#2 by 2 comparisons and chi2 test
total <- as.table(rbind(cT<-c(21,34)))#is males
dimnames(total) <- list(mating=c("#obs"),mated=c("ISexp1","ISexp2"))
pi.theo<-c(1/2,1/2)
chisq.test(total,p=pi.theo)

#graph:
graphics.off()
pi<-total/margin.table(total)
barplot(pi,ylim=range(0,max(pi,pi.theo)+0.15),xlab= "Experiment", ylab="Mating proportions",main="IS males",cex.lab=1.3,beside=TRUE,col=c("green","brown"),space=c(0,0.2))
points(barplot(1:length(total),plot=FALSE),pi.theo, pch="-",cex=5)


#DS males: EXP1 (20 replicats) vs. EXP2 (20 replicats) 
total <- as.table(rbind(cT<-c(21,20)))
dimnames(total) <- list(mating=c("#obs"),mated=c("DSexp1","DSexp2"))
(Xsq <- chisq.test(total))  # Prints test summary

#2 by 2 comparisons and chi2 test
total <- as.table(rbind(cT<-c(21,20)))#dry males
dimnames(total) <- list(mating=c("#obs"),mated=c("DSexp1","DSexp2"))
pi.theo<-c(1/2,1/2)
chisq.test(total,p=pi.theo)

#graph:
graphics.off()
pi<-total/margin.table(total)
barplot(pi,ylim=range(0,max(pi,pi.theo)+0.15),xlab= "Experiment", ylab="Mating proportions",main="DS males",cex.lab=1.3,beside=TRUE,col=c("green","brown"),space=c(0,0.2))
points(barplot(1:length(total),plot=FALSE),pi.theo, pch="-",cex=5)


#WS males: EXP1 (17 replicats par règle de 3) vs. EXP3 (17 replicats)
22*17/20#19
total <- as.table(rbind(cT<-c(19,18)))
dimnames(total) <- list(mating=c("#obs"),mated=c("WSexp1","WSexp3"))
(Xsq <- chisq.test(total))  # Prints test summary

#2 by 2 comparisons and chi2 test
total <- as.table(rbind(cT<-c(19,18)))#wet males
dimnames(total) <- list(mating=c("#obs"),mated=c("WSexp1","WSexp3"))
pi.theo<-c(1/2,1/2)
chisq.test(total,p=pi.theo)

#graph:
graphics.off()
pi<-total/margin.table(total)
barplot(pi,ylim=range(0,max(pi,pi.theo)+0.15),xlab= "Experiment", ylab="Mating proportions",main="WS males",cex.lab=1.3,beside=TRUE,col=c("green","brown"),space=c(0,0.2))
points(barplot(1:length(total),plot=FALSE),pi.theo, pch="-",cex=5)


#IS males: EXP1 (17 replicats par règle de 3) vs. EXP3 (17 replicats)
21*17/20#18
total <- as.table(rbind(cT<-c(18,18)))
dimnames(total) <- list(mating=c("#obs"),mated=c("ISexp1","ISexp3"))
(Xsq <- chisq.test(total))  # Prints test summary

#2 by 2 comparisons and chi2 test
total <- as.table(rbind(cT<-c(18,18)))#is males
dimnames(total) <- list(mating=c("#obs"),mated=c("ISexp1","ISexp3"))
pi.theo<-c(1/2,1/2)
chisq.test(total,p=pi.theo)

#graph:
graphics.off()
pi<-total/margin.table(total)
barplot(pi,ylim=range(0,max(pi,pi.theo)+0.15),xlab= "Experiment", ylab="Mating proportions",main="IS males",cex.lab=1.3,beside=TRUE,col=c("green","brown"),space=c(0,0.2))
points(barplot(1:length(total),plot=FALSE),pi.theo, pch="-",cex=5)


#DS males: EXP1 (17 replicats par règle de 3) vs. EXP3 (17 replicats)
21*17/20#18
total <- as.table(rbind(cT<-c(18,27)))
dimnames(total) <- list(mating=c("#obs"),mated=c("DSexp1","DSexp3"))
(Xsq <- chisq.test(total))  # Prints test summary

#2 by 2 comparisons and chi2 test
total <- as.table(rbind(cT<-c(18,27)))#dry males
dimnames(total) <- list(mating=c("#obs"),mated=c("DSexp1","DSexp3"))
pi.theo<-c(1/2,1/2)
chisq.test(total,p=pi.theo)

#graph:
graphics.off()
pi<-total/margin.table(total)
barplot(pi,ylim=range(0,max(pi,pi.theo)+0.15),xlab= "Experiment", ylab="Mating proportions",main="DS males",cex.lab=1.3,beside=TRUE,col=c("green","brown"),space=c(0,0.2))
points(barplot(1:length(total),plot=FALSE),pi.theo, pch="-",cex=5)


#WS males: EXP1 (15 replicats par règle de 3) vs. EXP4 (15 replicats) 
22*15/20#17
total <- as.table(rbind(cT<-c(17,16)))
dimnames(total) <- list(mating=c("#obs"),mated=c("WSexp1","WSexp4"))
(Xsq <- chisq.test(total))  # Prints test summary

#2 by 2 comparisons and chi2 test
total <- as.table(rbind(cT<-c(17,16)))#wet males
dimnames(total) <- list(mating=c("#obs"),mated=c("WSexp1","WSexp4"))
pi.theo<-c(1/2,1/2)
chisq.test(total,p=pi.theo)

#graph:
graphics.off()
pi<-total/margin.table(total)
barplot(pi,ylim=range(0,max(pi,pi.theo)+0.15),xlab= "Experiment", ylab="Mating proportions",main="WS males",cex.lab=1.3,beside=TRUE,col=c("green","brown"),space=c(0,0.2))
points(barplot(1:length(total),plot=FALSE),pi.theo, pch="-",cex=5)


#IS males: EXP1 (15 replicats par règle de 3) vs. EXP4 (15 replicats) 
21*15/20#16
total <- as.table(rbind(cT<-c(16,26)))
dimnames(total) <- list(mating=c("#obs"),mated=c("ISexp1","ISexp4"))
(Xsq <- chisq.test(total))  # Prints test summary

#2 by 2 comparisons and chi2 test
total <- as.table(rbind(cT<-c(16,26)))#is males
dimnames(total) <- list(mating=c("#obs"),mated=c("ISexp1","ISexp4"))
pi.theo<-c(1/2,1/2)
chisq.test(total,p=pi.theo)

#graph:
graphics.off()
pi<-total/margin.table(total)
barplot(pi,ylim=range(0,max(pi,pi.theo)+0.15),xlab= "Experiment", ylab="Mating proportions",main="IS males",cex.lab=1.3,beside=TRUE,col=c("green","brown"),space=c(0,0.2))
points(barplot(1:length(total),plot=FALSE),pi.theo, pch="-",cex=5)


#DS males: EXP1 (15 replicats par règle de 3) vs. EXP4 (15 replicats) 
21*15/20#16
total <- as.table(rbind(cT<-c(16,25)))
dimnames(total) <- list(mating=c("#obs"),mated=c("DSexp1","DSexp4"))
(Xsq <- chisq.test(total))  # Prints test summary

#2 by 2 comparisons and chi2 test
total <- as.table(rbind(cT<-c(16,25)))#dry males
dimnames(total) <- list(mating=c("#obs"),mated=c("DSexp1","DSexp4"))
pi.theo<-c(1/2,1/2)
chisq.test(total,p=pi.theo)

#graph:
graphics.off()
pi<-total/margin.table(total)
barplot(pi,ylim=range(0,max(pi,pi.theo)+0.15),xlab= "Experiment", ylab="Mating proportions",main="DS males",cex.lab=1.3,beside=TRUE,col=c("green","brown"),space=c(0,0.2))
points(barplot(1:length(total),plot=FALSE),pi.theo, pch="-",cex=5)


###########
4. Chi2 on Mating probability between exp1 and the other exps pairwise
###########
par(mfrow=c(2, 2))


#EXP1 - naive females as theoretic probablilities (20 replicats) vs. EXP2 (20 replicats) 
total <- as.table(rbind(cT<-c(23,34,20)))
dimnames(total) <- list(mating=c("#obs"),mated=c("WS","IS", "DS"))
(Xsq <- chisq.test(total))  # Prints test summary

total <- as.table(rbind(cT<-c(23,34,20)))
dimnames(total) <- list(mating=c("#obs"),mated=c("WS","IS", "DS"))
pi.theo<-c(22/64,21/64,21/64)#correction by no of courtship attempts
chisq.test(total,p=pi.theo)

#graph:
graphics.off()
pi<-total/margin.table(total)
barplot(pi,ylim=range(0,max(pi,pi.theo)+0.15),xlab= "Male phenotype", ylab="Mating proportions",main="EXp2 vs. Exp1",cex.lab=1.3,beside=TRUE,col=c("green","brown","grey"),space=c(0,0.2))
points(barplot(1:length(total),plot=FALSE),pi.theo, pch="-",cex=5)


#EXP1 - naive females as theoretic probablilities (20 replicats) vs. EXP3 (17 replicats) 
total <- as.table(rbind(cT<-c(18,18,27)))
dimnames(total) <- list(mating=c("#obs"),mated=c("WS","IS", "DS"))
(Xsq <- chisq.test(total))  # Prints test summary

total <- as.table(rbind(cT<-c(18,18,27)))
dimnames(total) <- list(mating=c("#obs"),mated=c("WS","IS", "DS"))
pi.theo<-c(22/64,21/64,21/64)#correction by no of courtship attempts
chisq.test(total,p=pi.theo)

#graph:
graphics.off()
pi<-total/margin.table(total)
barplot(pi,ylim=range(0,max(pi,pi.theo)+0.15),xlab= "Male phenotype", ylab="Mating proportions",main="EXp3 vs. Exp1",cex.lab=1.3,beside=TRUE,col=c("green","brown","grey"),space=c(0,0.2))
points(barplot(1:length(total),plot=FALSE),pi.theo, pch="-",cex=5)


#EXP1 - naive females as theoretic probablilities (17 replicats par règle de 3)  vs. EXP3 (17 replicats) 
total <- as.table(rbind(cT<-c(18,18,27)))
dimnames(total) <- list(mating=c("#obs"),mated=c("WS","IS", "DS"))
(Xsq <- chisq.test(total))  # Prints test summary

total <- as.table(rbind(cT<-c(18,18,27)))
dimnames(total) <- list(mating=c("#obs"),mated=c("WS","IS", "DS"))
pi.theo<-c(19/55,18/55,18/55)#correction by no of courtship attempts
chisq.test(total,p=pi.theo)

#graph:
graphics.off()
pi<-total/margin.table(total)
barplot(pi,ylim=range(0,max(pi,pi.theo)+0.15),xlab= "Male phenotype", ylab="Mating proportions",main="EXp3 vs. Exp1",cex.lab=1.3,beside=TRUE,col=c("green","brown","grey"),space=c(0,0.2))
points(barplot(1:length(total),plot=FALSE),pi.theo, pch="-",cex=5)


#EXP1 - naive females as theoretic probablilities (20 replicats) vs. EXP4 (15 replicats) 
total <- as.table(rbind(cT<-c(16,26,25)))
dimnames(total) <- list(mating=c("#obs"),mated=c("WS","IS", "DS"))
(Xsq <- chisq.test(total))  # Prints test summary

total <- as.table(rbind(cT<-c(16,26,25)))
dimnames(total) <- list(mating=c("#obs"),mated=c("WS","IS", "DS"))
pi.theo<-c(22/64,21/64,21/64)#correction by no of courtship attempts
chisq.test(total,p=pi.theo)

#graph:
graphics.off()
pi<-total/margin.table(total)
barplot(pi,ylim=range(0,max(pi,pi.theo)+0.15),xlab= "Male phenotype", ylab="Mating proportions",main="EXp4 vs. Exp1",cex.lab=1.3,beside=TRUE,col=c("green","brown","grey"),space=c(0,0.2))
points(barplot(1:length(total),plot=FALSE),pi.theo, pch="-",cex=5)


#EXP1 - naive females as theoretic probablilities (15 replicats par règle de 3)  vs. EXP4 (15 replicats) 
total <- as.table(rbind(cT<-c(16,26,25)))
dimnames(total) <- list(mating=c("#obs"),mated=c("WS","IS", "DS"))
(Xsq <- chisq.test(total))  # Prints test summary

total <- as.table(rbind(cT<-c(16,26,25)))
dimnames(total) <- list(mating=c("#obs"),mated=c("WS","IS", "DS"))
pi.theo<-c(17/49,16/49,16/49)#correction by no of courtship attempts
chisq.test(total,p=pi.theo)

#graph:
graphics.off()
pi<-total/margin.table(total)
barplot(pi,ylim=range(0,max(pi,pi.theo)+0.15),xlab= "Male phenotype", ylab="Mating proportions",main="EXp4 vs. Exp1",cex.lab=1.3,beside=TRUE,col=c("green","brown","grey"),space=c(0,0.2))
points(barplot(1:length(total),plot=FALSE),pi.theo, pch="-",cex=5)
```
