## Supplementary material for "Warmer temperatures result in maladaptive learning of sexual preferences": Datasets and R Scripts: Script S5 fertility longevity.R

```
#Dataset S3
setwd("C:\\Users\\mullerd\\Desktop\\Projet anynana\\Th?se\\exp?rience volet 2\\exp2b\\ms finalisation\\ms_learning_soumission1")
setwd("F:/20180711_PCtravail/travail/Postdoc UCL/ms/B anynana/Holveck Muller/data and analyses")
fitness <- read.table("DatasetS3.txt", dec=".",header=TRUE, sep="\t")
summary(fitness)
str(fitness)
names(fitness)

#Matechoiceallowed and "MatingcageDensity","MatingcageSexratio","MatingcageVolume" are 100% correlated => can't be together in model
cor.test(fitness$MatingcageDensity,fitness$MatingcageVolume)
cor.test(as.numeric(fitness$MatingcageSexratio),fitness$MatingcageVolume)
cor.test(as.numeric(fitness$Matechoiceallowed),fitness$MatingcageVolume)

#######
#Fertility
#######
#All mated females to assess fertilisation success
#the needed categories = without females or males mated twice
fitnessOK2=fitness[fitness$NOmating=="1" & fitness$PartnerMatingorder=="1",
                   c("ButterflyID","Treatment","Treatmentduration","Experience","Matingage","PartnerMatingage","PartnerMatingorder",
                     "PartnerID","PartnerTreatment","NOmating","Fertility","Matechoiceallowed","Experiment","Experimenter","Replicate","Feedingstatus")]
summary(fitnessOK2)
str(fitnessOK2)

fitnessOK2sansNA<-na.omit(fitnessOK2)
summary(fitnessOK2sansNA)
str(fitnessOK2sansNA)
plot(factor(fitnessOK2sansNA$Fertility)~factor(fitnessOK2sansNA$PartnerTreatment))
plot(fitnessOK2sansNA$Fertility~fitnessOK2sansNA$PartnerTreatment)
abline(lm(fitnessOK2sansNA$Fertility~fitnessOK2sansNA$PartnerTreatment))

#only naive females CAN'T choose... => Experience and Matechoiceallowed CAN NOT be together in the same model
#Feedingstatus = only no food
#Experimenter = DM & LC
#Experiment = independent, Experiment2 and Experiment3

#GLM
mod<-glm(Fertility~scale(PartnerTreatment,center=TRUE,scale=TRUE)*Experience+
           scale(Matingage,center=TRUE,scale=TRUE)+
           scale(PartnerMatingage,center=TRUE,scale=TRUE)+
           scale(Treatmentduration,center=T,scale=T),family=binomial,data=fitnessOK2sansNA,na.action = "na.fail")
summary(mod) #OVERDISPERSION (Residual deviance > residual d.f.)

#VIF
library(car)
vif(glm(formula = fitnessOK2sansNA$Fertility ~ scale(fitnessOK2sansNA$PartnerTreatment,center=TRUE,scale=TRUE)+
          fitnessOK2sansNA$Experience+
          scale(fitnessOK2sansNA$Matingage,center=TRUE,scale=TRUE)+
          scale(fitnessOK2sansNA$PartnerMatingage,center=TRUE,scale=TRUE)+
          scale(fitnessOK2sansNA$Treatmentduration,center=T,scale=T)),binomial)

#GLMM with individual-level variability (accounting for overdispersion) = adding an observation-level random effect will capture the overdispersion.
library(lme4)
m0<-glmer(Fertility~scale(PartnerTreatment,center=T,scale=T)*Experience+
            scale(Matingage,center=T,scale=T)+
            scale(PartnerMatingage,center=T,scale=T)+
            scale(Treatmentduration,center=T,scale=T) + (1|Experimenter/Experiment/Replicate),family=binomial, data=fitnessOK2sansNA,na.action = "na.fail")
summary(m0)
m0_Nelder<- update(m0, control = glmerControl(optimizer="Nelder_Mead"))
m0_bobyqa<- update(m0, control = glmerControl(optimizer="bobyqa"))
summary(m0_bobyqa)

#GLMER dispersion
library(blmeco) 
dispersion_glmer(m0_bobyqa) #it should be 1 and it shouldn't be over 1.4

fitnessOK2sansNA$obs <- 1:nrow(fitnessOK2sansNA)
m1<-glmer(Fertility~scale(PartnerTreatment,center=T,scale=T)*Experience+
            scale(Matingage,center=T,scale=T)+
            scale(PartnerMatingage,center=T,scale=T)+
            scale(Treatmentduration,center=T,scale=T)+ (1|Experimenter/Experiment/Replicate)+(1|fitnessOK2sansNA$obs),family=binomial, data=fitnessOK2sansNA,na.action = "na.fail")
summary(m1)
m1_Nelder<- update(m1, control = glmerControl(optimizer="Nelder_Mead"))
m1_bobyqa<- update(m1, control = glmerControl(optimizer="bobyqa"))
summary(m1_bobyqa)

#You can also specify the proper order when the factor is created.
fitnessOK2sansNA$EXPER<- factor( fitnessOK2sansNA$Experience,
                             levels = c("naive","ISlearn","Wslearn"))
m2<-glmer(Fertility~scale(PartnerTreatment,center=T,scale=T)*EXPER+
            scale(Matingage,center=T,scale=T)+
            scale(PartnerMatingage,center=T,scale=T)+
            scale(Treatmentduration,center=T,scale=T)+ (1|Experimenter/Experiment/Replicate)+(1|fitnessOK2sansNA$obs),family=binomial, data=fitnessOK2sansNA,na.action = "na.fail")
summary(m2)
m2_Nelder<- update(m2, control = glmerControl(optimizer="Nelder_Mead"))
m2_bobyqa<- update(m2, control = glmerControl(optimizer="bobyqa"))
summary(m2_bobyqa)

#information theory model selection
#AICc should be used instead AIC when sample size is small in comparison to the number of estimated
#parameters (Burnham & Anderson 2002 recommend its use when n/K < 40); n=sample size; K=no of parameters.
165/15

library(MuMIn)
mod3<-dredge(m2_bobyqa, rank = "AICc",
             beta = "sd")#coefficients standardized by SD
mod3
mod_average <- summary(model.avg(mod3, subset = delta < 2))
mod_average
#cc
#the best model is the intercept....


#Only 'naive' and 'exposed to WS males' mated females to assess fertilisation success
#the needed categories = without females or males mated twice
fitnessOK2=fitness[fitness$NOmating=="1" & fitness$PartnerMatingorder=="1" & fitness$Experience=="naive" | fitness$Experience=="WSlearn",
                   c("ButterflyID","Treatment","Treatmentduration","Experience","Matingage","PartnerMatingage","PartnerMatingorder",
                     "PartnerID","PartnerTreatment","NOmating","Fertility","Matechoiceallowed","Experiment","Experimenter","Replicate","Feedingstatus")]
summary(fitnessOK2)
str(fitnessOK2)

fitnessOK2sansNA<-na.omit(fitnessOK2)
summary(fitnessOK2sansNA)
str(fitnessOK2sansNA)
plot(factor(fitnessOK2sansNA$Fertility)~factor(fitnessOK2sansNA$PartnerTreatment))
plot(fitnessOK2sansNA$Fertility~fitnessOK2sansNA$PartnerTreatment)
abline(lm(fitnessOK2sansNA$Fertility~fitnessOK2sansNA$PartnerTreatment))

#only naive females CAN'T choose... => Experience and Matechoiceallowed CAN NOT be together in the same model
#Feedingstatus = only no food
#Experimenter = DM & LC
#Experiment = independent by DM, Experiment2 by LC
#Experimenter and Experiment are 100% correlated

#GLM
mod<-glm(Fertility~scale(PartnerTreatment,center=TRUE,scale=TRUE)*Experience+
           scale(Matingage,center=TRUE,scale=TRUE)+
           scale(PartnerMatingage,center=TRUE,scale=TRUE)+
           scale(Treatmentduration,center=T,scale=T),family=binomial,data=fitnessOK2sansNA,na.action = "na.fail")
summary(mod) #OVERDISPERSION (Residual deviance > residual d.f.)

#VIF
library(car)
vif(glm(formula = fitnessOK2sansNA$Fertility ~ scale(fitnessOK2sansNA$PartnerTreatment,center=TRUE,scale=TRUE)+
          fitnessOK2sansNA$Experience+
          scale(fitnessOK2sansNA$Matingage,center=TRUE,scale=TRUE)+
          scale(fitnessOK2sansNA$PartnerMatingage,center=TRUE,scale=TRUE)+
          scale(fitnessOK2sansNA$Treatmentduration,center=T,scale=T)),binomial)

#GLMM with individual-level variability (accounting for overdispersion) = adding an observation-level random effect will capture the overdispersion.
library(lme4)
m0<-glmer(Fertility~scale(PartnerTreatment,center=T,scale=T)*Experience+
            scale(Matingage,center=T,scale=T)+
            scale(PartnerMatingage,center=T,scale=T)+
            scale(Treatmentduration,center=T,scale=T) + (1|Experiment/Replicate),family=binomial, data=fitnessOK2sansNA,na.action = "na.fail")
summary(m0)
m0_Nelder<- update(m0, control = glmerControl(optimizer="Nelder_Mead"))
m0_bobyqa<- update(m0, control = glmerControl(optimizer="bobyqa"))
summary(m0_bobyqa)

#GLMER dispersion
library(blmeco) 
dispersion_glmer(m0_bobyqa) #it should be 1 and it shouldn't be over 1.4

fitnessOK2sansNA$obs <- 1:nrow(fitnessOK2sansNA)
m1<-glmer(Fertility~scale(PartnerTreatment,center=T,scale=T)*Experience+
            scale(Matingage,center=T,scale=T)+
            scale(PartnerMatingage,center=T,scale=T)+
            scale(Treatmentduration,center=T,scale=T)+ (1|Experiment/Replicate)+(1|fitnessOK2sansNA$obs),family=binomial, data=fitnessOK2sansNA,na.action = "na.fail")
summary(m1)
m1_Nelder<- update(m1, control = glmerControl(optimizer="Nelder_Mead"))
m1_bobyqa<- update(m1, control = glmerControl(optimizer="bobyqa"))
summary(m1_bobyqa)

#information theory model selection
#AICc should be used instead AIC when sample size is small in comparison to the number of estimated
#parameters (Burnham & Anderson 2002 recommend its use when n/K < 40); n=sample size; K=no of parameters.
154/14

library(MuMIn)
mod3<-dredge(m1_bobyqa, rank = "AICc",
             beta = "sd")#coefficients standardized by SD
mod3
mod_average <- summary(model.avg(mod3, subset = delta < 2))
mod_average
#CC
#interaction !


#######
#Ageatdeath
#######


#All mated females 
#the needed categories = without females or males mated twice
dataOK=fitness[fitness$NOmating=="1" & fitness$PartnerMatingorder=="1",
                   c("ButterflyID","Treatment","Treatmentduration","Experience","Matingage","PartnerMatingage","PartnerMatingorder",
                     "PartnerID","PartnerTreatment","NOmating","Ageatdeath","Matechoiceallowed","Experiment","Experimenter","Replicate","Feedingstatus")]
summary(dataOK)
str(dataOK)

dataOKsansNA<-na.omit(dataOK)
summary(dataOKsansNA)
str(dataOKsansNA)
plot(dataOKsansNA$Ageatdeath~factor(dataOKsansNA$PartnerTreatment))
plot(dataOKsansNA$Ageatdeath~dataOKsansNA$PartnerTreatment)
abline(lm(dataOKsansNA$Ageatdeath~dataOKsansNA$PartnerTreatment))
plot(dataOKsansNA$Ageatdeath~factor(dataOKsansNA$Experience))

#naive females CAN or CAN'T choose... => Experience and Matechoiceallowed CAN be together in the same model
#Feedingstatus = fed, starved
#Experimenter = DM & LC & AT
#Experiment = independent, Experiment2 and Experiment3 and Experiment 1

hist(dataOKsansNA$Ageatdeath)

#GLM
mod<-glm(Ageatdeath~scale(PartnerTreatment,center=T,scale=T)*Experience+
           scale(PartnerTreatment,center=T,scale=T)*Feedingstatus+
           Matechoiceallowed+
           scale(Matingage,center=T,scale=T)+
           scale(PartnerMatingage,center=T,scale=T)+
           scale(Treatmentduration,center=T,scale=T),family=poisson,data=dataOKsansNA,na.action = "na.fail")
summary(mod) #OVERDISPERSION (Residual deviance > residual d.f.)
mod2<-glm(Ageatdeath~scale(PartnerTreatment,center=T,scale=T)*Experience+
            scale(PartnerTreatment,center=T,scale=T)*Feedingstatus+
            Matechoiceallowed+
            scale(Matingage,center=T,scale=T)+
            scale(PartnerMatingage,center=T,scale=T)+
            scale(Treatmentduration,center=T,scale=T),family=quasipoisson,data=dataOKsansNA,na.action = "na.fail")
summary(mod2) #OVERDISPERSION (Residual deviance > residual d.f.)


#VIF
library(car)
vif(glm(formula = dataOKsansNA$Ageatdeath ~ scale(dataOKsansNA$PartnerTreatment,center=T,scale=T)+
          dataOKsansNA$Experience+
          dataOKsansNA$Feedingstatus+
          dataOKsansNA$Matechoiceallowed+
          scale(dataOKsansNA$Matingage,center=T,scale=T)+
          scale(dataOKsansNA$PartnerMatingage,center=T,scale=T)+
          scale(dataOKsansNA$Treatmentduration,center=T,scale=T)),poisson)


#GLMM with individual-level variability (accounting for overdispersion) = adding an observation-level random effect will capture the overdispersion.
library(lme4)
m0<-glmer(Ageatdeath~scale(PartnerTreatment,center=T,scale=T)*Experience+
            scale(PartnerTreatment,center=T,scale=T)*Feedingstatus+
            Matechoiceallowed+
            scale(Matingage,center=T,scale=T)+
            scale(PartnerMatingage,center=T,scale=T)+
            scale(Treatmentduration,center=T,scale=T) + (1|Experimenter/Experiment/Replicate),family=poisson, data=dataOKsansNA,na.action = "na.fail")
summary(m0)
m0_Nelder<- update(m0, control = glmerControl(optimizer="Nelder_Mead"))
m0_bobyqa<- update(m0, control = glmerControl(optimizer="bobyqa"))
summary(m0_bobyqa)

#GLMER dispersion
library(blmeco) 
dispersion_glmer(m0_bobyqa) #it should be 1 and it shouldn't be over 1.4

dataOKsansNA$obs <- 1:nrow(dataOKsansNA)
m1<-glmer(Ageatdeath~scale(PartnerTreatment,center=T,scale=T)*Experience+
            scale(PartnerTreatment,center=T,scale=T)*Feedingstatus+
            Matechoiceallowed+
            scale(Matingage,center=T,scale=T)+
            scale(PartnerMatingage,center=T,scale=T)+
            scale(Treatmentduration,center=T,scale=T)+ (1|Experimenter/Experiment/Replicate)+(1|dataOKsansNA$obs),family=poisson, data=dataOKsansNA,na.action = "na.fail")
summary(m1)
m1_Nelder<- update(m1, control = glmerControl(optimizer="Nelder_Mead"))
m1_bobyqa<- update(m1, control = glmerControl(optimizer="bobyqa"))
summary(m1_bobyqa)

#You can also specify the proper order when the factor is created.
dataOKsansNA$EXPER<- factor( dataOKsansNA$Experience,
                             levels = c("naive","ISlearn","WSlearn"))
m2<-glmer(Ageatdeath~scale(PartnerTreatment,center=T,scale=T)*EXPER+
            scale(PartnerTreatment,center=T,scale=T)*Feedingstatus+
            Matechoiceallowed+
            scale(Matingage,center=T,scale=T)+
            scale(PartnerMatingage,center=T,scale=T)+
            scale(Treatmentduration,center=T,scale=T)+ (1|Experimenter/Experiment/Replicate)+(1|dataOKsansNA$obs),family=poisson, data=dataOKsansNA,na.action = "na.fail")
summary(m2)
m2_Nelder<- update(m2, control = glmerControl(optimizer="Nelder_Mead"))
m2_bobyqa<- update(m2, control = glmerControl(optimizer="bobyqa"))
summary(m2_bobyqa)


#information theory model selection
#AICc should be used instead AIC when sample size is small in comparison to the number of estimated
#parameters (Burnham & Anderson 2002 recommend its use when n/K < 40); n=sample size; K=no of parameters.
222/17 

library(MuMIn)
mod3<-dredge(m1_bobyqa, rank = "AICc",
             beta = "sd")#coefficients standardized by SD
mod3<-dredge(m2_bobyqa, rank = "AICc",
             beta = "sd")#coefficients standardized by SD
mod3
mod_average <- summary(model.avg(mod3, subset = delta < 2))
mod_average

#Best model = effect of experience
#naive females survived longer than experienced females
#cost of learning?


#Only 'naive' and 'exposed to WS males' mated females
#the needed categories = without females or males mated twice
dataOK=fitness[fitness$NOmating=="1" & fitness$PartnerMatingorder=="1" & fitness$Experience=="naive" | fitness$Experience=="WSlearn",
               c("ButterflyID","Treatment","Treatmentduration","Experience","Matingage","PartnerMatingage","PartnerMatingorder",
                 "PartnerID","PartnerTreatment","NOmating","Ageatdeath","Matechoiceallowed","Experiment","Experimenter","Replicate","Feedingstatus")]
summary(dataOK)
str(dataOK)

dataOKsansNA<-na.omit(dataOK)
summary(dataOKsansNA)
str(dataOKsansNA)
plot(dataOKsansNA$Ageatdeath~factor(dataOKsansNA$PartnerTreatment))
plot(dataOKsansNA$Ageatdeath~dataOKsansNA$PartnerTreatment)
abline(lm(dataOKsansNA$Ageatdeath~dataOKsansNA$PartnerTreatment))
plot(dataOKsansNA$Ageatdeath~factor(dataOKsansNA$Experience))

#naive females CAN or CAN'T choose... => Experience and Matechoiceallowed CAN be together in the same model
#Feedingstatus = fed, starved
#Experimenter = DM & LC & AT
#Experiment = independent, Experiment2 and Experiment 1

hist(dataOKsansNA$Ageatdeath)

#GLM
mod<-glm(Ageatdeath~scale(PartnerTreatment,center=T,scale=T)*Experience+
           scale(PartnerTreatment,center=T,scale=T)*Feedingstatus+
           Matechoiceallowed+
           scale(Matingage,center=T,scale=T)+
           scale(PartnerMatingage,center=T,scale=T)+
           scale(Treatmentduration,center=T,scale=T),family=poisson,data=dataOKsansNA,na.action = "na.fail")
summary(mod) #OVERDISPERSION (Residual deviance > residual d.f.)
mod2<-glm(Ageatdeath~scale(PartnerTreatment,center=T,scale=T)*Experience+
            scale(PartnerTreatment,center=T,scale=T)*Feedingstatus+
            Matechoiceallowed+
            scale(Matingage,center=T,scale=T)+
            scale(PartnerMatingage,center=T,scale=T)+
            scale(Treatmentduration,center=T,scale=T),family=quasipoisson,data=dataOKsansNA,na.action = "na.fail")
summary(mod2) #OVERDISPERSION (Residual deviance > residual d.f.)


#VIF
library(car)
vif(glm(formula = dataOKsansNA$Ageatdeath ~ scale(dataOKsansNA$PartnerTreatment,center=T,scale=T)+
          dataOKsansNA$Experience+
          dataOKsansNA$Feedingstatus+
          dataOKsansNA$Matechoiceallowed+
          scale(dataOKsansNA$Matingage,center=T,scale=T)+
          scale(dataOKsansNA$PartnerMatingage,center=T,scale=T)+
          scale(dataOKsansNA$Treatmentduration,center=T,scale=T)),poisson)


#GLMM with individual-level variability (accounting for overdispersion) = adding an observation-level random effect will capture the overdispersion.
library(lme4)
m0<-glmer(Ageatdeath~scale(PartnerTreatment,center=T,scale=T)*Experience+
            scale(PartnerTreatment,center=T,scale=T)*Feedingstatus+
            Matechoiceallowed+
            scale(Matingage,center=T,scale=T)+
            scale(PartnerMatingage,center=T,scale=T)+
            scale(Treatmentduration,center=T,scale=T) + (1|Experimenter/Experiment/Replicate),family=poisson, data=dataOKsansNA,na.action = "na.fail")
summary(m0)
m0_Nelder<- update(m0, control = glmerControl(optimizer="Nelder_Mead"))
m0_bobyqa<- update(m0, control = glmerControl(optimizer="bobyqa"))
summary(m0_bobyqa)

#GLMER dispersion
library(blmeco) 
dispersion_glmer(m0_bobyqa) #it should be 1 and it shouldn't be over 1.4

dataOKsansNA$obs <- 1:nrow(dataOKsansNA)
m1<-glmer(Ageatdeath~scale(PartnerTreatment,center=T,scale=T)*Experience+
            scale(PartnerTreatment,center=T,scale=T)*Feedingstatus+
            Matechoiceallowed+
            scale(Matingage,center=T,scale=T)+
            scale(PartnerMatingage,center=T,scale=T)+
            scale(Treatmentduration,center=T,scale=T)+ (1|Experimenter/Experiment/Replicate)+(1|dataOKsansNA$obs),family=poisson, data=dataOKsansNA,na.action = "na.fail")
summary(m1)
m1_Nelder<- update(m1, control = glmerControl(optimizer="Nelder_Mead"))
m1_bobyqa<- update(m1, control = glmerControl(optimizer="bobyqa"))
summary(m1_bobyqa)


#information theory model selection
#AICc should be used instead AIC when sample size is small in comparison to the number of estimated
#parameters (Burnham & Anderson 2002 recommend its use when n/K < 40); n=sample size; K=no of parameters.
211/17 

library(MuMIn)
mod3<-dredge(m1_bobyqa, rank = "AICc",
             beta = "sd")#coefficients standardized by SD
mod3
mod_average <- summary(model.avg(mod3, subset = delta < 2))
mod_average

#Best model = effect of experience
#naive females survived longer than experienced females
#cost of learning?


#Only 'naive' to assess fertilisation success
#the needed categories = without females or males mated twice
dataOK=fitness[fitness$NOmating=="1" & fitness$PartnerMatingorder=="1" & fitness$Experience=="naive",
               c("ButterflyID","Treatment","Treatmentduration","Experience","Matingage","PartnerMatingage","PartnerMatingorder",
                 "PartnerID","PartnerTreatment","NOmating","Ageatdeath","Experiment","Experimenter","Replicate",
                 "Feedingstatus","MatingcageVolume","MatingcageDensity","Matechoiceallowed","MatingcageSexratio")]
summary(dataOK)
str(dataOK)

dataOKsansNA<-na.omit(dataOK)
summary(dataOKsansNA)
str(dataOKsansNA)
plot(dataOKsansNA$Ageatdeath~factor(dataOKsansNA$PartnerTreatment))
plot(dataOKsansNA$Ageatdeath~dataOKsansNA$PartnerTreatment)
abline(lm(dataOKsansNA$Ageatdeath~dataOKsansNA$PartnerTreatment))
plot(dataOKsansNA$Ageatdeath~factor(dataOKsansNA$Experience))


#Matechoiceallowed and "MatingcageDensity","MatingcageSexratio","MatingcageVolume" and "Feedingstatus" are 100% correlated => can't be together in model
cor.test(dataOKsansNA$MatingcageDensity,dataOKsansNA$MatingcageVolume)
cor.test(as.numeric(dataOKsansNA$MatingcageSexratio),dataOKsansNA$MatingcageVolume)
cor.test(as.numeric(dataOKsansNA$Matechoiceallowed),dataOKsansNA$MatingcageVolume)
cor.test(as.numeric(dataOKsansNA$Feedingstatus),dataOKsansNA$MatingcageVolume)

#Feedingstatus = both
#Experimenter = DM & AT
#Experiment = independent by DM, experiment 1 by AT
#Experimenter and Experiment are 100% correlated


hist(dataOKsansNA$Ageatdeath)

#GLM
mod<-glm(Ageatdeath~scale(PartnerTreatment,center=T,scale=T)*Feedingstatus+
           scale(Matingage,center=T,scale=T)+
           scale(PartnerMatingage,center=T,scale=T)+
           scale(Treatmentduration,center=T,scale=T),family=poisson,data=dataOKsansNA,na.action = "na.fail")
summary(mod) #OVERDISPERSION (Residual deviance > residual d.f.)
mod2<-glm(Ageatdeath~scale(PartnerTreatment,center=T,scale=T)*Feedingstatus+
            scale(Matingage,center=T,scale=T)+
            scale(PartnerMatingage,center=T,scale=T)+
            scale(Treatmentduration,center=T,scale=T),family=quasipoisson,data=dataOKsansNA,na.action = "na.fail")
summary(mod2) #OVERDISPERSION (Residual deviance > residual d.f.)


#VIF
library(car)
vif(glm(formula = dataOKsansNA$Ageatdeath ~ scale(dataOKsansNA$PartnerTreatment,center=T,scale=T)+
          dataOKsansNA$Feedingstatus+
          scale(dataOKsansNA$Matingage,center=T,scale=T)+
          scale(dataOKsansNA$PartnerMatingage,center=T,scale=T)+
          scale(dataOKsansNA$Treatmentduration,center=T,scale=T)),poisson)


#GLMM with individual-level variability (accounting for overdispersion) = adding an observation-level random effect will capture the overdispersion.
library(lme4)
m0<-glmer(Ageatdeath~scale(PartnerTreatment,center=T,scale=T)*Feedingstatus+
            scale(Matingage,center=T,scale=T)+
            scale(PartnerMatingage,center=T,scale=T)+
            scale(Treatmentduration,center=T,scale=T) + (1|Experiment/Replicate),family=poisson, data=dataOKsansNA,na.action = "na.fail")
summary(m0)
m0_Nelder<- update(m0, control = glmerControl(optimizer="Nelder_Mead"))
m0_bobyqa<- update(m0, control = glmerControl(optimizer="bobyqa"))
summary(m0_bobyqa)

#GLMER dispersion
library(blmeco) 
dispersion_glmer(m0_bobyqa) #it should be 1 and it shouldn't be over 1.4

dataOKsansNA$obs <- 1:nrow(dataOKsansNA)
m1<-glmer(Ageatdeath~scale(PartnerTreatment,center=T,scale=T)*Feedingstatus+
            scale(Matingage,center=T,scale=T)+
            scale(PartnerMatingage,center=T,scale=T)+
            scale(Treatmentduration,center=T,scale=T)+ (1|Experiment/Replicate)+(1|dataOKsansNA$obs),family=poisson, data=dataOKsansNA,na.action = "na.fail")
summary(m1)
m1_Nelder<- update(m1, control = glmerControl(optimizer="Nelder_Mead"))
m1_bobyqa<- update(m1, control = glmerControl(optimizer="bobyqa"))
summary(m1_bobyqa)


#information theory model selection
#AICc should be used instead AIC when sample size is small in comparison to the number of estimated
#parameters (Burnham & Anderson 2002 recommend its use when n/K < 40); n=sample size; K=no of parameters.
140/9 

library(MuMIn)
mod3<-dredge(m0_bobyqa, rank = "AICc",
             beta = "sd")#coefficients standardized by SD
mod3
mod_average <- summary(model.avg(mod3, subset = delta < 2))
mod_average

#no effect; best model = intercept


#######
#NOeggs
#######

#Only 'naive' to assess fertilisation success
#the needed categories = without females or males mated twice
dataOK=fitness[fitness$NOmating=="1" & fitness$PartnerMatingorder=="1" & fitness$Experience=="naive",
               c("ButterflyID","Treatment","Treatmentduration","Experience","Matingage","PartnerMatingage","PartnerMatingorder",
                 "PartnerID","PartnerTreatment","NOmating","NOeggs","Experiment","Experimenter","Replicate",
                 "Feedingstatus","MatingcageVolume","MatingcageDensity","Matechoiceallowed","MatingcageSexratio")]
summary(dataOK)
str(dataOK)

dataOKsansNA<-na.omit(dataOK)
summary(dataOKsansNA)
str(dataOKsansNA)
plot(dataOKsansNA$NOeggs~factor(dataOKsansNA$PartnerTreatment))
plot(dataOKsansNA$NOeggs~dataOKsansNA$PartnerTreatment)
abline(lm(dataOKsansNA$NOeggs~dataOKsansNA$PartnerTreatment))
plot(dataOKsansNA$NOeggs~factor(dataOKsansNA$Experience))

plot(dataOKsansNA$NOeggs~dataOKsansNA$Feedingstatus)
plot(dataOKsansNA$NOeggs~dataOKsansNA$MatingcageVolume)
plot(dataOKsansNA$NOeggs~dataOKsansNA$MatingcageDensity)
plot(dataOKsansNA$NOeggs~dataOKsansNA$MatingcageSexratio)
plot(dataOKsansNA$NOeggs~dataOKsansNA$Matechoiceallowed)

#Matechoiceallowed and "MatingcageDensity","MatingcageSexratio","MatingcageVolume" and "Feedingstatus" are 100% correlated => can't be together in model
cor.test(dataOKsansNA$MatingcageDensity,dataOKsansNA$MatingcageVolume)
cor.test(as.numeric(dataOKsansNA$MatingcageSexratio),dataOKsansNA$MatingcageVolume)
cor.test(as.numeric(dataOKsansNA$Matechoiceallowed),dataOKsansNA$MatingcageVolume)
cor.test(as.numeric(dataOKsansNA$Feedingstatus),dataOKsansNA$MatingcageVolume)

#Feedingstatus = both
#Experimenter = DM & AT
#Experiment = independent by DM, experiment 1 by AT
#Experimenter and Experiment are 100% correlated


hist(dataOKsansNA$NOeggs)

#GLM
mod<-glm(NOeggs~scale(PartnerTreatment,center=T,scale=T)*Feedingstatus+
           scale(Matingage,center=T,scale=T)+
           scale(PartnerMatingage,center=T,scale=T)+
           scale(Treatmentduration,center=T,scale=T),family=poisson,data=dataOKsansNA,na.action = "na.fail")
summary(mod) #OVERDISPERSION (Residual deviance > residual d.f.)
mod2<-glm(NOeggs~scale(PartnerTreatment,center=T,scale=T)*Feedingstatus+
            scale(Matingage,center=T,scale=T)+
            scale(PartnerMatingage,center=T,scale=T)+
            scale(Treatmentduration,center=T,scale=T),family=quasipoisson,data=dataOKsansNA,na.action = "na.fail")
summary(mod2) #OVERDISPERSION (Residual deviance > residual d.f.)


#VIF
library(car)
vif(glm(formula = dataOKsansNA$NOeggs ~ scale(dataOKsansNA$PartnerTreatment,center=T,scale=T)+
          dataOKsansNA$Feedingstatus+
          scale(dataOKsansNA$Matingage,center=T,scale=T)+
          scale(dataOKsansNA$PartnerMatingage,center=T,scale=T)+
          scale(dataOKsansNA$Treatmentduration,center=T,scale=T)),quasipoisson)


#GLMM with individual-level variability (accounting for overdispersion) = adding an observation-level random effect will capture the overdispersion.
library(lme4)
m0<-glmer(NOeggs~scale(PartnerTreatment,center=T,scale=T)*Feedingstatus+
            scale(Matingage,center=T,scale=T)+
            scale(PartnerMatingage,center=T,scale=T)+
            scale(Treatmentduration,center=T,scale=T) + (1|Experiment/Replicate),family=poisson, data=dataOKsansNA,na.action = "na.fail")
summary(m0)
m0_Nelder<- update(m0, control = glmerControl(optimizer="Nelder_Mead"))
m0_bobyqa<- update(m0, control = glmerControl(optimizer="bobyqa"))
summary(m0_bobyqa)

#GLMER dispersion
library(blmeco) 
dispersion_glmer(m0_bobyqa) #it should be 1 and it shouldn't be over 1.4

dataOKsansNA$obs <- 1:nrow(dataOKsansNA)
m1<-glmer(NOeggs~scale(PartnerTreatment,center=T,scale=T)*Feedingstatus+
            scale(Matingage,center=T,scale=T)+
            scale(PartnerMatingage,center=T,scale=T)+
            scale(Treatmentduration,center=T,scale=T)+ (1|Experiment/Replicate)+(1|dataOKsansNA$obs),family=poisson, data=dataOKsansNA,na.action = "na.fail")
summary(m1)
m1_Nelder<- update(m1, control = glmerControl(optimizer="Nelder_Mead"))
m1_bobyqa<- update(m1, control = glmerControl(optimizer="bobyqa"))
summary(m1_bobyqa)


#information theory model selection
#AICc should be used instead AIC when sample size is small in comparison to the number of estimated
#parameters (Burnham & Anderson 2002 recommend its use when n/K < 40); n=sample size; K=no of parameters.
140/9 

library(MuMIn)
mod3<-dredge(m1_bobyqa, rank = "AICc",
             beta = "sd")#coefficients standardized by SD
mod3
mod_average <- summary(model.avg(mod3, subset = delta < 2))
mod_average

#effect of food after mating


##########
#GRAPHS
##########

#GRAPH interaction for fertility
fitnessOK3=fitness[fitness$NOmating=="1" & fitness$PartnerMatingorder=="1" & fitness$Experience== "naive",
                   c("ButterflyID","Treatment","Experience","Matingage","PartnerMatingage","PartnerMatingorder","PartnerID","PartnerTreatment","NOmating","Fertility")]
summary(fitnessOK3)
str(fitnessOK3)

fitnessOK3sansNA<-na.omit(fitnessOK3)
summary(fitnessOK3sansNA)
str(fitnessOK3sansNA)
plot(factor(fitnessOK3sansNA$Fertility)~factor(fitnessOK3sansNA$PartnerTreatment))


fitnessOK4=fitness[fitness$NOmating=="1" & fitness$PartnerMatingorder=="1" & fitness$Experience== "WSlearn",
                   c("ButterflyID","Treatment","Experience","Matingage","PartnerMatingage","PartnerMatingorder","PartnerID","PartnerTreatment","NOmating","Fertility")]
summary(fitnessOK4)
str(fitnessOK4)

fitnessOK4sansNA<-na.omit(fitnessOK4)
summary(fitnessOK4sansNA)
str(fitnessOK4sansNA)

par(mar = c(5, 5, 3, 5))
plot(factor(fitnessOK4sansNA$Fertility)~factor(fitnessOK4sansNA$PartnerTreatment),xlab="Male mate seasonal phenotype",ylab="Fertilisation success")
axis(side = 4)
mtext("Fertilization success proportion", side = 4, line = 3)


fitnessOK5=fitness[fitness$NOmating=="1" & fitness$PartnerMatingorder=="1" & fitness$Experience== "ISlearn",
                   c("ButterflyID","Treatment","Experience","Matingage","PartnerMatingage","PartnerMatingorder","PartnerID","PartnerTreatment","NOmating","Fertility")]
summary(fitnessOK5)
str(fitnessOK5)

fitnessOK5sansNA<-na.omit(fitnessOK5)
summary(fitnessOK5sansNA)
str(fitnessOK5sansNA)

par(mar = c(5, 5, 3, 5))
plot(factor(fitnessOK5sansNA$Fertility)~factor(fitnessOK5sansNA$PartnerTreatment),xlab="Male mate seasonal phenotype",ylab="Fertilisation success")
axis(side = 4)
mtext("Fertilization success proportion", side = 4, line = 3)


#GRAPH for longevity

fitnessOK6=fitness[fitness$NOmating=="1" & fitness$PartnerMatingorder=="1" & fitness$Experience== "naive" | fitness$Experience== "WSlearn" ,
                   c("ButterflyID","Treatment","Experience","Matingage","PartnerMatingage","PartnerMatingorder","PartnerID","PartnerTreatment","NOmating","Ageatdeath")]
summary(fitnessOK6)
str(fitnessOK6)

fitnessOK6sansNA<-na.omit(fitnessOK6)
summary(fitnessOK6sansNA)
str(fitnessOK6sansNA)

par(mar = c(5, 5, 3, 5))
plot(fitnessOK6sansNA$Ageatdeath~factor(fitnessOK6sansNA$PartnerTreatment),xlab="Male mate seasonal phenotype",ylab="Longevity")
axis(side = 4)
mtext("Longevity", side = 4, line = 3)


fitnessOK7=fitness[fitness$NOmating=="1" & fitness$PartnerMatingorder=="1",
                   c("ButterflyID","Treatment","Experience","Matingage","PartnerMatingage","PartnerMatingorder","PartnerID","PartnerTreatment","NOmating","Ageatdeath")]
summary(fitnessOK7)
str(fitnessOK7)

fitnessOK7sansNA<-na.omit(fitnessOK7)
summary(fitnessOK7sansNA)
str(fitnessOK7sansNA)

par(mar = c(5, 5, 3, 5))
plot(fitnessOK7sansNA$Ageatdeath~factor(fitnessOK7sansNA$PartnerTreatment),xlab="Male mate seasonal phenotype",ylab="Longevity")
axis(side = 4)
mtext("Longevity", side = 4, line = 3)


fitnessOK8=fitness[fitness$NOmating=="1" & fitness$PartnerMatingorder=="1" & fitness$Experience== "ISlearn" ,
                   c("ButterflyID","Treatment","Experience","Matingage","PartnerMatingage","PartnerMatingorder","PartnerID","PartnerTreatment","NOmating","Ageatdeath")]
summary(fitnessOK8)
str(fitnessOK8)

fitnessOK8sansNA<-na.omit(fitnessOK8)
summary(fitnessOK8sansNA)
str(fitnessOK8sansNA)

par(mar = c(5, 5, 3, 5))
plot(fitnessOK8sansNA$Ageatdeath~factor(fitnessOK8sansNA$PartnerTreatment),xlab="Male mate seasonal phenotype",ylab="Longevity")
axis(side = 4)
mtext("Longevity", side = 4, line = 3)


#Graph for NOeggs
fitnessOK9=fitness[fitness$NOmating=="1" & fitness$PartnerMatingorder=="1" & fitness$Experience=="naive",
               c("ButterflyID","Treatment","Treatmentduration","Experience","Matingage","PartnerMatingage","PartnerMatingorder",
                 "PartnerID","PartnerTreatment","NOmating","NOeggs","Experiment","Experimenter","Replicate",
                 "Feedingstatus","MatingcageVolume","MatingcageDensity","Matechoiceallowed","MatingcageSexratio")]
summary(fitnessOK9)
str(fitnessOK9)
fitnessOK9sansNA<-na.omit(fitnessOK9)
summary(fitnessOK9sansNA)
str(fitnessOK9sansNA)


#order WS - IS23 - DS
library(ggplot2)

fitnessOK3sansNA$PF<-factor(fitnessOK3sansNA$PartnerTreatment, levels = c("27","23","17"))
tapply(fitnessOK3sansNA$PF,fitnessOK3sansNA$Fertility,length)
library(plyr)
library(dplyr)
db<-ddply(fitnessOK3sansNA,.(PF,factor(Fertility)), summarise,
          N      = length(Fertility),
          Moy = mean(Fertility),
          sd     = sd(Fertility),
          se     = sd(Fertility) / sqrt(length(Fertility)))
db

p1<-
  ggplot(fitnessOK3sansNA, aes(x=fitnessOK3sansNA$PF, fill=factor(fitnessOK3sansNA$Fertility))) +
  geom_bar(position = "fill")+
  theme(panel.background = element_blank(),axis.line.x = element_line(colour = "black",size=0.75), axis.line.y = element_line(colour = "black",size=0.75), axis.text=element_text(size=13),axis.title=element_text(size=14,face="bold"))+
  xlab("Male mate seasonal phenotype") + ylab("Fertilization success proportion")+
  theme(strip.text.x = element_text(size=14),strip.background = element_rect(color="grey70", fill="white", size=1.5, linetype="solid"))+
  theme(legend.position="none",panel.border = element_rect(linetype = "dashed", fill = NA, color='grey80'))+
  scale_fill_manual(values=c("grey90","darkgrey"))+
  scale_x_discrete(labels = c('27'='wet','23'='intermediate','17'='dry'))+
  annotate("text", x = 3, y=0.05, label = "13") + 
  annotate("text", x = 3, y=0.95, label = "7")+
  annotate("text", x = 2, y=0.05, label = "31")+
  annotate("text", x = 2, y=0.95, label = "3")+
  annotate("text", x = 1, y=0.05, label = "30")+
  annotate("text", x = 1, y=0.95, label = "3")+
  annotate("text", x = 0.6, y=1.05, label = "(a) Naive females",cex=5,hjust = 0)+
  annotate("text", x = 3.4, y=0.5, label = "For (a) and (b)",hjust = 1)+
  annotate("text", x = 3.4, y=0.45, label = "Male mate seasonal phenotype: p=0.023",hjust = 1)+
  annotate("text", x = 3.4, y=0.4, label = "Female experience x Male mate seasonal phenotype: p=0.018",hjust = 1)
p1


fitnessOK4sansNA$PF<-factor(fitnessOK4sansNA$PartnerTreatment, levels = c("27","23","17"))
tapply(fitnessOK4sansNA$PF,fitnessOK4sansNA$Fertility,length)
library(plyr)
db<-ddply(fitnessOK4sansNA,.(PF,factor(Fertility)), summarise,
          N      = length(Fertility),
          Moy = mean(Fertility),
          sd     = sd(Fertility),
          se     = sd(Fertility) / sqrt(length(Fertility)))
db

p2<-
  ggplot(fitnessOK4sansNA, aes(x=fitnessOK4sansNA$PF, fill=factor(fitnessOK4sansNA$Fertility))) +
  geom_bar(position = "fill")+
  theme(panel.background = element_blank(),axis.line.x = element_line(colour = "black",size=0.75), axis.line.y = element_line(colour = "black",size=0.75), axis.text=element_text(size=13),axis.title=element_text(size=14,face="bold"))+
  xlab("Male mate seasonal phenotype") + ylab("")+
  theme(strip.text.x = element_text(size=14),strip.background = element_rect(color="grey70", fill="white", size=1.5, linetype="solid"))+
  theme(legend.position="none",panel.border = element_rect(linetype = "dashed", fill = NA, color='grey80'))+
  scale_fill_manual(values=c("grey90","darkgrey"))+
  scale_x_discrete(labels = c('27'='wet','23'='intermediate','17'='dry'))+
  annotate("text", x = 3, y=0.05, label = "13") + 
  annotate("text", x = 3, y=0.95, label = "2")+
  annotate("text", x = 2, y=0.05, label = "21")+
  annotate("text", x = 2, y=0.95, label = "5")+
  annotate("text", x = 1, y=0.05, label = "10")+
  annotate("text", x = 1, y=0.95, label = "5")+
  annotate("text", x = 0.6, y=1.05, label = "(b) Females exposed to wet season males",cex=5,hjust = 0)
  
p2


fitnessOK5sansNA$PF<-factor(fitnessOK5sansNA$PartnerTreatment, levels = c("27","23","17"))
tapply(fitnessOK5sansNA$PF,fitnessOK5sansNA$Fertility,length)
library(plyr)
db<-ddply(fitnessOK5sansNA,.(PF,factor(Fertility)), summarise,
          N      = length(Fertility),
          Moy = mean(Fertility),
          sd     = sd(Fertility),
          se     = sd(Fertility) / sqrt(length(Fertility)))
db

p3<-
  ggplot(fitnessOK5sansNA, aes(x=fitnessOK5sansNA$PF, fill=factor(fitnessOK5sansNA$Fertility))) +
  geom_bar(position = "fill")+
  theme(panel.background = element_blank(),axis.line.x = element_line(colour = "black",size=0.75), axis.line.y = element_line(colour = "black",size=0.75), axis.text=element_text(size=13),axis.title=element_text(size=14,face="bold"))+
  xlab("Male mate seasonal phenotype") + ylab("Fertilization success proportion")+
  theme(strip.text.x = element_text(size=14),strip.background = element_rect(color="grey70", fill="white", size=1.5, linetype="solid"))+
  theme(legend.position="none",panel.border = element_rect(linetype = "dashed", fill = NA, color='grey80'))+
  scale_fill_manual(values=c("grey90","darkgrey"))+
  scale_x_discrete(labels = c('27'='wet','23'='intermediate','17'='dry'))+
  annotate("text", x = 3, y=0.05, label = "4") + 
  annotate("text", x = 2, y=0.05, label = "8")+
  annotate("text", x = 2, y=0.95, label = "1")+
  annotate("text", x = 1, y=0.05, label = "6")+
  annotate("text", x = 1, y=0.95, label = "3")+
  annotate("text", x = 0.6, y=1.05, label = "(c) Females exposed to intermediate season males",cex=5,hjust = 0)

p3


fitnessOK6sansNA$PF<-factor(fitnessOK6sansNA$PartnerTreatment, levels = c("27","23","17"))
p4<-
  ggplot(fitnessOK6sansNA, aes(x=fitnessOK6sansNA$PF, y=fitnessOK6sansNA$Ageatdeath,fill=fitnessOK6sansNA$Experience)) +
  geom_boxplot()+
  theme(panel.background = element_blank(),axis.line.x = element_line(colour = "black",size=0.75), axis.line.y = element_line(colour = "black",size=0.75), axis.text=element_text(size=13),axis.title=element_text(size=14,face="bold"))+
  xlab("Male mate seasonal phenotype") + ylab("Longevity (days)")+
  theme(strip.text.x = element_text(size=14),strip.background = element_rect(color="grey70", fill="white", size=1.5, linetype="solid"))+
  theme(legend.position="none",panel.border = element_rect(linetype = "dashed", fill = NA, color='grey80'))+
  scale_fill_manual(values=c("white","black"))+
  scale_x_discrete(labels = c('27'='wet','23'='intermediate','17'='dry'))+
  ylim(0, 35)+
  annotate("text", x = 3.2, y=0.05, label = "16") + 
  annotate("text", x = 2.8, y=0.05, label = "34")+
  annotate("text", x = 2.2, y=0.05, label = "29")+
  annotate("text", x = 1.8, y=0.05, label = "50") + 
  annotate("text", x = 1.2, y=0.05, label = "20")+
  annotate("text", x = 0.8, y=0.05, label = "51")+
  annotate("text", x = 0.6, y=35, label = "(c) Naive versus exposed to wet season males",cex=5,hjust = 0)+
  annotate("text", x = 3.4, y=30, label = "Female experience: p<0.001",hjust = 1)

p4


fitnessOK7sansNA$PF<-factor(fitnessOK7sansNA$PartnerTreatment, levels = c("27","23","17"))
fitnessOK7sansNA$EXPER<- factor(fitnessOK7sansNA$Experience,
                             levels = c("naive","WSlearn","ISlearn"))
p5<-
  ggplot(fitnessOK7sansNA, aes(x=fitnessOK7sansNA$PF, y=fitnessOK7sansNA$Ageatdeath,fill=fitnessOK7sansNA$EXPER)) +
  geom_boxplot()+
  theme(panel.background = element_blank(),axis.line.x = element_line(colour = "black",size=0.75), axis.line.y = element_line(colour = "black",size=0.75), axis.text=element_text(size=13),axis.title=element_text(size=14,face="bold"))+
  xlab("Male mate seasonal phenotype") + ylab("Longevity (days)")+
  theme(strip.text.x = element_text(size=14),strip.background = element_rect(color="grey70", fill="white", size=1.5, linetype="solid"))+
  theme(legend.position="none",panel.border = element_rect(linetype = "dashed", fill = NA, color='grey80'))+
  scale_fill_manual(values=c("white","black","grey30"))+
  scale_x_discrete(labels = c('27'='wet','23'='intermediate','17'='dry'))+
  ylim(0, 35)+
  annotate("text", x = 3.25, y=0.05, label = "4") + 
  annotate("text", x = 3, y=0.05, label = "16") + 
  annotate("text", x = 2.75, y=0.05, label = "34") + 
  annotate("text", x = 2.25, y=0.05, label = "9")+
  annotate("text", x = 2, y=0.05, label = "29")+
  annotate("text", x = 1.75, y=0.05, label = "50")+
  annotate("text", x = 1.25, y=0.05, label = "9")+
  annotate("text", x = 1, y=0.05, label = "20")+
  annotate("text", x = 0.75, y=0.05, label = "51")+
  annotate("text", x = 0.6, y=35, label = "(d) Naive vs. exposed to wet vs. exposed to intermediate",cex=5,hjust = 0)

p5


fitnessOK8sansNA$PF<-factor(fitnessOK8sansNA$PartnerTreatment, levels = c("27","23","17"))
p6<-
  ggplot(fitnessOK8sansNA, aes(x=fitnessOK8sansNA$PF, y=fitnessOK8sansNA$Ageatdeath,fill=fitnessOK8sansNA$Experience)) +
  geom_boxplot()+
  theme(panel.background = element_blank(),axis.line.x = element_line(colour = "black",size=0.75), axis.line.y = element_line(colour = "black",size=0.75), axis.text=element_text(size=13),axis.title=element_text(size=14,face="bold"))+
  xlab("Male mate seasonal phenotype") + ylab("Longevity (days)")+
  theme(strip.text.x = element_text(size=14),strip.background = element_rect(color="grey70", fill="white", size=1.5, linetype="solid"))+
  theme(legend.position="none",panel.border = element_rect(linetype = "dashed", fill = NA, color='grey80'))+
  scale_fill_manual(values=c("grey30"))+
  scale_x_discrete(labels = c('27'='wet','23'='intermediate','17'='dry'))+
  ylim(0, 35)+
  annotate("text", x = 3, y=0.05, label = "4") + 
  annotate("text", x = 2, y=0.05, label = "9")+
  annotate("text", x = 1, y=0.05, label = "9")+
  annotate("text", x = 0.6, y=35, label = "(b)",cex=5,hjust = 0)

p6


fitnessOK9sansNA$PF<-factor(fitnessOK9sansNA$PartnerTreatment, levels = c("27","23","17"))
p7<-
  ggplot(fitnessOK9sansNA, aes(x=fitnessOK9sansNA$PF, y=fitnessOK9sansNA$NOeggs,fill=fitnessOK9sansNA$Feedingstatus)) +
  geom_boxplot()+
  theme(panel.background = element_blank(),axis.line.x = element_line(colour = "black",size=0.75), axis.line.y = element_line(colour = "black",size=0.75), axis.text=element_text(size=13),axis.title=element_text(size=14,face="bold"))+
  xlab("Male mate seasonal phenotype") + ylab("Number of eggs")+
  theme(strip.text.x = element_text(size=14),strip.background = element_rect(color="grey70", fill="white", size=1.5, linetype="solid"))+
  theme(legend.position="none",panel.border = element_rect(linetype = "dashed", fill = NA, color='grey80'))+
  scale_fill_manual(values=c("white","black"))+ #Fed versus starved naive females
  scale_x_discrete(labels = c('27'='wet','23'='intermediate','17'='dry'))+
  ylim(0, 220)+
  annotate("text", x = 3.2, y=220, label = "14") + 
  annotate("text", x = 2.8, y=220, label = "20") + 
  annotate("text", x = 2.2, y=220, label = "16")+
  annotate("text", x = 1.8, y=220, label = "34")+
  annotate("text", x = 1.2, y=220, label = "18")+
  annotate("text", x = 0.8, y=220, label = "33")+
  annotate("text", x = 0.6, y=220, label = "",cex=5,hjust = 0)

p7


##ALL GRAPHS on 1 page

# Multiple plot function
#
# ggplot objects can be passed in ..., or to plotlist (as a list of ggplot objects)
# - cols:   Number of columns in layout
# - layout: A matrix specifying the layout. If present, 'cols' is ignored.
#
# If the layout is something like matrix(c(1,2,3,3), nrow=2, by.row=TRUE),
# then plot 1 will go in the upper left, 2 will go in the upper right, and
# 3 will go all the way across the bottom.
#
multiplot <- function(..., plotlist=NULL, file, cols=1, layout=NULL) {
  require(grid)
  
  # Make a list from the ... arguments and plotlist
  plots <- c(list(...), plotlist)
  
  numPlots = length(plots)
  
  # If layout is NULL, then use 'cols' to determine layout
  if (is.null(layout)) {
    # Make the panel
    # ncol: Number of columns of plots
    # nrow: Number of rows needed, calculated from # of cols
    layout <- matrix(seq(1, cols * ceiling(numPlots/cols)),
                     ncol = cols, nrow = ceiling(numPlots/cols))
  }
  
  if (numPlots==1) {
    print(plots[[1]])
    
  } else {
    # Set up the page
    grid.newpage()
    pushViewport(viewport(layout = grid.layout(nrow(layout), ncol(layout))))
    
    # Make each plot, in the correct location
    for (i in 1:numPlots) {
      # Get the i,j matrix positions of the regions that contain this subplot
      matchidx <- as.data.frame(which(layout == i, arr.ind = TRUE))
      
      print(plots[[i]], vp = viewport(layout.pos.row = matchidx$row,
                                      layout.pos.col = matchidx$col))
    }}}


dev.new(width=17, height=13)

multiplot(p1,p3,p2,p5, cols=2)

multiplot(p1,p4,p2, cols=2)
```
